# Prediction of allosteric sites and signalling: insights from benchmarking datasets

**DOI:** 10.1101/2021.08.16.456251

**Authors:** Nan Wu, Léonie Strömich, Sophia N. Yaliraki

## Abstract

Allostery is a pervasive mechanism which regulates the activity of proteins in living systems through binding of a molecule at a distant site from the orthosteric site of the protein. The universality of allosteric regulation complemented by the benefits of highly specific, potentially non-toxic and protein activity modulating allosteric drugs makes uncovering allosteric sites on proteins invaluable for drug discovery. However, there are few computational methods to effectively predict them. Bond-to-bond propensity analysis, a recently developed method, has successfully predicted allosteric sites for a diverse group of proteins with only the knowledge of the orthosteric sites and the corresponding ligands in 19 of 20 cases. The method is based on an energy-weighted atomistic protein graph and allows for computationally highly efficient analysis in atomistic detail. We here extended the analysis onto 432 structures of 146 proteins from two existing benchmarking datasets for allosteric proteins: ASBench and CASBench. We further refined the metrics to account for the cumulative effect of residues with high propensities and the crucial residues in a given site with two additional measures. The allosteric site is recovered for 95/113 proteins (99/118 structures) from ASBench and 32/33 proteins (304/314 structures) from CASBench, with the only *a priori* knowledge being the orthosteric site residues. Knowing the orthosteric ligands of the protein, the allosteric site is identified for 32/33 proteins (308/314 structures) from CASBench.

## 1 Introduction

Proteins are ubiquitous in all aspects of cellular life where they fulfil crucial functions, while their malfunction could result in disease states [1, 2]. By 2017, 70% of small molecule drugs on the market targeted four types of proteins, namely protein kinases, ion channels, rhodopsin-like G protein-coupled receptors (GPCRs), and nuclear hormone receptors [3]. Most current small molecule drugs modify or inhibit the action of a protein by directly binding to the primary active site (also known as the orthosteric site) of the protein. The main advantage of this drug type is the high affinity and generally high specificity towards the orthosteric site as proved by a large number of successful drugs on the market [4]. Despite such advantages, the configuration of orthosteric sites is similar for proteins performing related functions and a low selectivity leads to off-target toxicity [5]. For instance, orthosteric sites for adenosine triphosphate (ATP) binding in different kinases are similar, thus making the optimisation of selective kinase inhibitor challenging [6]. In addition, prolonged exposure to the drugs results in drug resistance, through either modifications of the drug molecules [7] or changes to the orthosteric sites [8, 9, 10, 11, 12]. Moreover, orthosteric drugs act as complete inhibitors or activators rather than modulators of proteins and hence their therapeutic effect may not be the most optimal [10].

Modulation of protein activity, achieved through binding of small molecules at the allosteric site, is termed allosterism [4]. These binding events result in conformational changes of the targeted proteins and affect the binding of natural substrates to orthosteric sites. Conformational modification can enhance or reduce the binding affinity of natural substrates at orthosteric sites and can, therefore, lead to a controlled upregulation and downregulation of protein activities which is difficult to achieve by orthosteric site binding [13]. Allosteric modulators therefore have a lower potential for adverse side effects. Once all the allosteric sites are fully occupied, the drug reaches saturation (a ceiling level) and there is no further pharmacological effect. This indicates that on-target safety can be guaranteed even with overdosing [14, 15]. Contributing to the low off-target effects of allosteric drugs is the low evolutionary pressure for allosteric sites to accommodate an endogenous substrate compared to the well-conserved orthosteric sites [16]. This would allow for highly selective drug targeting in closely related protein families by exploiting allosterism.

The two main challenges for using allostery in drug development are finding suitable allosteric sites in the first place and designing molecules which bind and exert modulation effects. The design of allosteric site binders could follow well-established approaches used to develop molecules that bind to orthosteric sites, such as high-throughput screening [17], structure-based drug design [18] and peptide phage display [19]. To achieve a high specificity as well as the intended modulation, it is indispensable to search for unique allosteric sites for the targeted protein. Therefore, efficient and effective methods for identifying putative allosteric sites are of great interest to guide the rational design of allosteric modulators and contribute to the field of drug discovery and development [20].

Experimental methods including tethering [21, 22], nuclear magnetic resonance (NMR) [23, 24] and traditional high-throughput screening followed by X-ray crystallography [25, 26] have successfully led to the discovery of a few novel allosteric sites. All of these methods involve screening of huge compound libraries which is laborious and time-consuming. To circumvent the challenges associated with the experimental methods, numerous computational methods have been developed to predict allosteric sites (reviewed in [27, 28]) with various degrees of success. The continuous growth of the Allosteric Database (ASD) which contains data of 1949 allosteric proteins, their binding sites and other relevant information [29, 30, 31] and the construction of benchmarking datasets for allosteric proteins, ASBench [32] and CASBench [33], have provided comprehensive resources in aiding the identification of allosteric sites with computational methods.

There are two general ways of approaching the problem of identifying putative allosteric sites computationally: (1) identifying allosteric sites without considering the communication with orthosteric sites and (2) uncovering the allosteric-communication pathways between orthosteric and allosteric sites [34]. Several studies have followed the first approach: Huang *et al.* developed Allosite to find allosteric sites based on topological and physicochemical characteristics of allosteric and non-allosteric sites using a support vector machine (SVM) classifier [35], while Chen *et al.* built a random forest model which utilised calculated descriptors of orthosteric, allosteric and regular sites (binding sites without any function) and their bound ligands to classify potential sites on a given protein and identify putative allosteric sites [36]. Similarly, not concentrating on cognate ligands, Fogha *et al.* performed computational analysis of the density and clustering of crystallisation additives which are used to stabilise proteins during the process of crystallisation [37]. These methods, although achieving some promising predictability for putative allosteric sites, focus merely on the potential binding pockets on the proteins and do not consider the effects of binding at these sites on the protein, which is the key concept of allostery. Therefore, these approaches alone are not sufficient to identify potential allosteric sites. Molecular dynamics (MD) simulations and normal mode analysis (NMA) of elastic network models (ENM) are widely used within the second approach of identifying allosteric signalling paths based on protein dynamics described by Newton’s equation of motion. MD simulations can be applied to model proteins at atomic resolution and aid the understanding of communication pathways in proteins [38, 39]. For example, Shukla *et al.* applied MD simulations to reveal the structures of intermediates of a non-receptor tyrosine kinase c-Src and analysed its activation pathways to discover inhibitory allosteric sites [40]. However, MD simulations require a vast amount of computational resources if applied at an atomistic level for large proteins [41] and conventional all-atom MD simulations are unable to access the timescales of ligand-binding processes of proteins [42]. To retain crucial characteristics of dynamics and alleviate high computational demands, ENM were introduced. Performing NMA of ENM on proteins can result in a good match to MD simulations [43, 44, 45]. Most available methods include NMA of ENM as the main component and use a perturbation approach to measure the response of the protein to ligand binding or unbinding [34], thereby predicting allosteric sites, such as PARS [46, 47]. The results obtained from NMA of ENM can be combined with machine learning for the identification of allosteric sites and have been applied in AlloPred [48] and AllositePro [49]. Guarnera and Berezovsky introduced a structure-based statistical mechanical model of allostery (SBSMMA) which differs from ENM [50] to predict allosteric sites [51]. Although both ENM and SBSMMA are successful in modelling proteins and require much less computational power than MD simulations, they have two inherent limitations – not providing atomistic details of the protein and not considering long-range interactions above a certain distance. ENM treats each residue as a mass and represents a protein as a network of masses connected by virtual strings if they are within a cutoff distance [52]. SBSMMA uses the coarse-grained representation of proteins based on C*α* harmonic models and the allosteric potential is calculated only if the distance between two C*α* atoms is less than 11 Å [50]. This means that proteins represented by these two models are coarse-grained at the residue level and as a result subtle changes in protein conformations cannot be captured.

Bond-to-bond propensity analysis was introduced recently to circumvent these limitations, mainly to retain atomistic detail and remain computationally efficient. It has been shown capable of predicting allosteric sites requiring only knowledge of orthosteric sites and ligands [53]. The method builds on the construction of an atomistic graph from a biomolecular structure with atoms described as nodes and bonds, whether covalent or noncovalent, as weighted edges. The resulting protein graph is analysed with an edge-to-edge transfer matrix *M* (Methods) and the effect of fluctuations of an edge on any other edge is calculated and represented by a propensity score. Therefore, this approach enables the measurement of long-range coupling between bonds which is crucial for allosteric signalling. This graph-theoretical model differs from all of the computational methods discussed above, except MD simulations, as it uses a fully atomistic representation of a protein which retains the physico-chemical details of a protein [54, 55]. Despite keeping the atomistic details of the protein structure, the method is computationally efficient: by employing advances in algorithmic matrix theory [56, 57], the computation time scales approximately linearly with respect to the number of edges, which makes the method applicable to large and multimeric proteins [58, 59] and high-throughput analysis in general. Furthermore, since there is no cutoff distance for interactions, both weak and long-range interactions within a protein can be captured by this model. Therefore, bond-to-bond propensity analysis presents a more cost-effective computational method to analyse proteins at the atomistic level and predict potential allosteric sites.

Bond-to-bond propensity analysis has successfully predicted 19 out 20 allosteric sites for a test set of 20 proteins [53] and showcased the allostery in aspartate carbamoyltransferase (ATCase) and the main protease of the severe acute respiratory syndrome coronavirus 2 (SARS-CoV-2) [58, 60]. It has also been built into an efficient web application, ProteinLens, for the study of allostery [61]. To further benchmark this methodology and provide comparable insights into its performance across as diverse proteins as possible, we apply it here to two recently developed large, encompassing datasets, ASBench and CASBench. ASBench contains 235 allosteric sites [32] and computational methods such as AlloPred [48], AllositePro [49] and SBSMMA [51] have made use of this dataset for method validation. However, it is important to note that some of these methods use only the chain of the protein that contain orthosteric and allosteric sites. This means they may potentially miss communication between the sites if the pathway involves multiple chains or the entire protein structure as seen in multimeric proteins. We show in this work that bond-to-bond propensity analysis achieves overall higher accuracy in the ASBench dataset. We further tested bond-to-bond propensities with a more recent dataset, CASBench, which contains 91 protein entries with multiple crystal structures [33]. We evaluated the allosteric site prediction performance of our method in these datasets based on the four statistical measures used in [53] and two new measures introduced in this work.

## 2 Results

### 2.1 Bond-to-bond propensity analysis on the ASBench database

Proteins with annotated orthosteric residues, allosteric residues and ligands were collected from the ASBench and ASD databases as described in Methods and resulted in 118 structures of 113 distinct allosteric proteins. Bond-to-bond propensity analysis utilises the orthosteric ligand as the perturbation source to mimic the ligand-binding event [53] and identify regions on the protein which are functionally coupled to the orthosteric site. However, as orthosteric ligands are not available in structures from the ASBench database, the orthosteric site residues were selected as the source instead. For each protein, quantile scores, both intrinsic (*p*_*b*, allosteric site_, *p*_*R*, allosteric site_) and absolute 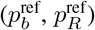, of all its bonds and residues were calculated with no *a priori* knowledge about the allosteric site (See Methods for more details). To assess the performance of the method and the significance of these calculated quantile scores, the allosteric site residues were used as the target point and evaluated with six statistical measures as described in Methods.

We here exemplify the method on bovine seminal ribonuclease (PDB ID: 11BG [62]), where we used the orthosteric site residues (Chain A: Asp14, Asn24, Asn27, Leu28, Asn94, Cys95, Chain B: Cys32 and Arg33) as the perturbation source. Figure 1 shows the propensity quantile score results mapped onto the protein structure where blue (0) indicates a low and red (1) a high connectivity to the active site. The values obtained from the statistical measures for the allosteric residues (allosteric ligand excluded if present) are summarised in Table 1.

**Figure 1:**
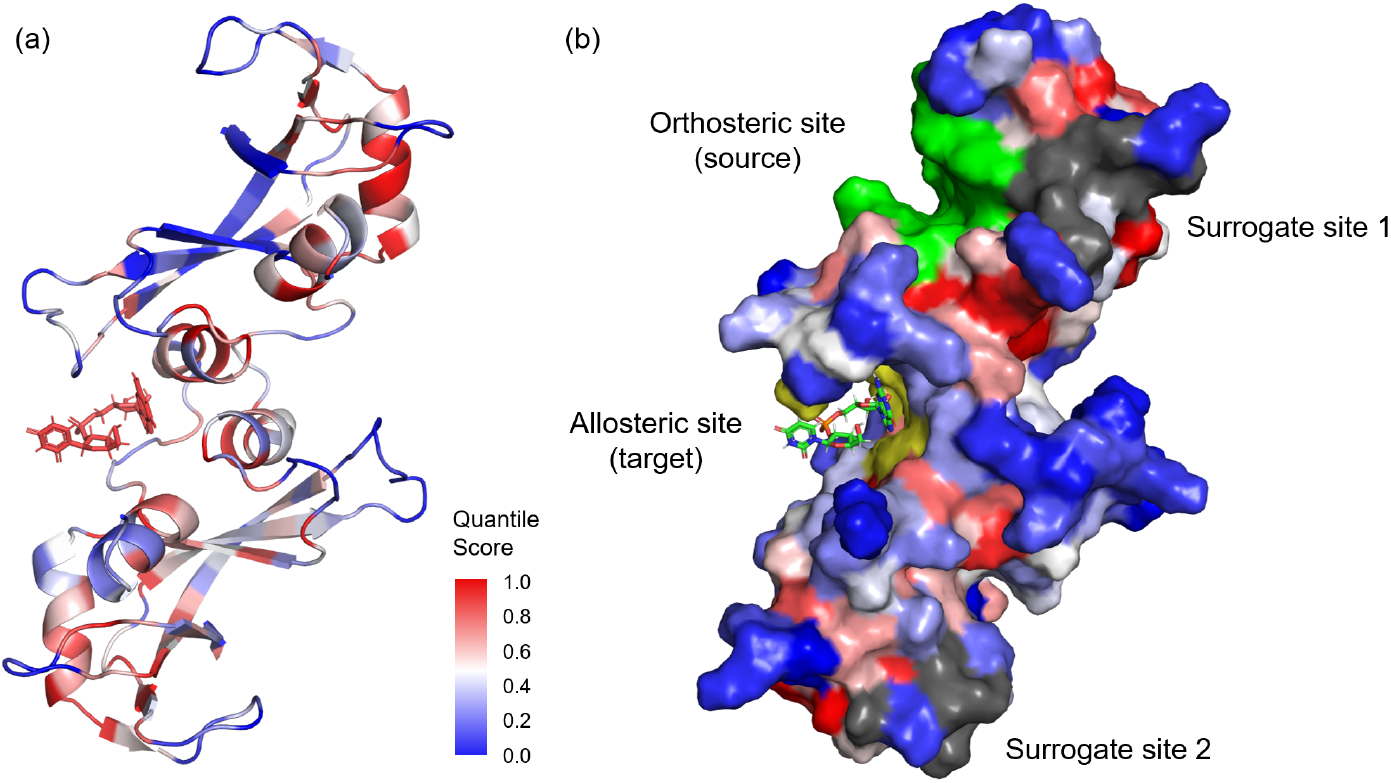
Bond-to-bond propensity analysis on the atomistic graph of bovine seminal ribonuclease (PDB ID: 11BG) where the orthosteric residues (green) are used as the perturbation source. **Note that:** (a) All residues are coloured by quantile score (QS) (see legend) obtained from bond-to-bond propensity analysis. (b) Surface representation of the protein structure coloured by QS. Relevant sites are highlighted and labelled accordingly.

**Table 1:** Results of bond-to-bond propensity analysis with six statistical measures for bovine seminal ribonuclease (11BG) (95% CI: 95% confidence interval)

| Statistical Measures | Results | Allosteric Site Detection |
| --- | --- | --- |
| $\overline{p_b}$, allosteric site<br>95% CI | 0.529 (> 0.495)<br>0.487, [0.478, 0.495] | Success |
| $\overline{p_R}$, allosteric site<br>95% CI | 0.665 (> 0.528)<br>0.525, [0.522, 0.528] | Success |
| $P(p_b, \text{allosteric site} > 0.95)$ | 0.081 (> 0.05) | Success |
| $P(p_R, \text{allosteric site} > 0.95)$ | 0.125 (> 0.05) | Success |
| $\overline{p_b^{\text{ref}}}$, allosteric site | 0.508 (> 0.5) | Success |
| $\overline{p_R^{\text{ref}}}$, allosteric site | 0.780 (> 0.5) | Success |

Based on the criteria described, the experimentally identified allosteric site can be detected with all six statistical measures. This process was conducted for all 118 proteins obtained from ASBench under two conditions – with and without the allosteric ligand in the structure. The results are shown in Fig 2.

**Figure 2:**
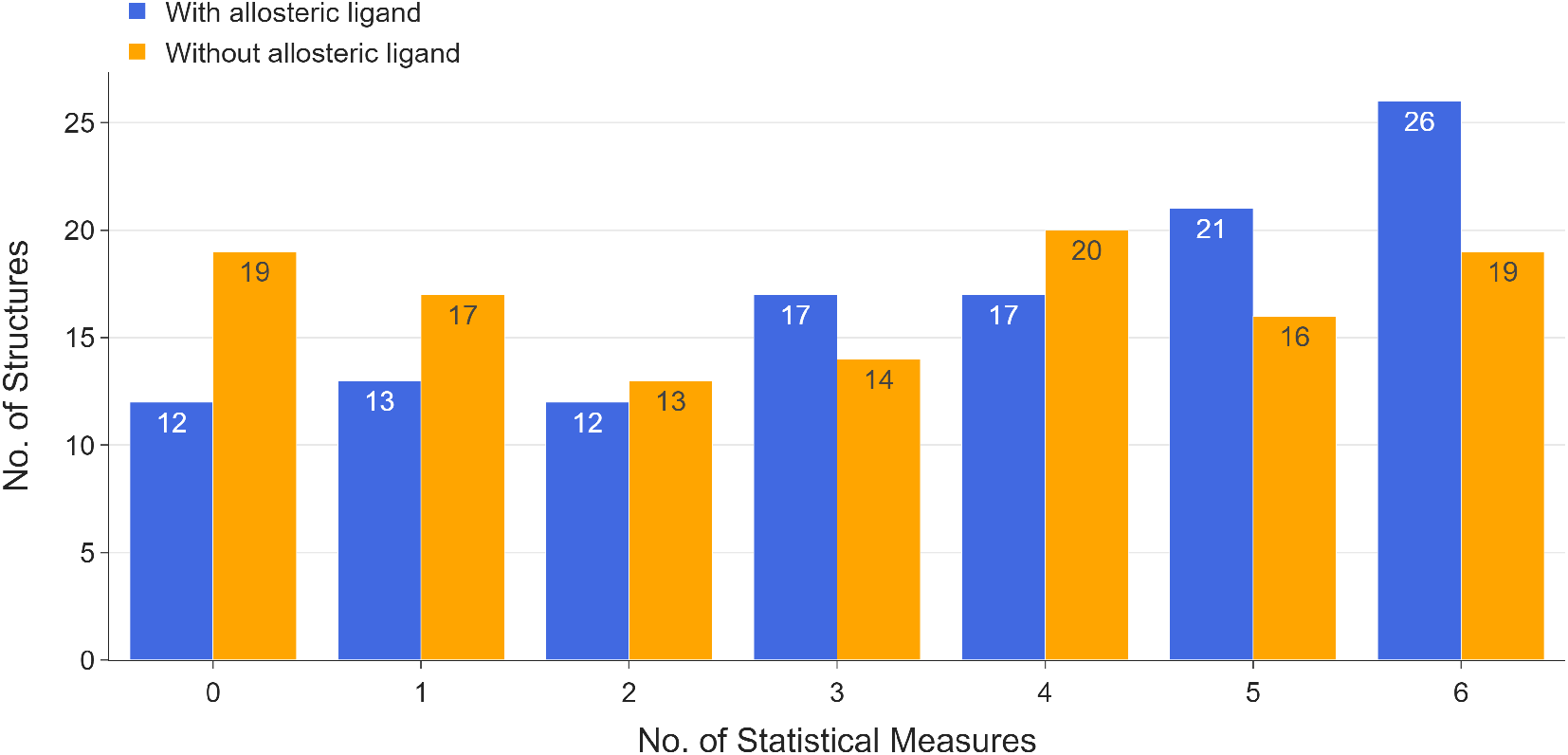
Allosteric site detection results for 118 structures in the ASBench database. The x-axis indicates the number of statistical measures for successful allosteric site detection.

In the presence of the allosteric ligand, the allosteric site is detected for 106/118 structures, according to at least one statistical measure, and for 81/118 structures, according to at least three statistical measures. When the allosteric ligand is removed from the protein structure and the same analysis is applied, the allosteric site is detected for 99/118 structures, according to at least one statistical measure, and for 69/118 structures, according to at least three statistical measures. The slight decrease in success rate is probably owing to the interaction of the allosteric ligand with the allosteric site residues. Since these allosteric ligands are effective allosteric modulators of the corresponding protein, the binding of the allosteric ligand would strengthen the functional coupling of the allosteric site to the orthosteric site which can be highlighted by the method. The average residue QS of the allosteric site for 109/118 structures decreases when the allosteric ligand is not present and those for the other nine structures only increased by less than 0.01 suggesting the same conclusion. Despite a lower success rate without the allosteric ligand, allosteric sites of 84% of the structures can be identified with only the knowledge of orthosteric site residues.

### 2.2 Prediction accuracy of bond-to-bond propensity analysis on the ASBench database

We focus here on the 12 structures with allosteric ligands where the allosteric site could not be detected by any of the measures. From those 12, the orthosteric residues of three structures (PDB IDs: 1UXV, 2VD3 and 3QH0) reported in the ASD database are incorrect (that is they do not form a binding site) and those of one further structure (PDB ID: 2ATS) do not match with the data in ASBench. From the remaining eight, six structures (PDB IDs: 1M8P, 3D2P, 3DC2, 3HQP, 3R1R and 4HYW) obtained from the ASBench are only one part of a large and complex multimeric protein, where the effect of cooperativity might play a crucial role. For example, it has been demonstrated with aspartate cabamoyltransferase (ATCase), a large dodecameric protein with six orthosteric sites, that only when at least three orthosteric sites are involved, allosteric behaviour is detected [58]. Since only one orthosteric site is reported in ASBench for these structures, this could explain the failure of identification of allosteric sites in these proteins when using only one orthosteric site as the perturbation source. From the remaining two structures, the G336V mutant of E.coli phosphoglycerate dehydrogenase (PDB ID: 2PA3) displays a different allosteric mechanism – the flip flop mechanism [63], which involves large scale mechanical changes. Lastly, the human muscle glycogen phosphorylase (PDB ID: 1Z8D) contains two allosteric sites [64] with only allosteric site 1 being detected, highlighted in red in Fig 3. This is due to the other site (highlighted in blue) being in close proximity to the orthosteric site where direct interactions, instead of long-range coupling, occur between the two sites.

**Figure 3:**
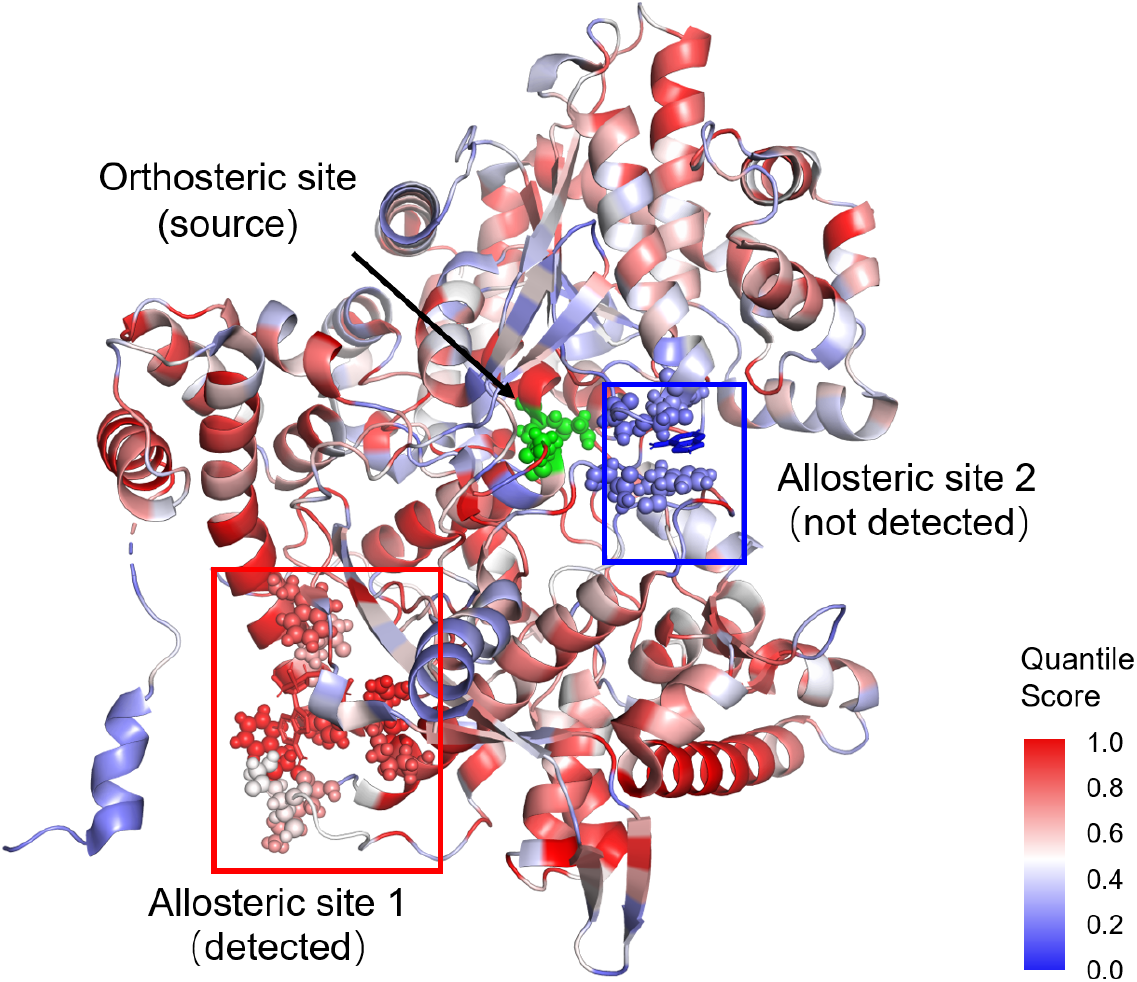
Structure of human muscle glycogen phosphorylase (PDB ID: 1Z8D [64]). The orthosteric (green) and two allosteric (cricled in blue and red) site residues are highlighted as spheres

Upon removing the allosteric ligands, allosteric sites of seven more structures could not be identified. For the structure of UDP-glucose dehydrogenase (PDB ID: 3PJG), ASBench has incorrect orthosteric residues reported (not forming a binding pocket) and hence, a wrong perturbation source was used. Haemoglobin (PDB ID: 1B86) is a well-known protein with cooperativity underpinning its activity [65] and contains four orthosteric sites. As only one orthosteric site is reported in ASBench, the coupling of the allosteric site to this one site could not be detected as it might not be strong enough. Two structures (PDB IDs: 3C1N and 3H6O) are large and complex multimeric proteins where again cooperativity would affect the results. The orthosteric sites and allosteric sites of the other three structures (PDB IDs: 2W4I, 3MWB, 4B1F), similar to those of 1Z8D above, are in close proximity. The allosteric effect is not mediated by long-range coupling and is thus not revealed by propensity analysis.

It is worth noting that the allosteric sites are generally large in size based on the definition provided in the ASBench database (residues within 6 Å from the allosteric ligand). In the previous bovine seminal ribonuclease (PDB ID: 11BG) example, the allosteric site contains eight residues but only four residues form direct interactions with the allosteric ligand. Defining the allosteric site using these four residues, which is essentially a sub-site of the original allosteric site, and rerunning all calculations give slightly different results as shown in Table 2.

**Table 2:**
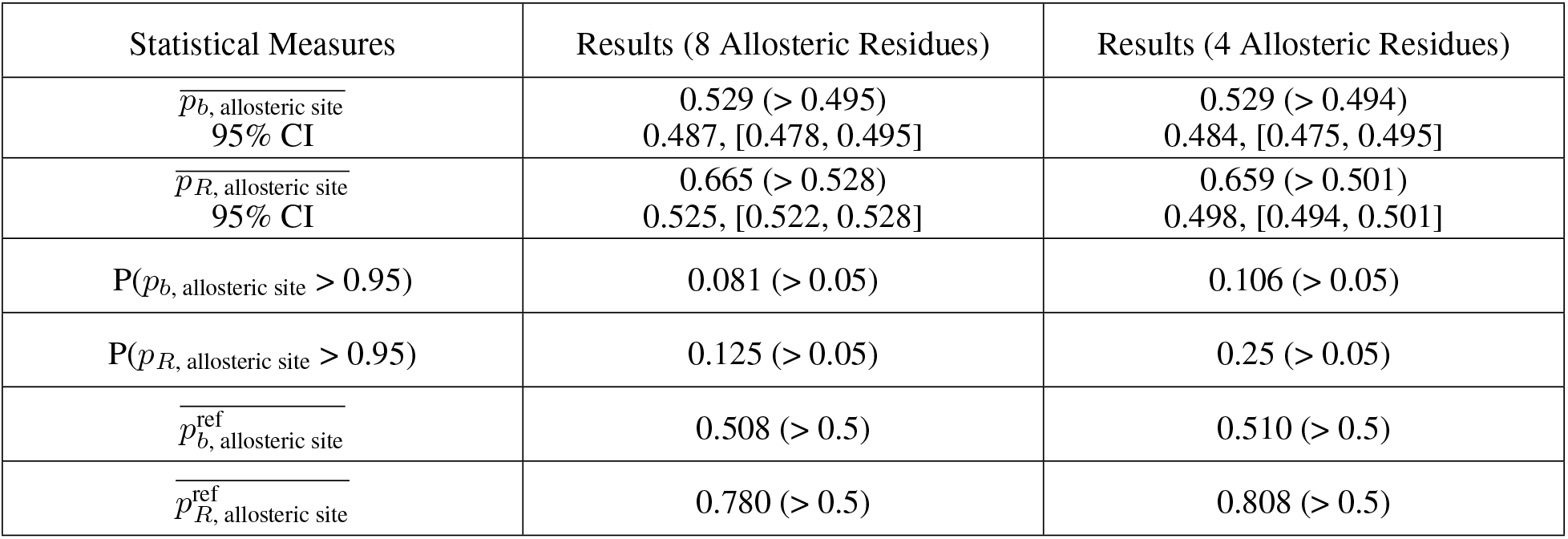
Results of bond-to-bond propensity analysis with six statistical measures for bovine seminal ribonuclease (PDB ID: 11BG) (95% CI: 95% confidence interval)

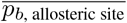 does not change while 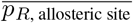 decreases slightly when only four allosteric residues were scored, however, comparisons with 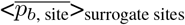 and 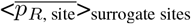 calculated from the 1,000 surrogate sites indicates that the allosteric site is more significant compared with other surrogate sites. The increase of values for the other four measures complements this argument. Therefore, defining the allosteric site with the four interacting residues leads to better detection of the allosteric site and one needs to take note that actual results may be buried by the definition of a large allosteric site. Hence, it is important to characterise the allosteric site and include relevant residues properly which presents an ongoing problem [66].

Similarly, not all residues in the orthosteric site defined in the database interact with the orthosteric ligand or support its binding. Due to the absence of orthosteric ligands in the structures from the ASBench database, comparisons between using the orthosteric site residues and the orthosteric ligand as perturbation source cannot be achieved.

### 2.3 Bond-to-bond propensity analysis on the CASBench database

314 structures of 33 allosteric proteins with orthosteric ligands and description of orthosteric and allosteric residues were collected from the CASBench database. As seen in the ASBench data analysis above, the presence of the allosteric ligand strengthens the coupling to the orthosteric site and makes the result biased towards successful detection of the allosteric site. Hence, the allosteric ligand (if present in the structure) is removed when carrying out bond-to-bond propensity analysis for the CASBench database.

Bond-to-bond propensity analysis was conducted for these 314 structures using the orthosteric ligand or orthosteric site residues (with orthosteric ligand removed) as the perturbation source in two separate runs. When multiple orthosteric ligands or sites are present, all of them were used as the source. Moreover, when there are multiple allosteric sites in the protein structure, each of them is investigated separately with the six statistical measures and the average value for each of the measures is used to decide whether the allosteric sites can be detected for the protein. Taking *Escherichia coli* biotin repressor (PDB ID: 2EWN [67]) as an example, which has two allosteric sites, the results are summarised in Table 3.

**Table 3:** Results of bond-to-bond propensity analysis with six statistical measures and averaging for *Escherichia coli* biotin repressor (PDB ID: 2EWN) (95% CI: 95% confidence interval). The two allosteric sites were scored separately based on the six metrics separately and the averaged scored was used to assess whether the allosteric sites of *Escherichia coli* biotin repressor can be detected by each measure.

| Statistical Measures | Results | Average | Allosteric Site Detection |
| --- | --- | --- | --- |
| $\overline{p_b}$, allosteric site<br>95% CI | Site 1: 0.333 (< 0.523)<br>0.516, [0.510, 0.523]<br>Site 2: 0.326 (< 0.528)<br>0.521, [0.515, 0.528] | 0.329 (< 0.523)<br>0.516, [0.510, 0.523] | Failure |
| $\overline{p_R}$, allosteric site<br>95% CI | Site 1: 0.532 (> 0.509)<br>0.506, [0.504, 0.509]<br>Site 2: 0.500 (< 0.510)<br>0.507, [0.504, 0.510] | 0.516 (< 0.523)<br>0.516, [0.510, 0.523] | Failure |
| $P(p_b, \text{allosteric site} > 0.95)$ | Site 1: 0.013 (< 0.05)<br>Site 2: 0 (< 0.05) | 0.007 (< 0.05) | Failure |
| $P(p_R, \text{allosteric site} > 0.95)$ | Site 1: 0 (< 0.05)<br>Site 2: 0 (< 0.05) | 0 (< 0.05) | Failure |
| $\overline{p_b^{\text{ref}}}$, allosteric site | Site 1: 0.438 (< 0.5)<br>Site 2: 0.444 (< 0.5) | 0.441 (< 0.5) | Failure |
| $\overline{p_R^{\text{ref}}}$, allosteric site | Site 1: 0.686 (> 0.5)<br>Site 2: 0.668 (> 0.5) | 0.677 (> 0.5) | Success |

It is observed in some cases that some of the allosteric sites of the protein can be detected by a particular measure while the other sites cannot be detected (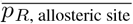 in this case). Therefore, the criteria used here are stringent and would be effective and meaningful in assessing the performance of bond-to-bond propensity analysis and the performance summary is shown in Fig 4.

**Figure 4:**
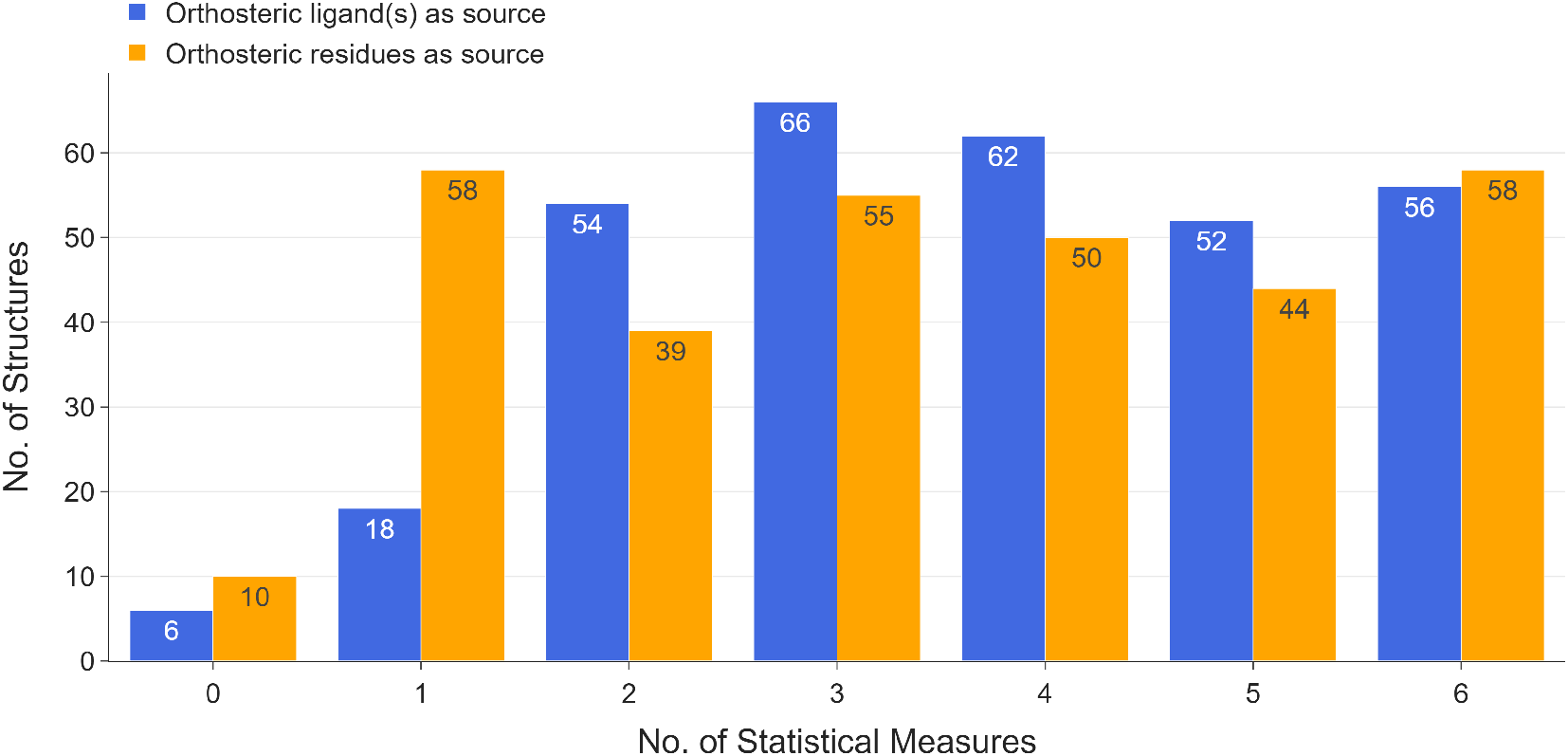
Allosteric site detection results for 314 structures in the CASBench database. The x-axis indicates the number of statistical measures for successful allosteric site detection.

When the orthosteric ligand is selected as the perturbation source, the allosteric site is detected for 308/314 structures (32/33 proteins), according to at least one statistical measure. When using the orthosteric site residues as the source, the allosteric site is detected for 304/314 structures (32/33 proteins), according to at least one statistical measure. It is observed that, in general, the allosteric site of a protein structure can be identified with more statistical measures when the orthosteric ligand is set as the perturbation source.

If the orthosteric ligand is selected as the source, the source bonds include the weak bonds formed by the ligand and the surrounding residues. The orthosteric site includes all residues within 5 Åof the orthosteric ligand [33]. Therefore, the number of source bonds is much lower compared to when using the entire orthosteric site residues as the source. The different and better results obtained by using the ligand as the source suggest that the allosteric site is closely coupled to the ligand-binding event at the orthosteric site. Although successful allosteric site detection is achieved by fewer statistical measures using the whole orthosteric site as the source, the method still succeeds in identifying allosteric sites for more than 96% of the 314 structures. Combined with the results from analysing the ASBench database, for which orthosteric residues are used as the source, the results indicate that propensity analysis reveals the intrinsic coupling of the allosteric site to the region where the orthosteric binding occurs. Using the orthosteric ligand as the perturbation source allows a more accurate detection of allosteric sites. However, if there is no structure containing the orthosteric ligand, the approximate site containing orthosteric residues would still be a good choice to uncover distant sites coupled to the region and provide guidance on allosteric site detection.

### 2.4 Prediction accuracy of bond-to-bond propensity analysis on the CASBench database

We focus here on the six structures for which the allosteric site cannot be detected by any of the measures when using orthosteric ligands as the source. One of them (PDB ID: 4R1R) is ribonucleotide reductase protein R1 (CAS0047). It is a large and complex multimeric protein and only one orthosteric site is reported in the CASBench database. Hence, the effect of cooperativity could affect the performance of propensity analysis as previously discussed. Another two structures (PDB IDs: 1FUQ, 1KQ7) are two out of the four structures of fumarase (CAS0085). This is also a complex multimeric protein where bond-to-bond propensity analysis may not perform well if not all orthosteric ligands are present. The remaining three structures are epoxide hydrolase (CAS0002) (PDB IDs: 5AIA, 5ALN and 5ALT). We analysed 28 structures of epoxide hydrolase in total, each with a different orthosteric ligand. Hence, different ligands, even when binding at the same orthosteric site, exert different perturbation effects on the protein.

When orthosteric residues were used as the perturbation source, the allosteric sites of two structures (PDB IDs: 1LLD, 1LTH) of L-lactate dehydrogenase (CAS0028) were not identified. This can be partly explained by the changed perturbation effects as the allosteric sites were identified when sourcing from the orthosteric ligands. In CASBench, the orthosteric sites include residues within 5 Å from the orthosteric ligands which leads to a large region as the perturbation source. This shows that the specific ligand-site interactions are crucial for accurate allosteric site detection. This is consistent with the overall trend since it has been shown above that successful allosteric site detection is achieved by more statistical measures using the orthosteric ligand as the source. Moreover, allosteric sites of another eight structures were not detected when only using the orthosteric site residues as the source. This further strengthens the idea that the method is sensitive to specific interactions between the ligand and the protein and holds the potential to evaluate the performance of different ligands in the orthosteric site.

## 3 Discussion

Allosteric sites are of great interest in understanding biological function as well as in drug targeting, but, are difficult to predict and in general poorly understood. They are usually discovered serendipitously and require experimental verification. Two recently introduced allosteric protein databases, ASBench [32] and CASBench [33], aim to collect available information on known allosteric sites and are hence excellent benchmarking tools for promising computational approaches. To test the capability of bond-to-bond propensity analysis, a recently developed method that was shown to be able to predict allosteric sites, we deployed the method to both databases, which, after cleaning, provided 432 protein structures for analysis.

An important part of this process is the scoring of the target sites. In addition to previously used scoring measures, we introduced two additional statistical measures, namely the average reference residue quantile score of the allosteric residues, 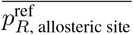 and the proportion of allosteric residues with QS above 0.95, P(*p*_*R*, allosteric site_ > 0.95). The first measures the absolute propensities of residues in the allosteric site compared to the SCOP reference set and the second counts the number of high scoring residues in the allosteric site. These two measures complement the existing four metrics and enable thorough analysis of the significance of the quantile scores computed from bond-to-bond propensity analysis.

Benchmarking datasets of allosteric proteins, namely the ASBench and the CASBench databases, were used for analysis. For structures in ASBench, the orthosteric residues were used as the perturbation source. With the presence of the allosteric ligand, the allosteric site is identified for 106/118 (89.8%) structures and the allosteric site is detected for 99/118 (83.9%) structures when the allosteric ligand is removed, according to at least one statistical measure. Despite the strengthening of functional coupling of the allosteric site to the orthosteric site by the allosteric ligand, propensity analysis is still able to reveal the intrinsic connectivity between the two sites. For the CASBench database we conducted our analysis sourced from the orthosteric ligands or the orthosteric residues and managed to detect the allosteric sites according to at least one statistical measure for 308/314 (98.1%) structures (32/33 proteins) and for 304/314 (96.8%) structures (32/33 proteins), respectively. The allosteric site of a protein structure can be identified with more statistical measures when choosing the orthosteric ligand as the source. This observation suggests that using the ligand as the source confers the perturbation effect of the binding event more accurately. However, if the information on the orthosteric substrate is not available, it is viable to select the orthosteric residues as the perturbation source.

The results presented here strengthen confidence in allosteric site identification as predicted by bond-to-bond propensity, which coupled with the efficiency of the method make it an attractive approach. Generally, the definition of orthosteric and allosteric residues, which would significantly affect the size and residues involved, plays an essential part when evaluating allosteric site prediction methods and was also highlighted for bond-to-bond propensity analysis. Finally, more detailed analysis would be usually required in cases where the allosteric site and the orthosteric site are in very close proximity, to elucidate the effect of cooperativity in large and complex multimeric proteins or the role of structural water molecules, which could still be possible given the computational efficiency of the approach.

## 4 Methods

### 4.1 Allosteric protein datasets

#### The ASBench database

235 X-ray crystal structures of allosteric proteins were downloaded from the ASBench database. Experimentally determined orthosteric and allosteric site residues for these proteins were attained from ASD Release 4.1079. The data was further processed to exclude entries without orthosteric site information or incomplete structures. The resulting 118 structures were all analysed by bond-to-bond propensity. Details can be found in Supplementary Information Table S2. Note that results on the first 4 of the 6 scoring measures were first reported in the supplementary information of reference [61] without any analysis.

#### The CASBench database

X-ray crystal structures containing various orthosteric and allosteric ligands of 91 allosteric proteins in PDB format were downloaded from the CASBench website together with the corresponding experimentally determined orthosteric and allosteric site residues. This data was further processed to exclude incomplete structures and the resulting 314 structures of 33 distinct proteins were used for bond-to-bond propensity analysis. The proteins in CASBench are labelled with CAS ID and the list of proteins with corresponding CAS ID used in this work can be found in Supplementary Information Table S5.

### 4.2 Construction of the atomistic protein graph

Bond-to-bond propensity analysis starts by constructing a weighted atomistic graph using the 3-dimensional coordinates of the atoms of the protein in the PDB files. Atoms are represented by nodes and bond and interactions that link the atoms are represented by edges. The weights of edges correspond to the interaction energies between the atoms with weights derived from relevant interatomic potentials. An in-depth procedure for the atomistic protein graph construction has been described in refs [54, 55]. In this work, Biochemical, atomistic graph construction software in Python for proteins, etc. (BagPype) [68, 61] was used to construct the atomistic protein graph and Fig. 5 illustrates the main features of this process using bovine seminal ribonuclease (11BG) as an example. The crystal structures in the PDB files are cleaned accordingly and hydrogen atoms are added using Reduce (v.3.23) [69], which is incorporated in BagPype. Covalent bonds are weighted using standard bond energies [70]. The weighting of *π − π* stacking, hydrophobic interaction, hydrogen bonding and electrostatic interactions is done based on potentials in references [71, 72, 73], respectively. The weighted graph is then converted to an *N* × *N* adjacency matrix, where *N* is the number of nodes (atoms).

**Figure 5:**
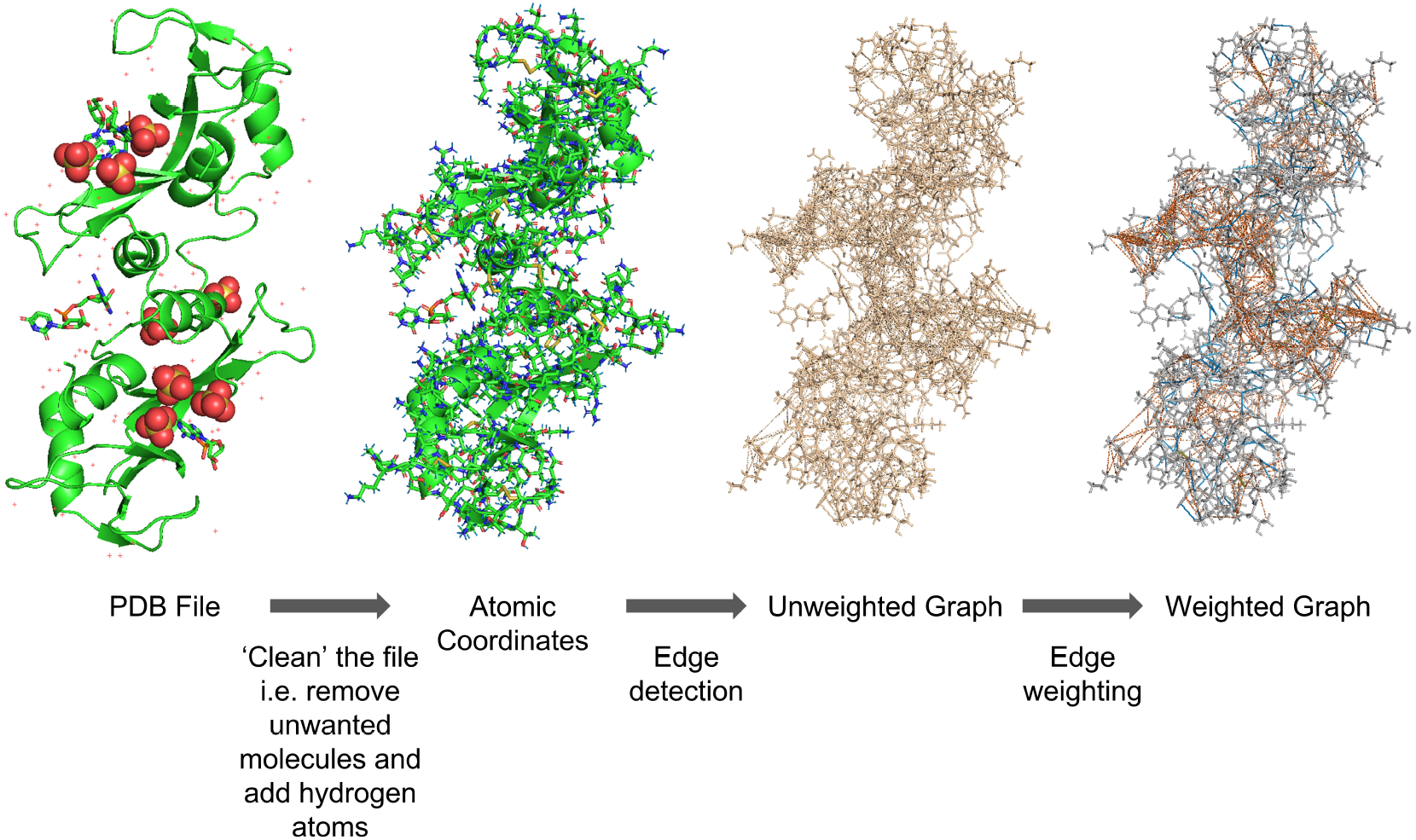
Atomistic graph construction. Main steps of the atomistic protein graph construction package, BagPype, using the structure of bovine seminal ribonuclease (PDB ID: 11BG [62]) as an example.

### 4.3 Bond-to-bond propensities

Bond-to-bond Propensity was first introduced in Ref. [53] and further discussed in Ref. [58], hence it is only briefly summarised here. The edge-to-edge transfer matrix *M* was introduced to study non-local edge-coupling in graphs [74] and an alternative interpretation of *M* is employed to analyse the atomistic protein graph. The element *M_ij_* describes the effect that a perturbation at edge *i* has on edge *j*. *M* is given by

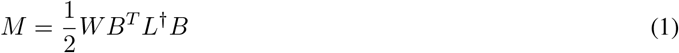

where B is the *n* × *m* incidence matrix for the atomistic protein graph with *n* nodes and *m* edges; *W* = diag(*w_ij_*) is an *m* × *m* diagonal matrix which possesses all edge interaction energies with *w_ij_* as the weight of the edge connecting nodes *i* and *j*, i.e. the bond energy between the atoms. *L*^†^ is the pseudo-inverse of the weighted graph Laplacian matrix *L* [75]. *L*, which defines the diffusion dynamics on the energy-weighted graph [76] and is defined as:

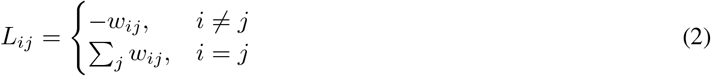

To evaluate the effect of perturbations from a group of bonds *b*′, which belong to the orthosteric ligand or the orthosteric site residues (i.e., the source), on a bond *b* anywhere else in the protein, we calculate:

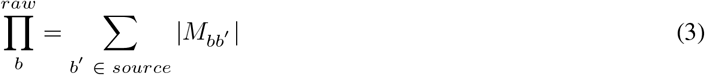

This is the raw propensity of an individual bond which reflects how strongly the bond is coupled to the source. As different proteins contain different numbers of bonds, the raw propensity is normalised and the bond propensity is defined as:

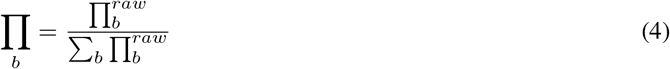

The residue propensity is then defined as the sum of normalised bond propensities of all the bonds of a residue, *R*:

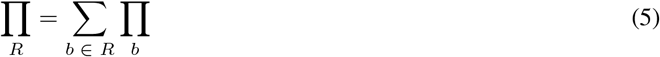

### 4.4 Quantile regression

Bond and residue propensities naturally decrease as the distance of the bond or residue from the perturbation source increases. To determine the bonds and residues that are significant, bond and residue propensities at a similar distance from the source are compared using conditional quantile regression (QR) [77]. The distance of a bond *b* from the perturbation source is defined as the minimum distance, *d_b_*, between *b* and any bond of the source:

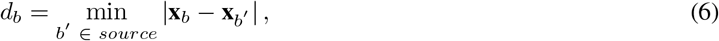

where the vector **x**_*b*_ contains the cartesian coordinates of the midpoint of bond b. As propensity ∏_*b*_ decays exponentially with distance *d*, a linear model for the logarithm of the propensities is adopted to solve the QR minimisation problem:

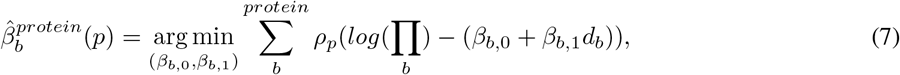

where *ρ_p_*(·) is the tilted absolute value function:

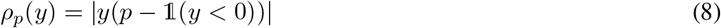

*p* is the quantile and 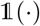 is the indicator function. The optimised model 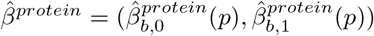 describes the sum of the quantiles of the propensities for all bonds in the protein. The bond quantile score of bond *b* with propensity ∏_*b*_ at distance *d_b_* from the source can be calculate by finding the quantile *p_b_* such that:

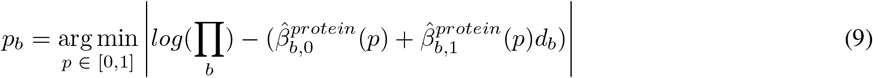

The residue quantile score of residue *R* is defined similarly by using the residue propensity as shown in eq. 5 and the distance *d_p_* which is the minimum distance between the atoms of a residue and those of the source. Therefore,

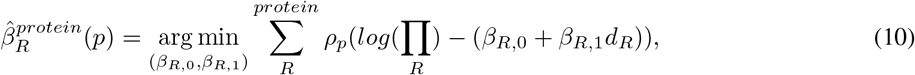

and

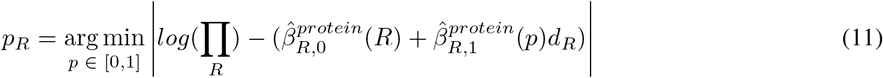

are used to calculate the residue quantile score.

### 4.5 Statistical evaluation of allosteric bond and residue quantile scores (QS)

Four statistical measures have been used to evaluate the significance of the quantile scores (QS) by Amor *et al*. [53] and were employed in this project as listed below:

1. **The average bond quantile score of the allosteric site:**

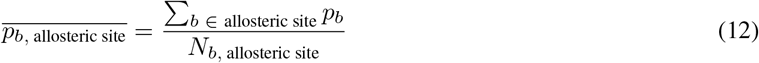

where *N_b_,* allosteric site is the number of bonds in the allosteric site.
2. **The average residue quantile score of the allosteric site:**

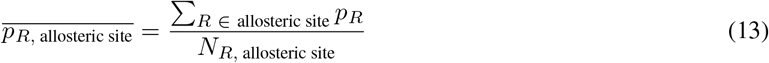

where *N_R_*, allosteric site is the number of residues in the allosteric site.
3. **The proportion of bonds in the allosteric site with bond quantile score greater than 0.95** i.e. P(*p*_*b*, allosteric site_ > 0.95).
4. **The average reference bond quantile score of the allosteric site:**

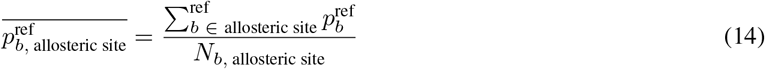

where *N_b_*, allosteric site is the number of bonds in the allosteric site. For the purpose of complementing these previous measures and to investigate more aspects of allosteric site detection, two additional measures were introduced in this work:
5. **The proportion of residues in the allosteric site with residue quantile score greater than 0.95** i.e. P(*p*_*R,* allosteric site_ > 0.95).
6. **The average reference residue quantile score of the allosteric site:**

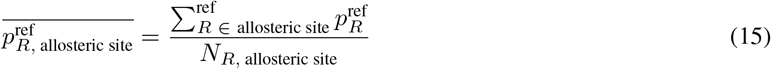

where *N_R_*, allosteric site is the number of residues in the allosteric site.

To assess the significance of the average bond and residue quantile score 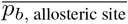 and 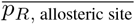, structural bootstrap is used to sample random surrogate sites from the same protein. These surrogate sites need to follow two structural rules: (1) the number of residues is equal to the number of residues in the allosteric site and (2) the diameter (maximum distance between any two atoms in the site) is smaller than that of the allosteric site. For each protein, 1,000 surrogate sites are generated and the average bond and residue quantile scores 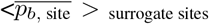 and 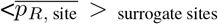 of these sites are calculated. The scores are compared with those of the allosteric sites (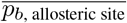 and 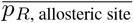). A 95% confidence interval is obtained for each protein to assess the statistical significance by using bootstrap with 10,000 resamples with replacement [78]. Fig 1 illustrates the process using 11BG as an example. If the average quantile score, whether bond or residue of the allosteric residues, is above the upper bound of the 95% confidence interval, the allosteric site is assumed to be detected according to the corresponding statistical measure. The proportion of both bonds and residues of the allosteric residues with a quantile score above 0.95 (P(*p*_*b*, allosteric site_ > 0.95) and P(*p*_*R*, allosteric site_ > 0.95)) is then calculated. If the proportion exceeds the expected proportion of 0.05, the allosteric site is classified as identified. Lastly, the average reference bond and residue quantile scores of the allosteric residues (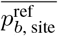 and 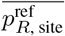) are computed and a value above 0.5 (the expected value) suggests that the allosteric site is uncovered.

## Data availability

All data presented in this study are available upon request.

## Acknowledgements

We acknowledge helpful discussions with Florian Song, Ching Ching Lam and Jerzy Pilipczuk. This work was funded by the President’s PhD Scholarships to N.W.. L.S. acknowledges funding from a Wellcome Trust studentship [grant number 215360/Z/19/Z]. N.W. and S.N.Y. acknowledge funding from the EPSRC award EP/N014529/1 supporting the EPSRC Centre for Mathematics of Precision Healthcare.

## Author contributions

N.W., L.S. and S.N.Y. conceived the study. N.W. performed the computations and created the figures and all authors analysed the data and wrote the manuscript.

## Materials & Correspondence

All requests for data and code shall be directed to.

## References

[1] Casem, M. L. Chapter 3 - Proteins. In Case studies in cell biology, 23–71 (Academic Press, Amsterdam, 2016).

[2] Gonzalez, M. W. & Kann, M. G. Chapter 4: Protein Interactions and Disease. PLOS Computational Biology 8, e1002819 (2012). URL https://doi.org/10.1371/journal.pcbi.1002819.

[3] Santos, R. et al. A comprehensive map of molecular drug targets. Nature reviews. Drug discovery 16, 19–34 (2017). URL https://pubmed.ncbi.nlm.nih.gov/27910877 https://www.ncbi.nlm.nih.gov/pmc/articles/PMC6314433/.

[4] Abdel-Magid, A. F. Allosteric modulators: an emerging concept in drug discovery. ACS medicinal chemistry letters 6, 104–107 (2015). URL https://pubmed.ncbi.nlm.nih.gov/25699154 https://www.ncbi.nlm.nih.gov/pmc/articles/PMC4329591/.

[5] Grover, A. K. Use of Allosteric Targets in the Discovery of Safer Drugs. Medical Principles and Practice 22, 418–426 (2013). URL https://www.karger.com/DOI/10.1159/000350417.

[6] Traxler, P. & Furet, P. Strategies toward the Design of Novel and Selective Protein Tyrosine Kinase Inhibitors. Pharmacology & Therapeutics 82, 195–206 (1999). URL http://www.sciencedirect.com/science/article/pii/S0163725898000448.

[7] Munita, J. M. & Arias, C. A. Mechanisms of Antibiotic Resistance. Microbiology spectrum 4, 0016–2015 (2016). URL https://pubmed.ncbi.nlm.nih.gov/27227291 https://www.ncbi.nlm.nih.gov/pmc/articles/PMC4888801/.

[8] Li, W. et al. Mechanism of tetracycline resistance by ribosomal protection protein Tet(O). Nature communications 4, 1477 (2013). URL https://pubmed.ncbi.nlm.nih.gov/23403578 https://www.ncbi.nlm.nih.gov/pmc/articles/PMC3576927/.

[9] Dönhöfer, A. et al. Structural basis for TetM-mediated tetracycline resistance. Proceedings of the National Academy of Sciences of the United States of America 109, 16900–16905 (2012). URL https://pubmed.ncbi.nlm.nih.gov/23027944 https://www.ncbi.nlm.nih.gov/pmc/articles/PMC3479509/.

[10] Hooper, D. C. Fluoroquinolone resistance among Gram-positive cocci. The Lancet Infectious Diseases 2, 530–538 (2002). URL http://www.sciencedirect.com/science/article/pii/S1473309902003699.

[11] Leclercq, R. Mechanisms of Resistance to Macrolides and Lincosamides: Nature of the Resistance Elements and Their Clinical Implications. Clinical Infectious Diseases 34, 482–492 (2002). URL https://doi.org/10.1086/324626.

[12] Hiramatsu, K. et al. Genomic Basis for Methicillin Resistance in Staphylococcus aureus. Infection & chemotherapy 45, 117–136 (2013). URL https://pubmed.ncbi.nlm.nih.gov/24265961 https://www.ncbi.nlm.nih.gov/pmc/articles/PMC3780952/.

[13] Peracchi, A. & Mozzarelli, A. Exploring and exploiting allostery: Models, evolution, and drug targeting. Biochimica et Biophysica Acta (BBA) - Proteins and Proteomics 1814, 922–933 (2011). URL http://www.sciencedirect.com/science/article/pii/S1570963910002827.

[14] Kenakin, T. & Miller, L. J. Seven transmembrane receptors as shapeshifting proteins: the impact of allosteric modulation and functional selectivity on new drug discovery. Pharmacological reviews 62, 265–304 (2010). URL https://pubmed.ncbi.nlm.nih.gov/20392808 https://www.ncbi.nlm.nih.gov/pmc/articles/PMC2879912/.

[15] De Smet, F., Christopoulos, A. & Carmeliet, P. Allosteric targeting of receptor tyrosine kinases. Nature Biotechnology 32, 1113–1120 (2014). URL https://doi.org/10.1038/nbt.3028.

[16] Christopoulos, A., May, L. T., Avlani, V. A. & Sexton, P. M. G-protein-coupled receptor allosterism: the promise and the problem(s). Biochemical Society Transactions 32, 873–877 (2004). URL https://doi.org/10.1042/BST0320873.

[17] Fox, S. et al. High-Throughput Screening: Update on Practices and Success. Journal of Biomolecular Screening 11, 864–869 (2006). URL https://doi.org/10.1177/1087057106292473.

[18] Andricopulo, A. D., Abraham, L. B. S. & J, D. Structure-Based Drug Design Strategies in Medicinal Chemistry (2009). URL http://www.eurekaselect.com/node/85033/article.

[19] Molek, P., Strukelj, B. & Bratkovic, T. Peptide phage display as a tool for drug discovery: targeting membrane receptors. Molecules (Basel, Switzerland) 16, 857–887 (2011). URL https://pubmed.ncbi.nlm.nih.gov/21258295 https://www.ncbi.nlm.nih.gov/pmc/articles/PMC6259427/.

[20] Nussinov, R. & Tsai, C.-J. Allostery in Disease and in Drug Discovery. Cell 153, 293–305 (2013). URL https://doi.org/10.1016/j.cell.2013.03.034.

[21] Hardy, J. A. & Wells, J. A. Searching for new allosteric sites in enzymes. Current Opinion in Structural Biology 14, 706–715 (2004). URL http://www.sciencedirect.com/science/article/pii/S0959440X0400185X.

[22] Erlanson, D. A., Wells, J. A. & Braisted, A. C. Tethering: Fragment-Based Drug Discovery. Annual Review of Biophysics and Biomolecular Structure 33, 199–223 (2004). URL https://doi.org/10.1146/annurev.biophys.33.110502.140409.

[23] Selvaratnam, R., Chowdhury, S., VanSchouwen, B. & Melacini, G. Mapping allostery through the covariance analysis of NMR chemical shifts. Proceedings of the National Academy of Sciences 108, 6133 LP – 6138 (2011). URL http://www.pnas.org/content/108/15/6133.abstract.

[24] Oyen, D., Wechselberger, R., Srinivasan, V., Steyaert, J. & Barlow, J. N. Mechanistic analysis of allosteric and non-allosteric effects arising from nanobody binding to two epitopes of the dihydrofolate reductase of Escherichia coli. Biochimica et Biophysica Acta (BBA) - Proteins and Proteomics 1834, 2147–2157 (2013). URL http://www.sciencedirect.com/science/article/pii/S1570963913002823.

[25] Rath, V. L. et al. Human liver glycogen phosphorylase inhibitors bind at a new allosteric site. Chemistry & Biology 7, 677–682 (2000). URL http://www.sciencedirect.com/science/article/pii/S1074552100000041.

[26] Wright, S. W. et al. Anilinoquinazoline Inhibitors of Fructose 1,6-Bisphosphatase Bind at a Novel Allosteric Site: Synthesis, In Vitro Characterization, and X-ray Crystallography. Journal of Medicinal Chemistry 45, 3865–3877 (2002). URL https://doi.org/10.1021/jm010496a.

[27] Collier, G. & Ortiz, V. Emerging computational approaches for the study of protein allostery. Archives of Biochemistry and Biophysics 538, 6–15 (2013). URL http://www.sciencedirect.com/science/article/pii/S0003986113002324.

[28] Sheik Amamuddy, O. et al. Integrated Computational Approaches and Tools forAllosteric Drug Discovery. International journal of molecular sciences 21, 847 (2020). URL https://pubmed.ncbi.nlm.nih.gov/32013012 https://www.ncbi.nlm.nih.gov/pmc/articles/PMC7036869/.

[29] Huang, Z. et al. ASD: a comprehensive database of allosteric proteins and modulators. Nucleic Acids Research 39, D663–D669 (2010). URL https://doi.org/10.1093/nar/gkq1022.

[30] Huang, Z. et al. ASD v2.0: updated content and novel features focusing on allosteric regulation. Nucleic Acids Research 42, D510–D516 (2013). URL https://doi.org/10.1093/nar/gkt1247.

[31] Shen, Q. et al. ASD v3.0: unraveling allosteric regulation with structural mechanisms and biological networks. Nucleic Acids Research 44, D527–D535 (2015). URL https://doi.org/10.1093/nar/gkv902.

[32] Huang, W. et al. ASBench: benchmarking sets for allosteric discovery. Bioinformatics 31, 2598–2600 (2015). URL https://doi.org/10.1093/bioinformatics/btv169.

[33] Zlobin, A., Suplatov, D., Kopylov, K. & Švedas, V. CASBench: A Benchmarking Set of Proteins with Annotated Catalytic and Allosteric Sites in Their Structures. Acta naturae 11, 74–80 (2019). URL https://pubmed.ncbi.nlm.nih.gov/31024751 https://www.ncbi.nlm.nih.gov/pmc/articles/PMC6475866/.

[34] Daura, X. Advances in the Computational Identification of Allosteric Sites and Pathways in Proteins BT - Protein Allostery in Drug Discovery. 141–169 (Springer Singapore, Singapore, 2019). URL https://doi.org/10.1007/978-981-13-8719-7_7.

[35] Huang, W. et al. Allosite: a method for predicting allosteric sites. Bioinformatics 29, 2357–2359 (2013). URL https://doi.org/10.1093/bioinformatics/btt399.

[36] Chen, A. S.-Y. et al. A Random Forest Model for Predicting Allosteric and Functional Sites on Proteins. Molecular Informatics 35, 125–135 (2016). URL https://doi.org/10.1002/minf.201500108.

[37] Fogha, J., Diharce, J., Obled, A., Aci-Sèche, S. & Bonnet, P. Computational Analysis of Crystallization Additives for the Identification of New Allosteric Sites. ACS Omega 5, 2114–2122 (2020). URL https://doi.org/10.1021/acsomega.9b02697.

[38] van Gunsteren, W. F. et al. Biomolecular Modeling: Goals, Problems, Perspectives. Angewandte Chemie International Edition 45, 4064–4092 (2006). URL https://doi.org/10.1002/anie.200502655.

[39] Ghosh, A. & Vishveshwara, S. A study of communication pathways in methionyl-tRNA synthetase by molecular dynamics simulations and structure network analysis. Proceedings of the National Academy of Sciences of the United States of America 104, 15711–15716 (2007). URL https://pubmed.ncbi.nlm.nih.gov/17898174 https://www.ncbi.nlm.nih.gov/pmc/articles/PMC2000407/.

[40] Shukla, D., Meng, Y., Roux, B. & Pande, V. S. Activation pathway of Src kinase reveals intermediate states as targets for drug design. Nature Communications 5, 3397 (2014). URL https://doi.org/10.1038/ncomms4397.

[41] Hollingsworth, S. A. & Dror, R. O. Molecular Dynamics Simulation for All. Neuron 99, 1129–1143 (2018). URL http://www.sciencedirect.com/science/article/pii/S0896627318306846.

[42] Dierynck, I. et al. Binding kinetics of darunavir to human immunodeficiency virus type 1 protease explain the potent antiviral activity and high genetic barrier. Journal of virology 81, 13845–13851 (2007).

[43] Dykeman, E. C. & Sankey, O. F. Normal mode analysis and applications in biological physics. Journal of Physics: Condensed Matter 22, 423202 (2010). URL http://dx.doi.org/10.1088/0953-8984/22/42/423202.

[44] Case, D. A. Normal mode analysis of protein dynamics. Current Opinion in Structural Biology 4, 285–290 (1994). URL http://www.sciencedirect.com/science/article/pii/S0959440X94903212.

[45] Bahar, I., Lezon, T. R., Bakan, A. & Shrivastava, I. H. Normal mode analysis of biomolecular structures: functional mechanisms of membrane proteins. Chemical reviews 110, 1463–1497 (2010). URL https://pubmed.ncbi.nlm.nih.gov/19785456 https://www.ncbi.nlm.nih.gov/pmc/articles/PMC2836427/.

[46] Panjkovich, A. & Daura, X. Exploiting protein flexibility to predict the location of allosteric sites. BMC Bioinformatics 13, 273 (2012). URL https://doi.org/10.1186/1471-2105-13-273.

[47] Panjkovich, A. & Daura, X. PARS: a web server for the prediction of Protein Allosteric and Regulatory Sites. Bioinformatics 30, 1314–1315 (2014). URL https://doi.org/10.1093/bioinformatics/btu002.

[48] Greener, J. G. & Sternberg, M. J. E. AlloPred: prediction of allosteric pockets on proteins using normal mode perturbation analysis. BMC Bioinformatics 16, 335 (2015). URL https://doi.org/10.1186/s12859-015-0771-1.

[49] Song, K. et al. Improved Method for the Identification and Validation of Allosteric Sites. Journal of Chemical Information and Modeling 57, 2358–2363 (2017). URL https://doi.org/10.1021/acs.jcim.7b00014.

[50] Guarnera, E. & Berezovsky, I. N. Structure-Based Statistical Mechanical Model Accounts for the Causality and Energetics of Allosteric Communication. PLoS computational biology 12, e1004678–e1004678 (2016). URL https://pubmed.ncbi.nlm.nih.gov/26939022 https://www.ncbi.nlm.nih.gov/pmc/articles/PMC4777440/.

[51] Tee, W.-V., Guarnera, E. & Berezovsky, I. N. Reversing allosteric communication: From detecting allosteric sites to inducing and tuning targeted allosteric response. PLOS Computational Biology 14, e1006228 (2018). URL https://doi.org/10.1371/journal.pcbi.1006228.

[52] Putz, I. & Brock, O. Elastic network model of learned maintained contacts to predict protein motion. PLOS ONE 12, e0183889 (2017). URL https://doi.org/10.1371/journal.pone.0183889.

[53] Amor, B. R., Schaub, M. T., Yaliraki, S. N. & Barahona, M. Prediction of allosteric sites and mediating interactions through bond-to-bond propensities. Nature Communications 7, 1–13 (2016). URL http://dx.doi.org/10.1038/ncomms12477.

[54] Delmotte, A., Tate, E. W., Yaliraki, S. N. & Barahona, M. Protein multi-scale organization through graph partitioning and robustness analysis: application to the myosin–myosin light chain interaction. Physical Biology 8, 55010 (2011). URL http://dx.doi.org/10.1088/1478-3975/8/5/055010.

[55] Amor, B., Yaliraki, S. N., Woscholski, R. & Barahona, M. Uncovering allosteric pathways in caspase-1 using Markov transient analysis and multiscale community detection. Molecular BioSystems 10, 2247–2258 (2014). URL http://dx.doi.org/10.1039/C4MB00088A.

[56] Spielman, D. A. & Teng, S.-H. Nearly-Linear Time Algorithms for Graph Partitioning, Graph Sparsification, and Solving Linear Systems. In Proceedings of the Thirty-Sixth Annual ACM Symposium on Theory of Computing, STOC ’04, 81–90 (Association for Computing Machinery, New York, NY, USA, 2004). URL https://doi.org/10.1145/1007352.1007372.

[57] Kelner, J. A., Orecchia, L., Sidford, A. & Zhu, Z. A. A Simple, Combinatorial Algorithm for Solving SDD Systems in Nearly-Linear Time. In Proceedings of the Forty-Fifth Annual ACM Symposium on Theory of Computing, STOC ’13, 911–920 (Association for Computing Machinery, New York, NY, USA, 2013). URL https://doi.org/10.1145/2488608.2488724.

[58] Hodges, M., Barahona, M. & Yaliraki, S. N. Allostery and cooperativity in multimeric proteins: bond-to-bond propensities in ATCase. Scientific Reports 8, 1–14 (2018). URL http://dx.doi.org/10.1038/s41598-018-27992-z.

[59] Vianello, F. Computational characterisation of protein interaction sites: from small ligand pockets to large domain interfaces. Ph.D. thesis, Imperial College London (2020). URL http://hdl.handle.net/10044/1/89838.

[60] Strömich, L., Wu, N., Barahona, M. & Yaliraki, S. N. Allosteric Hotspots in the Main Protease of SARS-CoV-2. bioRxiv 2020.11.06.369439 (2020). URL https://doi.org/10.1101/2020.11.06.369439.

[61] Mersmann, S. et al. ProteinLens: a web-based application for the analysis of allosteric signalling on atomistic graphs of biomolecules. Nucleic Acids Research (2021). URL https://doi.org/10.1093/nar/gkab350.

[62] Vitagliano, L. et al. A potential allosteric subsite generated by domain swapping in bovine seminal ribonuclease11Edited by A. R. Fersht. Journal of Molecular Biology 293, 569–577 (1999). URL https://www.sciencedirect.com/science/article/pii/S0022283699931583.

[63] Dey, S., Hu, Z., Xu, X. L., Sacchettini, J. C. & Grant, G. A. The Effect of Hinge Mutations on Effector Binding and Domain Rotation in Escherichia coli D-3-Phosphoglycerate Dehydrogenase. Journal of Biological Chemistry 282, 18418–18426 (2007). URL http://www.jbc.org/cgi/content/short/282/25/18418.

[64] Lukacs, C. M. et al. The crystal structure of human muscle glycogen phosphorylase a with bound glucose and AMP: An intermediate conformation with T-state and R-state features. Proteins: Structure, Function, and Bioinformatics 63, 1123–1126 (2006). URL https://doi.org/10.1002/prot.20939.

[65] Ciaccio, C., Coletta, A., De Sanctis, G., Marini, S. & Coletta, M. Cooperativity and allostery in haemoglobin function. IUBMB Life 60, 112–123 (2008). URL https://doi.org/10.1002/iub.6.

[66] Suplatov, D. & Švedas, V. Study of Functional and Allosteric Sites in Protein Superfamilies. Acta naturae 7, 34–45 (2015). URL https://pubmed.ncbi.nlm.nih.gov/26798490 https://www.ncbi.nlm.nih.gov/pmc/articles/PMC4717248/.

[67] Wood, Z. A., Weaver, L. H., Brown, P. H., Beckett, D. & Matthews, B. W. Co-repressor Induced Order and Biotin Repressor Dimerization: A Case for Divergent Followed by Convergent Evolution. Journal of Molecular Biology 357, 509–523 (2006). URL https://www.sciencedirect.com/science/article/pii/S0022283605016426.

[68] Song, F., Barahona, M. & Sophia, Y. N. BagPype: A Python package for the construction of atomistic,491energy-weighted graphs from biomolecular structures. Manuscript in preparation (2020).

[69] Word, J., Lovell, S. C., Richardson, J. S. & Richardson, D. C. Asparagine and glutamine: using hydrogen atom contacts in the choice of side-chain amide orientation11Edited by J. Thornton. Journal of Molecular Biology 285, 1735–1747 (1999). URL http://www.sciencedirect.com/science/article/pii/S0022283698924019.

[70] Huheey, J. E., Keiter, E. A., Keiter, R. L. & Medhi, O. K. Inorganic chemistry: principles of structure and reactivity (Pearson Education India, 2006).

[71] Lin, M. S., Fawzi, N. L. & Head-Gordon, T. Hydrophobic Potential of Mean Force as a Solvation Function for Protein Structure Prediction. Structure 15, 727–740 (2007). URL https://doi.org/10.1016/j.str.2007.05.004.

[72] Mayo, S. L., Olafson, B. D. & Goddard III, W. A. DREIDING: A generic force field for molecular simulations. Journal of Physical Chemistry; (USA) 94 (1990).

[73] Hunter, C. A. & Sanders, J. K. M. The nature of .pi.-.pi. interactions. Journal of the American Chemical Society 112, 5525–5534 (1990). URL https://doi.org/10.1021/ja00170a016.

[74] Schaub, M. T., Lehmann, J., Yaliraki, S. N. & Barahona, M. Structure of complex networks: Quantifying edge-to-edge relations by failure-induced flow redistribution. Network Science 2, 66–89 (2014). URL https://doi.org/10.1017/nws.2014.4.

[75] Biggs, N., Biggs, N. L. & Norman, B. Algebraic graph theory, vol. 67 (Cambridge university press, 1993).

[76] Lambiotte, R., Delvenne, J. & Barahona, M. Random Walks, Markov Processes and the Multiscale Modular Organization of Complex Networks. IEEE Transactions on Network Science and Engineering 1, 76–90 (2014).

[77] Koenker, R. & Hallock, K. F. Quantile Regression. Journal of Economic Perspectives 15, 143–156 (2001). URL https://doi.org/10.1257/jep.15.4.143.

[78] Efron, B. & Tibshirani, R. An introduction to the bootstrap (Chapman and Hall/CRC, New York, NY, United States, 1994), 1st editio edn.

