## Supplementary Information for "Prediction of allosteric sites and signalling: insights from benchmarking datasets"

Wu, N.; Strömich, L. & Yaliraki, S.N.

**Table S1: Performance of other computational methods in prediction of allosteric sites.** These methods have used ASBench [1] and AlloSteric Database (ASD) [2] for method validation.

| Methods | Prediction Accuracy | Remarks |
|---|---|---|
| PARS [3] | 65% | The method was tested on 58 proteins collected from the ASD [2]. |
| AlloPred [4] | 59% | 119 proteins were collected from ASBench [1] and relevant site information were retrieved from UniProt [5] and the Catalytic Site Atlas [6]. Only the chain(s) involving the orthosteric and allosteric sites and the connecting chain(s) were considered i.e. not the whole protein structure. An average prediction accuracy of 59% was achieved when tested on 40 proteins (79 was used for model training). |
| AllositePro [7] | 51.7% | The 147 nonredundant allosteric sites from the Core-Diversity set of ASBench [1] were used in this study and 76 out of 147 allosteric sites was successfully predicted. |
| SBSMMA [8] | Not reported | 41 proteins were selected based on the operational definition of allosteric sites in the paper from ASBench [1]. Predictive power is quantifiled by the area under the ROC curves (AUCs) and 28 out of 48 have an AUC above 0.6. |

### References


[1] Huang, W. *et al.* ASBench: benchmarking sets for allosteric discovery. *Bioinformatics* **31**, 2598–2600 (2015). URL https://doi.org/10.1093/bioinformatics/btv169.

[2] Huang, Z. *et al.* ASD: a comprehensive database of allosteric proteins and modulators. *Nucleic Acids Research* **39**, D663–D669 (2010). URL https://doi.org/10.1093/nar/gkq1022.

[3] Panjkovich, A. & Daura, X. Exploiting protein flexibility to predict the location of allosteric sites. *BMC Bioinformatics* **13**, 273 (2012). URL https://doi.org/10.1186/1471-2105-13-273.

[4] Greener, J. G. & Sternberg, M. J. E. AlloPred: prediction of allosteric pockets on proteins using normal mode perturbation analysis. *BMC Bioinformatics* **16**, 335 (2015). URL https://doi.org/10.1186/s12859-015-0771-1.

[5] Consortium, T. U. UniProt: a hub for protein information. *Nucleic Acids Research* **43**, D204–D212 (2015). URL https://doi.org/10.1093/nar/gku989.

[6] Furnham, N. *et al.* The Catalytic Site Atlas 2.0: cataloging catalytic sites and residues identified in enzymes. *Nucleic Acids Research* **42**, D485–D489 (2014). URL https://doi.org/10.1093/nar/gkt1243.

[7] Song, K. *et al.* Improved Method for the Identification and Validation of Allosteric Sites. *Journal of Chemical Information and Modeling* **57**, 2358–2363 (2017). URL https://doi.org/10.1021/acs.jcim.7b00014.

[8] Tee, W.-V., Guarnera, E. & Berezovsky, I. N. Reversing allosteric communication: From detecting allosteric sites to inducing and tuning targeted allosteric response. *PLOS Computational Biology* **14**, e1006228 (2018). URL https://doi.org/10.1371/journal.pcbi.1006228.

**Table S2*: Details of proteins collected from the ASD and ASBench databases.** For proteins with PDB ID 1CE8, 1Z8D, 2Q8M, 3ETE and 3KGF, there are two distinct allosteric sites reported.

| Protein | PDB | Allosteric Ligands | Allosteric Site Residues | Active Site Residues |
|---|---|---|---|---|
| **Seminal ribonuclease** | 11BG | U2G A131 | A14,A24,A27,A28,A94,A95,B32,B33 | A41,A43,A44,A45,A46,A66,A81,A83,A85,A100,A102,A104,A119,A120,A121,A122,A123,A124,B12 |
| **Pyruvate kinase 1** | 1A3W | FBP A1007 | A401,A402,A403,A404,A405,A406,A407,A408,A452,A459,A483,A484,A490,A491,A492 | A49,A51,A53,A84,A85,A89,A91,A213,A214,A240,A241,A242,A261,A262,A263,A264,A265,A266,A267,A297,A298,A330,A332 |
| **Hemoglobin subunit beta** | 1B86 | DG2 D701 | B145,B225,D545,D625 | A29,A43,A58,A62,A87,A101 |
| **Carbamoyl-phosphate synthase large chain** | 1CE8_1 | ORN A5011 | A783,A791,A793,A892,A893,A895,A907,A1039,A1040,A1041,A1042 | A690,A713,A715,A725,A727,A753,A754,A755,A756,A757,A761,A781,A784,A785,A786,A787,A788,A789,A790,A792,A829,A831,A840,A841,A843,A848,A908,A909,A910 |
| **Carbamoyl-phosphate synthase large chain** | 1CE8_2 | IMP A5012 | A948,A949,A954,A974,A975,A976,A977,A993,A994,A995,A1001,A1015,A1016,A1017,A1025,A1026,A1028,A1029,A1030 | A690,A713,A715,A725,A727,A753,A754,A755,A756,A757,A761,A781,A784,A785,A786,A787,A788,A789,A790,A792,A829,A831,A840,A841,A843,A848,A908,A909,A910 |
| **Ribose-phosphate pyrophosphokinase** | 1DKU | AP2 A1002 | A140,A148,A149,A310,A311,A312,A315,B105,B106,B107,B108,B109 | A99,A101,A102,A103,A104,A106,A107,A110,A135,A174,A227,B138 |
| **Glycogen phosphorylase, liver form** | 1EM6 | CP4 A862 | A37,A38,A40,A60,A63,A64,A67,A188,A189,A190,A191,A229,B38,B40,B60,B63,B64,B67,B188,B189,B190,B191,B229 | A134,A135,A675,A676 |
| **Fructose-1,6-bisphosphatase 1** | 1FRP | AMP A338 | A17,A20,A21,A24,A26,A27,A28,A29,A30,A31,A112,A113,A140,A160,A177 | A121,A122,A124,A125,A212,A215,A244,A246,A247,A248,A249,A250,A251,A252,A262,A264,A269,A274,A275,A280,B241,B242,B243 |
| **Glucose-1-phosphate thymidylyltransferase** | 1G3L | TRH A501 | A45,A114,A115,A116,A117,A118,A119,A120,A250,A251,A255,A256,A259,A293,C216,C218,C219,C220 | A8,A9,A10,A11,A12,A13,A14,A15,A16,A25,A26,A54,A55,A82,A84,A85,A86,A87,A88,A90,A108,A109,A110,A111,A162,A194,A196,A225,A227 |
| **NAD-dependent malic** | 1GZ3 | FUM A605 | A64,A67,A88,A91,A95,B127,B128 | A112,A165,A166,A167,A168,A279,A419,A420, |

| | | | | |
|---|---|---|---|---|
| **enzyme, mitochondrial** | | | | A421,A422,A466,A467 |
| **Glucose-1-phosphate thymidylyltransferase 1** | 1H5S | TMP A1292 | A46,A115,A116,A117,A118,A250,A251,A252,A256,A257,A260,D219,D220,D221 | A8,A9,A10,A11,A12,A13,A14,A15,A16,A25,A26,A54,A55,A82,A84,A85,A86,A87,A88,A90,A108,A109,A110,A111,A162,A194,A196,A225,A227 |
| **Anaerobic ribonucleoside-triphosphate reductase** | 1H78 | DCP A1589 | A98,A99,A100,A102,A103,A104,A107,A110,A111,A114,A146 | A58,A64,A65,A66,A67,A68,A69,A70,A441,A443,A444,A445,A446,A447,A448,A449,A450,A451,A580,A581 |
| **Anthranilate synthase component 1** | 1I7S | TRP A601 | A38,A39,A40,A49,A50,A291,A292,A293,A453,A454,A455,A463,A465 | A258,A259,A260,A262,B11,B56,B57,B58,B59,B60,B61,B62,B84,B85,B86,B87,B89,B107,B133,B134,B135,B136,B137,B170,B172 |
| **Hemoglobin subunit alpha** | 1IWH | PEM A501 | A57,A60,A61,A64,A65,A83 | A29,A43,A58,A62,A87,A101 |
| **Uracil phosphoribosyltransferase** | 1JLR | GTP A303 | A78,A101,A103,A104,A105,A124,A125,A129,A158,C44,C65,C68,D59 | B113,B166,B167,B168,B227,B228,B235,B236,B237 |
| **Phospho-2-dehydro-3-deoxyheptonate aldolase, Phe-sensitive** | 1KFL | PHE A1354 | A150,A151,A154,A178,A179,A180,A209,A211,A214,A221,B6,B7,B10 | A61,A92,A94,A96,A97,A98,A143,A161,A162,A163,A164,A186,A189,A234,A265,A267,A268,A269,A302,A326 |
| **Glucose-1-phosphate thymidylyltransferase** | 1LVW | TYD A3002 | A43,A112,A113,A114,A115,A116,A117,A247,A248,A249,A253,A254,C217 | A8,A9,A10,A11,A12,A13,A14,A15,A16,A25,A26,A54,A55,A82,A84,A85,A86,A87,A88,A90,A108,A109,A110,A111,A162,A194,A196,A225,A227 |
| **Sulfate adenylyltransferase** | 1M8P | PPS A574 | A405,A434,A437,A446,A451,A454,A455,A476,A477,A478,A479,A515,A517,A526,A527,A528,A529,A530 | A196,A197,A198,A199,A200,A205,A206,A209,A265,A267,A276,A289,A290,A291,A292,A293,A294,A295,A296,A330,A331,A332,A333,A334 |
| **Glucose-1-phosphate thymidylyltransferase** | 1MP3 | TTP A501 | A46,A115,A116,A117,A118,A119,A120,A252,A256,A257,A260,B219,B220,B221 | A8,A9,A10,A11,A12,A13,A14,A15,A16,A25,A26,A54,A55,A82,A84,A85,A86,A87,A88,A90,A108,A109,A110,A111,A162,A194,A196,A225,A227 |
| **Glucosamine-6-phosphate isomerase 1** | 1NE7 | 16G B2299 | A1,A2,A258,A262,B151,B152,B158,B159,B160,B161,B184 | A39,A40,A41,A42,A43,A44,A45,A71,A72,A85,A136,A137,A138,A139,A140,A143,A144,A145,A146,A166,A170,A172,A173,A207,A208 |
| **ATP phosphoribosyltransferase** | 1NH8 | HIS A289 | A216,A217,A218,A242,A273,A275 | A11,A12,A70,A71,A88,A89,A90,A116,A155,A156,A157,A158,A159,A160,A161,A162 |

| | | | | |
|---|---|---|---|---|
| **Ornithine decarboxylase** | 1NJJ | GET A601 | A22,A243,A339,A340,A341,A382,A384,A385 | A67,A69,A70,A88,A111,A113,A154,A197,A199,A200,A235,A236,A237,A238,A274,A275,A276,A277,A278,A331,A332,A333,A389,B323,B360,B361,B362,B363 |
| **Phospho-2-dehydro-3-deoxyheptonate aldolase, tyrosine-inhibited** | 1OF6 | DTY A1370 | A162,A166,A169,A193,A194,A195,A224,A226,A227,B21,B22,B25,B28 | A61,A92,A94,A96,A97,A98,A143,A161,A162,A163,A164,A186,A189,A234,A265,A267,A268,A269,A302,A326 |
| **ATP-dependent 6-phosphofructokinase isozyme 1** | 1PFK | ADP A326 | A154,A158,A185,A187,A211,A212,A213,A214,A215,A319,B21,B25,B54,B55,B58,B59 | A9,A10,A11,A12,A41,A71,A72,A73,A74,A75,A76,A77,A82,A101,A102,A103,A104,A105,A106,A107,A108,A109,A111,A124,A125,A129,A171 |
| **Parathion hydrolase** | 1QW7 | EBP A702 | A51,A350 | A55,A57,A131,A132,A201,A202,A230,A254,A301 |
| **Tyrosine-protein phosphatase non-receptor type 1** | 1T49 | 892 A301 | A189,A192,A193,A196,A197,A200,A276,A277,A279,A280,A281,A282 | A45,A46,A47,A48,A49,A111,A115,A120,A180,A181,A182,A215,A216,A217,A218,A219,A220,A221,A222,A262,A266 |
| **NAD(P)-dependent glyceraldehyde-3-phosphate dehydrogenase** | 1UXV | AMP A1503 | A72,A79,A132,A133,A134,A135,A154,A155,A156,A157,A184,A479 | A166,A168,A242,A296,A297,A397 |
| **Cytochrome P450 3A4** | 1W0F | STR A1499 | A213,A214,A217,A219,A220,A240 | A94,A105,A118,A119,A120,A126,A130,A137,A184,A271,A301,A302,A303,A305,A306,A307,A309,A310,A311,A313,A364,A368,A369,A370,A372,A373,A374,A375,A433,A434,A435,A436,A437,A439,A440,A441,A442,A443,A444,A445,A447,A448,A452 |
| **Response regulator PleD** | 1W25 | C2E A503, C2E A505 | A148,A153,A174,A175,A177,A178,A356,A357,A358,A359,A360,A362,A377,A379,A383,A387,A390 | A294,A331,A332,A335,A339,A340,A341,A342,A343,A344,A347,A366,A368,A369,A370,A371 |
| **Acetyl-CoA carboxylase** | 1W96 | S1A A1567 | A69,A73,A76,A77,A389,A392,A393,A396,A397,A398,A454,A485,A487,A510,A512 | A189,A352,A363,A364,A365,A366,A377,A378,A379,A380,A381 |
| **Glycogen phosphorylase, muscle form** | 1Z8D_1 | AMP A900 | A67,A71,A75,A309,A310,A315,A316,A317,A318 | A134,A135,A675,A676 |
| **Glycogen phosphorylase, muscle form** | 1Z8D_2 | ADE A902 | A282,A285,A610,A612,A613 | A134,A135,A675,A676 |

| | | | | |
|---|---|---|---|---|
| **Copper-containing nitrite reductase** | 1ZDS | ACM A2500 | A47,A60,A62,A93,A95,A144,A145,A148,A199 | A98,A100,A106,A135,A137,A142 |
| **4-hydroxy-tetrahydrodipicolinate synthase** | 2ATS | DLY A3003 | B83 | A8,A40,A43,A44,A45,A46,A101,A133,A161,A186,A203,A204,A205,A248 |
| **Aspartate carbamoyltransferase regulatory chain** | 2BE9 | CTP B401 | B14,B15,B20,B22,B23,B64,B88,B90,B93,B95,B98 | A48,A50,A51,A52,A53,A54,A55,A56,A105,A127,A130,A134,A137,A167,A168,A228,A229,A230,A231,A233,A234,A265,A266,A267,A268,A296 |
| **Pyruvate dehydrogenase kinase isozyme 2** | 2BU8 | TF4 A1379 | A53,A80,A111,A112,A115,A154,A157,A158,A161 | A243,A244,A246,A247,A248,A250,A251,A282,A284,A285,A286,A287,A295,A301,A315,A316,A317,A318,A319,A320,A321,A322,A323,A338,A340,A346 |
| **Serum albumin** | 2BXA | C1F A2001 | A150,A199,A214,A218,A219,A222,A223,A238,A242,A257,A260,A264,A290,A291 | A387,A388,A391,A392,A395,A403,A407,A410,A411,A414,A430,A431,A433,A434,A435,A437,A438,A449,A450,A453,A457,A485,A488,A489 |
| **Hemoglobin subunit beta** | 2D60 | L35 B1200 | A36,A99,A100,A103,B35,B37,B108,C95,C137,C141 | A29,A43,A58,A62,A87,A101 |
| **L-asparaginase 1** | 2HIM | ASN A8001 | A162,A240,A271,A272,A273,A301,A302,A303,C240 | A12,A13,A14,A17,A58,A59,A60,A61,A62,A89,A90,A91,A92,A93,A163,C246,C247 |
| **Putative deoxycytidylate deaminase** | 2HVW | DCP A1201 | A21,A23,A43,A44,A46,A47,A49,A50,A53,A75,B108 | A24,A26,A27,A28,A29,A44,A45,A46,A54,A61,A64,A65,A66,A67,A69,A70,A71,A72,A73,A94,A96,A97,A98,A99,A102,A120,A121 |
| **Glycogen phosphorylase, muscle form** | 2IEG | FRY A901 | A60,A63,A67,A188,A190,A191,A192,A194,A229,B37,B38,B39,B40,B53,B57,B185,B186,B188 | A134,A135,A675,A676 |
| **Lysine-sensitive aspartokinase 3** | 2J0X | LYS A1451 | A318,A321,A323,A324,A325,A344,A345,A346,B338,B339,B340 | A8,A9,A10,A11,A12,A13,A39,A199,A202,A219,A220,A221,A222,A223,A225,A226,A227,A228,A229,A230,A231,A232,A251,A255,A256,A257,A258,A259,A301,A302 |
| **Myosin-2 heavy chain** | 2JHR | PBQ A1780 | A265,A420,A423,A424,A427,A428,A431,A590,A592,A617,A618,A619,A620 | A179,A180,A181,A182,A183,A185,A186,A227,A233,A235,A236,A237,A454,A455,A457 |
| **Tryptophan 2,3-dioxygenase** | 2NW8 | TRP A308 | A85,A92,A220,A221,A224,A225,A228 | A51,A55,A113,A117,A123,A124,A125,A248,A252,A253,A254,A255,B24 |

| | | | | |
|---|---|---|---|---|
| **D-3-phosphoglycerate dehydrogenase** | 2PA3 | SER A451 | A344,A346,A347,A348,A349,A350,A351,A370 | A84,A105,A106,A108,A109,A112,A157,A158,A159,A160,A161,A162,A163,A180,A181,A182,A183,A185,A209,A210,A211,A212,A213,A214,A216,A217,A220,A238,A239,A240,A264,A265,A292,A293,A294,A295,A296 |
| **Glutamine--fructose-6-phosphate aminotransferase** | 2PUV | UD1 B5003 | B372,B383,B384,B474,B476,B479,B484,B487,B488,B489,B490,B491,B492 | A403,A404,A405,A406,A449,A450,A451,A452,A453,A454,A455,A458,A483,A501,A502,A503,A510,A587,A588,A591,B604,B605 |
| **Indole-3-pyruvate decarboxylase** | 2Q5O | PPY A5003 | A60,A214,A215,A238,A240,A241,A242,A375,A394,A395,A396,A397 | A380,A401,A402,A403,A461,A462,B23,B24,B25,B71,B74,B113 |
| **Fructose-1,6-bisphosphatase class 1** | 2Q8M_1 | BG6 A340 | A207,A210,A221,A222,A225 | A121,A122,A124,A125,A212,A215,A244,A246,A247,A248,A249,A250,A251,A252,A262,A264,A269,A274,A275,A280,B241,B242,B243 |
| **Fructose-1,6-bisphosphatase class 1** | 2Q8M_2 | AMP A341 | A8,A11,A12,A15,A18,A19,A20,A21,A22,A23,A104,A105,A132,A171 | A121,A122,A124,A125,A212,A215,A244,A246,A247,A248,A249,A250,A251,A252,A262,A264,A269,A274,A275,A280,B241,B242,B243 |
| **Prephenate dehydratase** | 2QMX | PHE A303 | A224,A225,A226,A227,A228,B206,B207,B209,B210,B211,B230,B240,B242 | A52,A53,A54,A55,A56,A57,A79,A80,A81,A82,A83,A166,A167,A168,A169,A170 |
| **Androgen receptor** | 2QPY | 4HY A1 | A723,A724,A727,A826,A829,A830,A833,A834,A837,A840 | A701,A704,A705,A706,A707,A708,A711,A741,A742,A745,A746,A749,A752,A763,A764,A765,A780,A787,A873,A876,A877,A880,A891,A895,A899 |
| **Ribonucleoside-diphosphate reductase 1 subunit alpha** | 2R1R | TTP A762 | A232,A233,A234,A262,A268,A269,A275,A276,B249 | A155,A207,A208,A209,A210,A224,A225,A226,A251,A252,A253,A254,A301,A437,A438,A439,A441,A442,A464,A620,A621,A622,A623,A624,A625,A626,A694 |
| **Acetylglutamate kinase, chloroplastic** | 2RD5 | ARG A1000 | A33,A210,A232,A284,A285,A287,A288,A289,A290,A291,A292,A294 | A41,A74,A75,A76,A77,A80,A81,A93,A96,A97,A98,A100,A108,A112,A156,A181,A192,A193,A194,A195,A196 |
| **Uridylate kinase** | 2V4Y | GTP A1242 | A92,A93,A96,A101,A102,A103,A130,E119,E120,E123,E124,E127,F72,F75 | A15,A16,A17,A18,A19,A55,A56,A57,A58,A59,A61,A62,A63,A64,A73,A76,A77,A78,A80,A84,A136,A137,A138,A139,A140,A141,A142,A143,A144,A145,A146,A147,A148,A163,A201 |

| | | | | |
|---|---|---|---|---|
| **ATP phosphoribosyltransferase** | 2VD3 | HIS A1290 | A235,A236,A237,A238,A240,A254,A255,A256 | A11,A12,A70,A71,A88,A89,A90,A116,A155,A156,A157,A158,A159,A160,A161,A162 |
| **Pyruvate kinase PKLR** | 2VGI | FBP A1574 | A474,A475,A476,A477,A478,A479,A480,A525,A532,A557,A559,A560,A561,A562,A563,A564,A565 | A116,A118,A120,A156,A157,A161,A286,A287,A313,A315,A334,A335,A336,A337,A338,A339,A340,A370,A371,A403,A405 |
| **Glutamate racemase** | 2VVT | I24 A1269 | A14,A15,A38,A152,A153,A155,A157,A187,A190,A246,A250,A254 | A7,A8,A9,A11,A12,A37,A38,A39,A40,A41,A69,A70,A71,A72,A73,A116,A119,A146,A150,A180,A181,A182,A183 |
| **Glutamate racemase** | 2W4I | VGA B1256 | A38,A41,A143,A146,A147,A150,A151,B37,B38,B41,B117,B146,B147,B150,B151 | A7,A8,A9,A11,A12,A37,A38,A39,A40,A41,A69,A70,A71,A72,A73,A116,A119,A146,A150,A180,A181,A182,A183 |
| **Cytosolic purine 5'-nucleotidase** | 2XJC | B4P A1490 | A144,A145,A152,A154,A354,A358,A362,A453,A456,A457 | A52,A53,A54,A55,A56,A65,A151,A155,A157,A158,A161,A202,A205,A206,A207,A209,A210,A214,A215,A248,A249,A250,A251,A252,A255,A292,A348,A351,A356 |
| **Multifunctional 2-oxoglutarate metabolism enzyme** | 2Y0P | ACO A2228 | A822,A827,A830,A1035,A1037,A1038,A1054,A1058,A1060,A1062,A1092,A1142,A1145,A1146,A1147,A1148,A1149,A1150 | A504,A506,A539,A540,A576,A578,A579,A602,A603,A604,A605,A606,A607,A643,A644,A645,A646,A647,A648,A651,A676,A678,A680,A681,A682,A743,A747,B901,B902,B950,B952,B976,B977,B980,B1019,B1020 |
| **Androgen receptor** | 2YHD | AV6 A1921 | A716,A720,A730,A733,A734,A738,A894,A898 | A701,A704,A705,A706,A707,A708,A711,A741,A742,A745,A746,A749,A752,A763,A764,A765,A780,A787,A873,A876,A877,A880,A891,A895,A899 |
| **Androgen receptor** | 2YLO | YLO A1922 | A723,A724,A725,A726,A727,A826,A829,A830,A833,A834 | A701,A704,A705,A706,A707,A708,A711,A741,A742,A745,A746,A749,A752,A763,A764,A765,A780,A787,A873,A876,A877,A880,A891,A895,A899 |
| **Glycogen phosphorylase, muscle form** | 3BCR | AZZ A940 | A282,A285,A610,A612,A613 | A134,A135,A675,A676 |
| **Myosin-2 heavy chain** | 3BZ7 | BL4 A800 | A238,A239,A240,A261,A262,A263,A264,A455,A456,A467,A470,A471,A474,A634,A637,A638 | A179,A180,A181,A182,A183,A185,A186,A227,A233,A235,A236,A237,A454,A455,A457 |
| **Probable aspartokinase** | 3C1N | THR A471 | A414,A416,A417,A419,A420,A421,A440,A444, | A6,A7,A8,A9,A10,A11,A40,A41,A43,A208,A21 |

| | | | | |
|---|---|---|---|---|
| | | | B434,B435 | 0,A211,A228,A229,A230,A231,A232,A234,A235,A236,A237,A238,A239,A240,A241,A260,A264,A265,A266,A267,A268,A311,A312,A313 |
| **Glycogen phosphorylase, liver form** | 3CEH | AVE A833 | A67,A68,A71,A72,A75,A191,A193,A227,B39,B40,B41,B42,B44,B45 | A134,A135,A675,A676 |
| **Chorismate mutase** | 3CSM | TRP A300 | A71,A74,A75,A76,A79,A82,A98,A100 | A12,A16,A19,A157,A164,A168,A192,A193,A194,A195,A197,A198,A201,A234,A238,A239,A240,A242,A243,A246 |
| **Amino-acid acetyltransferase** | 3D2P | ARG A438 | A17,A201,A220,A221,A257,A270,A271,A272,A273,A274,A275,A276,A277,A278,A279,A280,A334 | A307,A312,A354,A355,A356,A357,A358,A359,A363,A364,A365,A366,A367,A368,A369,A370,A371,A391,A392,A394,A395,A396,A397,A398,A399,A401,A402 |
| **D-3-phosphoglycerate dehydrogenase** | 3DC2 | SER A600 | A461,A463,A464,A465,A466,A467,A468,A487 | A84,A105,A106,A108,A109,A112,A157,A158,A159,A160,A161,A162,A163,A180,A181,A182,A183,A185,A209,A210,A211,A212,A213,A214,A216,A217,A220,A238,A239,A240,A264,A265,A292,A293,A294,A295,A296 |
| **Glycogen phosphorylase, muscle form** | 3E3N | AMP A843 | A67,A71,A75,A309,A310,A315,A316,A317,A318,B42,B44,B45 | A134,A135,A675,A676 |
| **Uridylate kinase** | 3EK5 | GTP E2006 | D115,D117,D118,D121,D125,E98,E100,E108,E109,E110,E111,E112,E118,E125,E127 | A15,A16,A17,A18,A19,A55,A56,A57,A58,A59,A61,A62,A63,A64,A73,A76,A77,A78,A80,A84,A136,A137,A138,A139,A140,A141,A142,A143,A144,A145,A146,A147,A148,A163,A201 |
| **Isocitrate dehydrogenase kinase/phosphatase** | 3EPS | AMP A1604 | A101,A104,A105,A113,A116,A291,A294,A295,A298,A375,A376,A377,A378 | A315,A316,A317,A318,A319,A320,A321,A322,A323,A324,A325,A334,A336,A346,A348,A353,A357,A416,A417,A418,A419,A420,A421,A423,A424,A457,A461,A462,A474,A475,A476,A477,A478 |
| **Glutamate dehydrogenase 1, mitochondrial** | 3ETE_1 | GTP A553 | A186,A187,A190,E150,E154,E186,E187,E189,E190 | A111,A114,A126,A167,A168,A211,A349,A374,A377,A378 |
| **Glutamate dehydrogenase 1, mitochondrial** | 3ETE_2 | H3P A552, H3P B552, H3P C552, | A212,A213,A217,A257,A258,A261,A262,A265,A292,A446,A450 | A111,A114,A126,A167,A168,A211,A349,A374,A377,A378 |

| | | | | |
|---|---|---|---|---|
| | | H3P C554, H3P D552, H3P F552 | | |
| **Glutamate dehydrogenase 1, mitochondrial** | 3ETG | GWD A552, GWD E552 | A142,A146,A147,A150,A181,A185 | A111,A114,A126,A167,A168,A211,A349,A374, A377,A378 |
| **Leukotriene A-4 hydrolase** | 3FUD | 692 A710 | A24,A25,A26,A35,A36,A161,A180,A182,A188 | A134,A136,A137,A266,A267,A268,A269,A270, A271,A291,A292,A293,A295,A296,A299,A314, A318,A321,A322,A325,A375,A378,A383,A563, A565 |
| **Casein kinase II subunit alpha** | 3H30 | RFZ A337 | A39,A40,A41,A67,A69,A101,A103,A104,A110 | A45,A46,A47,A48,A51,A53,A66,A68,A95,A113 ,A116,A117,A118,A119,A120,A123,A160,A163, A174,A175 |
| **Pyruvate kinase PKM** | 3H6O | FBP A541 | A431,A432,A433,A434,A436,A437,A482,A489, A514,A516,A517,A518,A519,A520,A521,A522 | A73,A75,A77,A113,A114,A118,A243,A270,A27 2,A291,A292,A293,A294,A295,A296,A297,A32 7,A328,A360,A362 |
| **Glutamate racemase** | 3HFR | 6JZ A270 | A153,A155,A156,A246,A250 | A7,A8,A9,A11,A12,A37,A38,A39,A40,A41,A69 ,A70,A71,A72,A73,A116,A119,A146,A150,A18 0,A181,A182,A183 |
| **Toxin A** | 3HO6 | IHP A270 | A35,A37,A57,A60,A61,A105,A107,A154,A211, A212,A224,A235,A252,A253 | A44,A46,A47,A49,A50,A108,A109,A110,A111, A112,A120,A153,A154,A155,A156,A199,A200, A201,A202,A203,A204,A205,A218 |
| **Pyruvate kinase** | 3HQP | FDP A700 | A399,A400,A401,A402,A404,A405,A453,A456, A480,A481,A485,A486,A487,A488,A489 | A26,A27,A28,A29,A49,A50,A51,A53,A54,A55, A59,A60,A83,A84,A88,A90,A144,A145,A172,A 173,A174,A175,A176,A211,A212,A238,A240,A 264,A296,A330,A331,A332,A334,A335 |
| **Fructose-1,6-bisphosphatase isozyme 2** | 3IFA | AMP A339 | A17,A20,A21,A24,A26,A27,A28,A29,A30,A31, A112,A113,A140,A177 | A121,A122,A124,A125,A212,A215,A244,A246, A247,A248,A249,A250,A251,A252,A262,A264, A269,A274,A275,A280,B241,B242,B243 |
| **HD domain protein** | 3IRH | DGT A458 | A54,A55,A247,A326,A330,A422,B14,B15,B16, B35,B36,B41,B44,B64 | B48,B49,B50,B51,B52,B63,B66,B111,B114,B11 8,B119,B122,B129,B183,B184,B187,B191,B235 ,B239,B242,B243,B244,B248,B252,B368,B369 |
| **Tyrosine-protein kinase ABL1** | 3K5V | STJ A1 | A356,A359,A360,A363,A448,A451,A452,A454, A481,A482,A483,A484,A487,A512,A521,A525, | A267,A272,A275,A288,A289,A290,A305,A308, A309,A312,A317,A318,A332,A333,A334,A335, |

| | | | | |
|---|---|---|---|---|
| | | | A529 | A336,A337,A338,A339,A340,A341,A373,A378,A379,A380,A381,A389,A398,A399,A400,A401 |
| **Phospho-2-dehydro-3-deoxyheptonate aldolase AroG** | 3KGF_1 | PHE A9003 | A91,A92,A171,A174,A175,A178,B3,B5,B6,B55,B173 | A87,A126,A130,A134,A248,A280,A281,A282,A283,A284,A306,A307,A337,A366,A369,A409,A411,A441 |
| **Phospho-2-dehydro-3-deoxyheptonate aldolase AroG** | 3KGF_2 | TRP A9004 | A107,A110,A111,A123,A192,A194,A237,A238,A240,A241 | A87,A126,A130,A134,A248,A280,A281,A282,A283,A284,A306,A307,A337,A366,A369,A409,A411,A441 |
| **Glutamate receptor 3** | 3LSW | 4MP A801 | A105,A106,A107,A108,A217,A218,A219 | A61,A62,A74,A89,A90,A91,A96,A111,A137,A138,A140,A141,A142,A143,A144,A174,A191,A192,A193,A196,A220 |
| **Glutamate receptor 3** | 3LSX | PZI A802 | A248,A252 | A61,A62,A74,A89,A90,A91,A96,A111,A137,A138,A140,A141,A142,A143,A144,A174,A191,A192,A193,A196,A220 |
| **Serum albumin** | 3LU6 | IMX A587 | A209,A212,A213,A216,A232,A235,A324,A327,A328,A347,A351,A354 | A387,A388,A391,A392,A395,A403,A407,A410,A411,A414,A430,A431,A433,A434,A435,A437,A438,A449,A450,A453,A457,A485,A488,A489 |
| **Glutamate receptor 3** | 3M3F | P99 A800 | A92,A104,A105,A106,A107,A108,A217,A218,A219,A239,A242,A247 | A61,A62,A74,A89,A90,A91,A96,A111,A137,A138,A140,A141,A142,A143,A144,A174,A191,A192,A193,A196,A220 |
| **Glutamate dehydrogenase 1, mitochondrial** | 3MW9 | NAI A604 | A195,A205,A206,A387,A388,A391,A392,A393,B85,B86,B115,B116,B119,B120,B121,B488,B491 | A111,A114,A126,A167,A168,A211,A349,A374,A377,A378 |
| **Prephenate dehydratase** | 3MWB | PHE B311 | A226,A227,A228,A229,A230,B205,B208,B209,B210,B211,B212,B213,B232,B242,B244 | A52,A53,A54,A55,A56,A57,A79,A80,A81,A82,A83,A166,A167,A168,A169,A170 |
| **Pyruvate kinase PKM** | 3N25 | PRO A1200 | A42,A43,A69,A105,A463,A465,A467,A468,A469,A470 | A73,A75,A77,A113,A114,A118,A243,A270,A272,A291,A292,A293,A294,A295,A296,A297,A327,A328,A360,A362 |
| **Mitogen-activated protein kinase 8** | 3O2M | 46A A701 | A178,A180,A197,A198,A199,A230,A231,A234,A253,A255,A256,A259,B184,B255 | A32,A33,A40,A52,A53,A55,A86,A108,A109,A110,A111,A112,A113,A114,A158,A168 |
| **Phospho-2-dehydro-3-deoxyheptonate aldolase** | 3PG9 | TYR A339 | A31,A33,A34,A35,A36,A38,F1,F2,F40,F41,F42,F43,F45,F65,F66 | A102,A131,A132,A164,A186,A247,A272,A309 |
| **UDP-glucose 6-** | 3PJG | UGA A902 | A256,A257,A259,A284,A319,A320,A322,A323, | A131,A161,A162,A163,A164,A165,A220,A275, |

| | | | | |
|---|---|---|---|---|
| **dehydrogenase** | | | A324,A326,A345 | A276,A280 |
| **UDP-glucose 6-dehydrogenase** | 3PTZ | UDX A501 | A131,A161,A162,A163,A164,A165,A220,A224,A227,A231,A265,A266,A267,A269,A272,A273,A276,A277,A338,A339,A442,B260 | A131,A161,A162,A163,A164,A165,A220,A275,A276,A280 |
| **Cyclin-dependent kinase 2** | 3PXF | 2AN A304, 2AN A305 | A15,A33,A35,A37,A52,A55,A56,A64,A66,A69,A71,A76,A78,A80,A144,A145,A146,A154 | A10,A11,A12,A13,A14,A15,A16,A17,A18,A31,A33,A64,A80,A81,A82,A83,A84,A85,A86,A89,A127,A129,A130,A131,A132,A133,A134,A144,A145,A162 |
| **Tyrosine-protein kinase ABL1** | 3PYY | 3YY A538 | A356,A359,A360,A363,A448,A451,A452,A481,A482,A483,A484,A487 | A267,A272,A275,A288,A289,A290,A305,A308,A309,A312,A317,A318,A332,A333,A334,A335,A336,A337,A338,A339,A340,A341,A373,A378,A379,A380,A381,A389,A398,A399,A400,A401 |
| **Prostaglandin G/H synthase 2** | 3QH0 | PLM A625 | A116,A120,A205,A348,A349,A353,A355,A385,A387,A523,A526,A527,A530,A531 | B75,B102,B106,B191,B330,B334,B335,B338,B339,B341,B345,B367,B370,B371,B373,B504,B508,B509,B510,B511,B512,B513,B514,B516,B517,B520 |
| **Ribonucleoside-diphosphate reductase 1 subunit alpha** | 3R1R | ATP A762 | A9,A15,A16,A17,A18,A21,A22,A25,A55,A59,A91 | A155,A207,A208,A209,A210,A224,A225,A226,A251,A252,A253,A254,A301,A437,A438,A439,A441,A442,A464,A620,A621,A622,A623,A624,A625,A626,A694 |
| **Glutaminase kidney isoform, mitochondrial** | 3UO9 | 04A B2 | A320,A321,A322,A323,A324,A325,A327,A394,B320,B321,B322,B323,B324,B325,B394,D317 | A249,A284,A285,A286,A287,A289,A335,A381,A387,A388,A414,A415,A418,A466,A482,A483,A484,A485 |
| **Kinesin-like protein KIF11** | 3ZCW | 4A2 A1366 | A104,A266,A269,A270,A287,A288,A289,A292,A293,A295,A296,A297,A299,A300,A332,A352,A353,A355,A356 | A24,A25,A26,A27,A74,A76,A78,A105,A106,A107,A108,A109,A110,A111,A112,A113,A114,A118,A132,A232,A233,A265,A335 |
| **Glutamate racemase** | 4B1F | KRH A1256 | A10,A11,A13,A17,A149,A150,A152,A154,A182,A183,A186,A248,A252,A253 | A7,A8,A9,A11,A12,A37,A38,A39,A40,A41,A69,A70,A71,A72,A73,A116,A119,A146,A150,A180,A181,A182,A183 |
| **Pyruvate kinase PKM** | 4B2D | SER A1532 | A43,A44,A45,A46,A70,A106,A464,A468,A469,A470,A471 | A73,A75,A77,A113,A114,A118,A243,A270,A272,A291,A292,A293,A294,A295,A296,A297,A327,A328,A360,A362 |
| **Kinesin-like protein KIF11** | 4BBG | V02 A1370 | A112,A116,A117,A118,A119,A130,A132,A133, | A24,A25,A26,A27,A74,A76,A78,A105,A106,A1 |

| | | | | |
|---|---|---|---|---|
| | | | A137,A211,A214,A215,A218,A221 | 07,A108,A109,A110,A111,A112,A113,A114,A118,A132,A232,A233,A265,A335 |
| **5'-AMP-activated protein kinase catalytic subunit alpha-2** | 4CFE | 992 A1553 | A11,A18,A24,A28,A29,A31,A46,A48,A88,A90,B81,B83,B106,B107,B111,B113 | A22,A23,A24,A25,A26,A30,A43,A45,A64,A77,A91,A93,A94,A95,A96,A97,A98,A99,A100,A141,A143,A144,A145,A146,A156,A157,A158 |
| **Mitogen-activated protein kinase 14** | 4E6C | 0O8 A500 | A195,A196,A197,A255 | A30,A33,A38,A39,A40,A51,A52,A53,A71,A75,A84,A85,A86,A88,A104,A105,A106,A107,A108,A109,A110,A111,A112,A115,A154,A156,A157,A158,A167,A168 |
| **Pyruvate kinase PKM** | 4G1N | NZT A603 | A26,A30,A353,A354,A389,A390,A393,A394,A397,B26,B27,B30,B311,B353,B354,B390,B394 | A73,A75,A77,A113,A114,A118,A243,A270,A272,A291,A292,A293,A294,A295,A296,A297,A327,A328,A360,A362 |
| **2-dehydro-3-deoxyphosphoheptonate aldolase** | 4GRS | TYR A401 | A1,A40,A41,A42,A43,A45,A66,C31,C33,C34,C35,C36,C38 | A102,A131,A132,A164,A186,A247,A272,A309 |
| **Glucose-1-phosphate thymidylyltransferase** | 4HO6 | UTP A301 | A43,A112,A113,A114,A115,A116,A117,A248,A249,A253,A254,A257 | A8,A9,A10,A11,A12,A13,A14,A15,A16,A25,A26,A54,A55,A82,A84,A85,A86,A87,A88,A90,A108,A109,A110,A111,A162,A194,A196,A225,A227 |
| **Pyruvate kinase 1** | 4HYW | FDP A503 | A400,A401,A402,A403,A405,A406,A454,A457,A481,A482,A486,A487,A488,A489,A490 | A50,A52,A54,A84,A85,A89,A212,A213,A239,A241,A260,A261,A262,A263,A264,A265,A266,A296,A297,A329,A331 |
| **Isocitrate dehydrogenase [NADP], mitochondrial** | 4JA8 | 1K9 A502 | A164,A294,A297,A298,A306,A311,A312,A315,A316,A319,A320,B160,B164,B294,B297,B298,B306,B311,B312,B315,B316,B319,B320 | A57,A59,A62,A112,A113,A114,A115,A116,A117,A118,A122,A136,A326,A327,A328,A345,A346,A347,A348,A349,A350,A351,A352,A353,A354,A365,A366,A367,A368,A412,A414,A422 |
| **Glutaminase kidney isoform, mitochondrial** | 4JKT | 04A D701 | A322,A325,A326,A327,A328,A329,A330,A399,C322,D322,D325,D326,D327,D328,D329,D330,D399 | A249,A284,A285,A286,A287,A289,A335,A381,A387,A388,A414,A415,A418,A466,A482,A483,A484,A485 |
| **N-acetylglutamate kinase / N-acetylglutamate synthase** | 4KZT | ARG A501 | A28,A206,A225,A265,A277,A278,A280,A281,A282,A283,A285,A286,A287,A365 | A307,A312,A354,A355,A356,A357,A358,A359,A363,A364,A365,A366,A367,A368,A369,A370,A371,A391,A392,A394,A395,A396,A397,A398,A399,A401,A402 |

| | | | | |
|---|---|---|---|---|
| **HD domain protein** | 4LRL | TTP B503 | A55,A56,A197,A241,A245,B15,B16,B17,D206,D209,D223 | B48,B49,B50,B51,B52,B63,B66,B111,B114,B118,B119,B122,B129,B183,B184,B187,B191,B235,B239,B242,B243,B244,B248,B252,B368,B369 |
| **Glycogen phosphorylase, muscle form** | 4MRA | QUE A901 | A121,A124,A495,A544,A545,A548,A551,A552,A655 | A134,A135,A675,A676 |
| **ATP-dependent 6-phosphofructokinase** | 4PFK | ADP A326 | A154,A185,A187,A211,A212,A213,A214,A215 | A9,A10,A11,A12,A41,A71,A72,A73,A74,A75,A76,A77,A82,A101,A102,A103,A104,A105,A106,A107,A108,A109,A111,A124,A125,A129,A171 |

* This table has been recently published in Ref. [1] and is included here to facilitate ease of reading.

[1] S. F. Mersmann *et al.*, “ProteinLens: a web-based application for the analysis of allosteric signalling on atomistic graphs of biomolecules,” *Nucleic Acids Res.*, May 2021.

**Table S3*: Allosteric site quantile scores of proteins in Table S2 (with the presence of allosteric ligands in the structures).** The results from six statistical scores described in Methods. Average site residue and bond quantile scores are compared with those of 1000 surrogate sites of the same size. The difference is shown in bold if it is above 0 and starred if it is above the 95% confidence interval. The proportion of residues/bonds with $p_{R/b,\text{allo}} > 0.95$ and the average reference quantile score $\overline{p^{\text{ref}}_{R/b,\text{ allo}}}$ are shown in bold if they are above the expected values of 0.05 and 0.5 respectively.

| Protein | PDB | $\overline{p_{R,\text{allo}}} - \langle\overline{p_{R,\text{site}}}\rangle_{\text{surr}}$ | $\overline{p_{b,\text{allo}}} - \langle\overline{p_{b,\text{site}}}\rangle_{\text{surr}}$ | $P(p_{R,\text{allo}} > 0.95)$ | $P(p_{b,\text{allo}} > 0.95)$ | $\overline{p^{\text{ref}}_{R,\text{ allo}}}$ | $\overline{p^{\text{ref}}_{b,\text{ allo}}}$ | Summary |
|---|---|---|---|---|---|---|---|---|
| **Fructose-1,6-bisphosphatase class 1** | 2Q8M_2 | **0.37*** | **0.19*** | **0.43** | **0.19** | **0.66** | **0.54** | ●●●●●● |
| **Cytochrome P450 3A4** | 1W0F | **0.37*** | **0.049*** | **0.17** | 0.0078 | **0.71** | **0.5** | ●●●●○● |
| **Tryptophan 2,3-dioxygenase** | 2NW8 | **0.34*** | **0.098*** | **0.43** | **0.13** | **0.83** | **0.58** | ●●●●●● |
| **Parathion hydrolase** | 1QW7 | **0.32*** | -0.15 | 0 | 0.022 | **0.78** | **0.51** | ●○○○●● |
| **Glutamate dehydrogenase 1, mitochondrial** | 3ETG | **0.3*** | **0.16*** | 0 | **0.085** | **0.65** | **0.75** | ●●○●●● |
| **Lysine-sensitive aspartokinase 3** | 2J0X | **0.29*** | **0.18*** | **0.36** | **0.11** | **0.63** | **0.62** | ●●●●●● |
| **Aspartate carbamoyltransferase regulatory chain** | 2BE9 | **0.29*** | **0.25*** | **0.091** | **0.1** | 0.096 | 0.11 | ●●●●○○ |
| **Copper-containing nitrite reductase** | 1ZDS | **0.28*** | **0.13*** | **0.22** | **0.12** | **0.81** | **0.78** | ●●●●●● |
| **Hemoglobin subunit beta** | 1B86 | **0.26*** | -0.0082 | **0.75** | **0.24** | **0.77** | **0.54** | ●○●●●● |
| **Androgen receptor** | 2QPY | **0.26*** | **0.21*** | **0.3** | **0.11** | **0.73** | **0.53** | ●●●●●● |
| **Tyrosine-protein kinase ABL1** | 3K5V | **0.25*** | **0.15*** | **0.18** | 0.034 | **0.7** | **0.63** | ●●●○●● |
| **Fructose-1,6-bisphosphatase isozyme 2** | 3IFA | **0.25*** | **0.1*** | **0.14** | **0.16** | **0.61** | **0.61** | ●●●●●● |
| **Tyrosine-protein kinase** | 3PYY | **0.21*** | **0.1*** | **0.17** | 0.033 | **0.62** | **0.59** | ●●●○●● |

| | | | | | | | | |
|---|---|---|---|---|---|---|---|---|
| **ABL1** | | | | | | | | |
| **Glutamate receptor 3** | 3LSX | **0.21*** | **0.011*** | 0 | **0.2** | **0.7** | 0.41 | ●●○●●○ |
| **HD domain protein** | 3IRH | **0.2*** | **0.035*** | **0.21** | **0.099** | **0.62** | **0.66** | ●●●●●● |
| **Isocitrate dehydrogenase kinase/phosphatase** | 3EPS | **0.19*** | **0.16*** | **0.15** | 0.047 | **0.68** | **0.76** | ●●●○●● |
| **Pyruvate kinase PKM** | 4G1N | **0.19*** | **0.013*** | 0 | 0.029 | **0.65** | **0.71** | ●●○○●● |
| **Serum albumin** | 3LU6 | **0.19*** | **0.12*** | **0.083** | **0.072** | **0.72** | **0.66** | ●●●●●● |
| **Toxin A** | 3HO6 | **0.19*** | **0.022*** | **0.29** | **0.13** | **0.75** | **0.6** | ●●●●●● |
| **Indole-3-pyruvate decarboxylase** | 2Q5O | **0.18*** | **0.052*** | **0.17** | **0.12** | **0.65** | **0.7** | ●●●●●● |
| **Glycogen phosphorylase, muscle form** | 1Z8D_1 | **0.18*** | -0.07 | **0.11** | 0.038 | **0.63** | 0.41 | ●○●○●○ |
| **Glycogen phosphorylase, liver form** | 1EM6 | **0.17*** | **0.09*** | **0.13** | **0.062** | **0.74** | **0.73** | ●●●●●● |
| **Pyruvate dehydrogenase kinase isozyme 2** | 2BU8 | **0.17*** | **0.074*** | 0 | **0.065** | **0.83** | **0.65** | ●●○●●● |
| **Anthranilate synthase component 1** | 1I7S | **0.17*** | **0.093*** | 0 | 0.04 | **0.56** | **0.61** | ●●○○●● |
| **Casein kinase II subunit alpha** | 3H30 | **0.16*** | **0.098*** | 0 | **0.086** | **0.77** | **0.75** | ●●○●●● |
| **Pyruvate kinase PKM** | 3N25 | **0.16*** | **0.07*** | 0 | **0.091** | **0.57** | **0.73** | ●●○●●● |
| **Carbamoyl-phosphate synthase large chain** | 1CE8_1 | **0.16*** | **0.036*** | **0.091** | **0.12** | **0.56** | **0.65** | ●●●●●● |
| **Fructose-1,6-bisphosphatase class 1** | 2Q8M_1 | **0.16*** | -0.045 | 0 | 0.015 | **0.62** | 0.48 | ●○○○●○ |
| **ATP-dependent 6-phosphofructokinase** | 4PFK | **0.16*** | **0.091*** | **0.12** | **0.12** | **0.56** | 0.41 | ●●●●●● |
| **ATP phosphoribosyltransferase** | 1NH8 | **0.16*** | **0.089*** | 0 | **0.065** | 0.087 | 0.087 | ●●○●○○ |
| **Glutamate dehydrogenase** | 3ETE_1 | **0.15*** | **0.074*** | 0 | 0.038 | **0.58** | **0.73** | ●●○○●● |

| | | | | | | | | |
|---|---|---|---|---|---|---|---|---|
| 1, mitochondrial | | | | | | | | |
| Ribonucleoside-diphosphate reductase 1 subunit alpha | 2R1R | **0.15*** | **0.054*** | **0.22** | **0.13** | **0.55** | **0.63** | ●●●●●● |
| Glycogen phosphorylase, liver form | 3CEH | **0.14*** | -0.0044 | 0 | **0.078** | **0.72** | **0.67** | ●○○●●● |
| Acetyl-CoA carboxylase | 1W96 | **0.14*** | -0.021 | 0 | **0.057** | **0.67** | **0.66** | ●○○●●● |
| Uridylate kinase | 2V4Y | **0.14*** | **0.054*** | 0 | **0.11** | **0.61** | **0.66** | ●●○●●● |
| Uridylate kinase | 3EK5 | **0.14*** | **0.0054*** | **0.2** | **0.11** | **0.68** | **0.62** | ●●●●●● |
| Seminal ribonuclease | 11BG | **0.14*** | **0.033*** | **0.12** | **0.081** | **0.78** | **0.51** | ●●●●●● |
| Serum albumin | 2BXA | **0.14*** | **0.018*** | **0.14** | **0.062** | **0.68** | **0.51** | ●●●●●● |
| Tyrosine-protein phosphatase non-receptor type 1 | 1T49 | **0.14*** | **0.025*** | **0.25** | 0.045 | **0.74** | 0.49 | ●●●○●● |
| Glutamate racemase | 3HFR | **0.13*** | **0.048*** | **0.2** | 0.038 | **0.64** | **0.62** | ●●●○●● |
| Leukotriene A-4 hydrolase | 3FUD | **0.13*** | -0.058 | 0 | **0.067** | **0.54** | 0.41 | ●○○●●○ |
| 5'-AMP-activated protein kinase catalytic subunit alpha-2 | 4CFE | **0.12*** | **0.15*** | **0.38** | **0.069** | **0.62** | **0.73** | ●●●●●● |
| Phospho-2-dehydro-3-deoxyheptonate aldolase | 3PG9 | **0.12*** | **0.085*** | **0.13** | **0.11** | **0.7** | **0.72** | ●●●●●● |
| Glucose-1-phosphate thymidylyltransferase 1 | 1H5S | **0.12*** | **0.1*** | **0.21** | **0.098** | **0.59** | **0.72** | ●●●●●● |
| Androgen receptor | 2YHD | **0.12*** | **0.07*** | **0.12** | 0.037 | **0.76** | **0.56** | ●●●○●● |
| Androgen receptor | 2YLO | **0.12*** | **0.08*** | 0 | 0.014 | **0.63** | 0.47 | ●●○○●○ |
| Pyruvate kinase PKM | 4B2D | **0.1*** | **0.11*** | **0.091** | **0.13** | **0.54** | **0.74** | ●●●●●● |
| Glucosamine-6-phosphate isomerase 1 | 1NE7 | **0.1*** | **0.064*** | 0 | **0.1** | **0.58** | **0.66** | ●●○●●● |
| Uracil phosphoribosyltransferase | 1JLR | **0.1*** | **0.012*** | **0.23** | **0.082** | **0.75** | **0.65** | ●●●●●● |

| | | | | | | | | |
|---|---|---|---|---|---|---|---|---|
| **Glucose-1-phosphate thymidylyltransferase** | 1LVW | **0.1*** | -0.027 | 0 | **0.07** | **0.55** | **0.6** | ●○○●●● |
| **Fructose-1,6-bisphosphatase 1** | 1FRP | **0.1*** | **0.028*** | 0 | **0.067** | **0.61** | **0.54** | ●●○●●● |
| **Carbamoyl-phosphate synthase large chain** | 1CE8_2 | **0.096*** | **0.063*** | **0.11** | **0.15** | **0.56** | **0.68** | ●●●●●● |
| **Hemoglobin subunit beta** | 2D60 | **0.09*** | **0.057*** | **0.1** | 0.034 | **0.73** | **0.66** | ●●●○●● |
| **L-asparaginase 1** | 2HIM | **0.077*** | **0.14*** | **0.11** | **0.097** | **0.55** | **0.76** | ●●●●●● |
| **Response regulator PleD** | 1W25 | **0.07*** | **0.083*** | **0.059** | **0.14** | **0.74** | **0.73** | ●●●●●● |
| **Putative deoxycytidylate deaminase** | 2HVW | **0.063*** | -0.16 | 0 | **0.13** | **0.72** | 0.39 | ●○○●●○ |
| **Prephenate dehydratase** | 2QMX | **0.062*** | -0.01 | 0 | 0.042 | **0.69** | **0.65** | ●○○○●● |
| **Glycogen phosphorylase, muscle form** | 3E3N | **0.057*** | -0.092 | **0.083** | 0.02 | **0.63** | **0.57** | ●○●○●● |
| **Mitogen-activated protein kinase 8** | 3O2M | **0.056*** | -0.0067 | 0 | 0.0047 | **0.62** | **0.57** | ●○○○●● |
| **Isocitrate dehydrogenase [NADP], mitochondrial** | 4JA8 | **0.051*** | -0.078 | 0.043 | 0.0028 | **0.5** | 0.48 | ●○○○●○ |
| **Ribose-phosphate pyrophosphokinase** | 1DKU | **0.047*** | **0.034*** | **0.083** | **0.17** | **0.6** | **0.62** | ●●●●●● |
| **Phospho-2-dehydro-3-deoxyheptonate aldolase AroG** | 3KGF_2 | **0.034*** | **0.011*** | **0.2** | 0.046 | **0.51** | **0.57** | ●●●○●● |
| **Chorismate mutase** | 3CSM | **0.033*** | **0.00024*** | 0 | 0.012 | **0.57** | 0.46 | ●●○○●○ |
| **Cytosolic purine 5'-nucleotidase** | 2XJC | **0.028*** | **0.051*** | **0.2** | **0.1** | **0.58** | **0.58** | ●●●●●● |
| **Glucose-1-phosphate thymidylyltransferase** | 1G3L | **0.019*** | **0.052*** | 0 | **0.071** | **0.58** | **0.71** | ●●○●●● |
| **Glutamate receptor 3** | 3M3F | **0.017*** | **0.032*** | **0.17** | 0.045 | **0.65** | **0.51** | ●●●○●● |
| **Probable aspartokinase** | 3C1N | **0.013*** | -0.12 | 0 | 0 | 0.36 | 0.47 | ●○○○○○ |

| | | | | | | | | |
|---|---|---|---|---|---|---|---|---|
| **ATP-dependent 6-phosphofructokinase isozyme 1** | 1PFK | **0.011*** | -0.096 | **0.062** | **0.059** | **0.57** | 0.46 | ●○●●●○ |
| **Mitogen-activated protein kinase 14** | 4E6C | **0.011*** | -0.027 | 0 | **0.11** | 0.27 | 0.24 | ●○○●○○ |
| **Anaerobic ribonucleoside-triphosphate reductase** | 1H78 | **0.008*** | -0.0058 | **0.091** | **0.053** | **0.54** | 0.45 | ●○●●●○ |
| **Multifunctional 2-oxoglutarate metabolism enzyme** | 2Y0P | **0.0056*** | **0.0081*** | 0 | 0.046 | 0.47 | **0.59** | ●●○○○● |
| **UDP-glucose 6-dehydrogenase** | 3PTZ | **0.0028*** | -0.16 | **0.14** | **0.063** | 0.46 | 0.43 | ●○●●○○ |
| **Kinesin-like protein KIF11** | 4BBG | **0.0021*** | -0.047 | **0.071** | 0.047 | **0.61** | **0.53** | ●○●○●● |
| **HD domain protein** | 4LRL | **0.0001*** | -0.018 | 0 | **0.067** | 0.48 | **0.64** | ●○○●○● |
| **Glycogen phosphorylase, muscle form** | 2IEG | -0.0035 | -0.015 | 0 | 0.034 | **0.62** | **0.68** | ○○○○●● |
| **Glucose-1-phosphate thymidylyltransferase** | 1MP3 | -0.01 | **0.1*** | **0.071** | **0.11** | **0.52** | **0.67** | ○●●●●● |
| **Cyclin-dependent kinase 2** | 3PXF | -0.012 | -0.079 | **0.056** | 0.032 | **0.57** | 0.46 | ○○●○●○ |
| **Glucose-1-phosphate thymidylyltransferase** | 4HO6 | -0.013 | **0.0016*** | 0 | 0.048 | **0.63** | **0.54** | ○●○○●● |
| **Myosin-2 heavy chain** | 2JHR | -0.014 | **0.046*** | 0 | **0.067** | **0.54** | **0.59** | ○●○●●● |
| **Pyruvate kinase 1** | 4HYW | -0.017 | -0.012 | 0 | 0.035 | 0.41 | 0.43 | ○○○○○○ |
| **N-acetylglutamate kinase / N-acetylglutamate synthase** | 4KZT | -0.022 | **0.042*** | 0 | **0.053** | 0.26 | **0.61** | ○●○●○● |
| **Myosin-2 heavy chain** | 3BZ7 | -0.024 | -0.037 | **0.12** | **0.061** | **0.69** | **0.64** | ○○●●●● |
| **Glutamate racemase** | 2VVT | -0.029 | **0.046*** | **0.17** | **0.066** | **0.53** | **0.59** | ○●●●●● |
| **Glutamine--fructose-6-phosphate** | 2PUV | -0.031 | **0.11*** | **0.077** | **0.064** | **0.6** | **0.7** | ○●●●●● |

| | | | | | | | | |
|---|---|---|---|---|---|---|---|---|
| **aminotransferase** | | | | | | | | |
| **Phospho-2-dehydro-3-deoxyheptonate aldolase AroG** | 3KGF_1 | -0.033 | -0.066 | **0.091** | 0.0078 | 0.48 | **0.53** | ○○●○○● |
| **Prostaglandin G/H synthase 2** | 3QH0 | -0.033 | -0.1 | 0 | 0.0047 | 0.43 | 0.45 | ○○○○○○ |
| **Hemoglobin subunit alpha** | 1IWH | -0.035 | -0.13 | 0 | 0.034 | **0.86** | 0.47 | ○○○○●○ |
| **Kinesin-like protein KIF11** | 3ZCW | -0.037 | **0.072*** | 0 | 0.035 | **0.62** | **0.62** | ○●○○●● |
| **NAD(P)-dependent glyceraldehyde-3-phosphate dehydrogenase** | 1UXV | -0.037 | -0.08 | 0 | 0.027 | 0.45 | 0.44 | ○○○○○○ |
| **Phospho-2-dehydro-3-deoxyheptonate aldolase, tyrosine-inhibited** | 1OF6 | -0.04 | -0.039 | **0.077** | 0.04 | 0.43 | **0.54** | ○○●○○● |
| **Pyruvate kinase 1** | 1A3W | -0.04 | **0.052*** | **0.13** | **0.07** | 0.38 | 0.49 | ○●●●○○ |
| **Phospho-2-dehydro-3-deoxyheptonate aldolase, Phe-sensitive** | 1KFL | -0.041 | -0.054 | 0 | 0.041 | **0.54** | **0.59** | ○○○○●● |
| **Ornithine decarboxylase** | 1NJJ | -0.043 | **0.079*** | 0 | **0.059** | **0.5** | **0.64** | ○●○●●● |
| **Glutamate racemase** | 4B1F | -0.043 | -0.025 | **0.14** | 0.014 | **0.5** | **0.59** | ○○●○●● |
| **Glutamate dehydrogenase 1, mitochondrial** | 3MW9 | -0.045 | **0.0086*** | 0 | 0.045 | 0.4 | **0.64** | ○●○○○● |
| **NAD-dependent malic enzyme, mitochondrial** | 1GZ3 | -0.049 | -0.059 | 0 | 0.038 | **0.51** | **0.65** | ○○○○●● |
| **Glycogen phosphorylase, muscle form** | 4MRA | -0.05 | -0.16 | **0.11** | 0.035 | **0.57** | 0.49 | ○○●○●○ |
| **Glutamate receptor 3** | 3LSW | -0.053 | -0.056 | **0.14** | **0.058** | **0.57** | 0.47 | ○○●●●○ |
| **Glutaminase kidney isoform, mitochondrial** | 4JKT | -0.057 | -0.072 | 0 | 0.0097 | 0.31 | **0.56** | ○○○○○● |
| **Acetylglutamate kinase, chloroplastic** | 2RD5 | -0.077 | -0.068 | 0 | 0.01 | 0.46 | **0.55** | ○○○○○● |

| | | | | | | | | |
|---|---|---|---|---|---|---|---|---|
| **Glutamate racemase** | 2W4I | -0.09 | -0.052 | 0 | 0.04 | 0.48 | **0.56** | ○○○○○● |
| **Prephenate dehydratase** | 3MWB | -0.098 | -0.12 | 0 | 0.014 | 0.49 | **0.5** | ○○○○○● |
| **Glutamate dehydrogenase 1, mitochondrial** | 3ETE_2 | -0.12 | -0.096 | **0.091** | 0.047 | 0.29 | **0.54** | ○○●○○● |
| **Pyruvate kinase PKLR** | 2VGI | -0.14 | -0.033 | 0 | 0.018 | 0.47 | **0.65** | ○○○○○● |
| **2-dehydro-3-deoxyphosphoheptonate aldolase** | 4GRS | -0.16 | -0.13 | 0 | 0.011 | 0.41 | **0.53** | ○○○○○● |
| **UDP-glucose 6-dehydrogenase** | 3PJG | -0.16 | -0.11 | 0 | 0.043 | **0.52** | 0.42 | ○○○○●○ |
| **Glutaminase kidney isoform, mitochondrial** | 3UO9 | -0.17 | -0.091 | 0 | 0.011 | 0.33 | **0.6** | ○○○○○○ |
| **Pyruvate kinase PKM** | 3H6O | -0.19 | -0.13 | 0 | 0.045 | 0.44 | **0.53** | ○○○○○● |
| **Pyruvate kinase** | 3HQP | -0.19 | -0.2 | 0 | 0.02 | 0.43 | 0.45 | ○○○○○○ |
| **Ribonucleoside-diphosphate reductase 1 subunit alpha** | 3R1R | -0.21 | -0.19 | 0 | 0 | 0.27 | 0.39 | ○○○○○○ |
| **Glycogen phosphorylase, muscle form** | 3BCR | -0.24 | -0.06 | 0 | 0.029 | 0.41 | **0.67** | ○○○○○● |
| **Amino-acid acetyltransferase** | 3D2P | -0.27 | -0.19 | 0 | 0.0075 | 0.17 | 0.32 | ○○○○○○ |
| **D-3-phosphoglycerate dehydrogenase** | 2PA3 | -0.29 | -0.17 | 0 | 0 | 0.039 | 0.082 | ○○○○○○ |
| **Sulfate adenylyltransferase** | 1M8P | -0.34 | -0.31 | 0 | 0 | 0.21 | 0.24 | ○○○○○○ |
| **ATP phosphoribosyltransferase** | 2VD3 | -0.4 | -0.27 | 0 | 0 | 0.12 | 0.29 | ○○○○○○ |
| **Glycogen phosphorylase, muscle form** | 1Z8D_2 | -0.43 | -0.27 | 0 | 0 | 0.15 | 0.45 | ○○○○○○ |
| **D-3-phosphoglycerate dehydrogenase** | 3DC2 | -0.43 | -0.41 | 0 | 0 | 0.0075 | 0.02 | ○○○○○○ |

| | | | | | | | | |
|---|---|---|---|---|---|---|---|---|
| **4-hydroxy-tetrahydrodipicolinate synthase** | 2ATS | -0.45 | -0.2 | 0 | 0 | 0.05 | 0.29 | ○○○○○○ |

* Results in columns that are shaded in grey have been used in Ref. [1] and are included here for complete and detailed analysis.

[1] S. F. Mersmann *et al.*, "ProteinLens: a web-based application for the analysis of allosteric signalling on atomistic graphs of biomolecules," *Nucleic Acids Res.*, May 2021.

**Table S4: Allosteric site quantile scores of proteins in Table S2 (without the presence of allosteric ligands in the structures).** The results from six statistical scores described in Methods. Average site residue and bond quantile scores are compared with those of 1000 surrogate sites of the same size. The difference is shown in bold if it is above 0 and starred if it is above the 95% confidence interval. The proportion of residues/bonds with $p_{R/b,\mathrm{allo}} > 0.95$ and the average reference quantile score $\overline{p^{\mathrm{ref}}_{R/b,\mathrm{allo}}}$ are shown in bold if they are above the expected values of 0.05 and 0.5 respectively.

| Protein | PDB | $\overline{p_{R,\mathrm{allo}}} - \langle\overline{p_{R,\mathrm{site}}}\rangle_{\mathrm{surr}}$ | $\overline{p_{b,\mathrm{allo}}} - \langle\overline{p_{b,\mathrm{site}}}\rangle_{\mathrm{surr}}$ | $P(p_{R,\mathrm{allo}} > 0.95)$ | $P(p_{b,\mathrm{allo}} > 0.95)$ | $\overline{p^{\mathrm{ref}}_{R,\mathrm{allo}}}$ | $\overline{p^{\mathrm{ref}}_{b,\mathrm{allo}}}$ | Summary |
|---|---|---|---|---|---|---|---|---|
| **Cytochrome P450 3A4** | 1W0F | **0.35*** | **0.084*** | **0.17** | 0.011 | **0.69** | **0.56** | ●●●○●● |
| **Fructose-1,6-bisphosphatase class 1** | 2Q8M_2 | **0.34*** | **0.18*** | **0.29** | **0.13** | **0.61** | **0.52** | ●●●●●● |
| **Tryptophan 2,3-dioxygenase** | 2NW8 | **0.33*** | **0.13*** | **0.43** | **0.16** | **0.83** | **0.62** | ●●●●●● |
| **Parathion hydrolase** | 1QW7 | **0.31*** | -0.12 | 0 | 0.029 | **0.77** | **0.51** | ●○○○●● |
| **Glutamate dehydrogenase 1, mitochondrial** | 3ETG | **0.3*** | **0.16*** | 0 | **0.094** | **0.64** | **0.75** | ●●○●●● |
| **Copper-containing nitrite reductase** | 1ZDS | **0.28*** | **0.13*** | **0.22** | **0.15** | **0.81** | **0.78** | ●●●●●● |
| **Aspartate carbamoyltransferase regulatory chain** | 2BE9 | **0.26*** | **0.15*** | **0.091** | **0.11** | 0.091 | 0.084 | ●●●●○○ |
| **Androgen receptor** | 2QPY | **0.25*** | **0.22*** | **0.3** | **0.11** | **0.73** | **0.54** | ●●●●●● |
| **Tyrosine-protein kinase ABL1** | 3K5V | **0.22*** | **0.14*** | **0.12** | **0.052** | **0.66** | **0.63** | ●●●●●● |
| **Glutamate receptor 3** | 3LSX | **0.22*** | **0.041*** | 0 | **0.22** | **0.7** | 0.45 | ●●○●●○ |
| **Fructose-1,6-bisphosphatase isozyme 2** | 3IFA | **0.21*** | **0.079*** | **0.14** | **0.11** | **0.56** | **0.59** | ●●●●●● |
| **Lysine-sensitive aspartokinase 3** | 2J0X | **0.19*** | **0.16*** | **0.36** | **0.074** | **0.56** | **0.6** | ●●●●●● |
| **Isocitrate dehydrogenase** | 3EPS | **0.17*** | **0.19*** | **0.15** | **0.052** | **0.66** | **0.79** | ●●●●●● |

| | | | | | | | | |
|---|---|---|---|---|---|---|---|---|
| **kinase/phosphatase** | | | | | | | | |
| **Pyruvate kinase PKM** | 4G1N | **0.17*** | **0.03*** | 0 | 0.036 | **0.64** | **0.72** | ●●○○●● |
| **Serum albumin** | 3LU6 | **0.17*** | **0.096*** | **0.17** | **0.092** | **0.7** | **0.63** | ●●●●●● |
| **Pyruvate kinase PKM** | 3N25 | **0.16*** | **0.084*** | 0 | **0.097** | **0.57** | **0.74** | ●●○●●● |
| **Anthranilate synthase component 1** | 1I7S | **0.16*** | **0.091*** | 0 | 0.045 | **0.55** | **0.6** | ●●○○●● |
| **Glutamate dehydrogenase 1, mitochondrial** | 3ETE_1 | **0.15*** | **0.079*** | 0 | 0.038 | **0.59** | **0.73** | ●●○○●● |
| **Pyruvate dehydrogenase kinase isozyme 2** | 2BU8 | **0.15*** | **0.084*** | 0 | **0.077** | **0.83** | **0.65** | ●●○●●● |
| **Glycogen phosphorylase, liver form** | 1EM6 | **0.14*** | **0.083*** | **0.13** | **0.066** | **0.73** | **0.73** | ●●●●●● |
| **ATP phosphoribosyltransferase** | 1NH8 | **0.14*** | **0.1*** | 0 | **0.073** | 0.087 | 0.094 | ●●○●○○ |
| **Casein kinase II subunit alpha** | 3H30 | **0.13*** | **0.091*** | 0 | **0.079** | **0.73** | **0.75** | ●●○●●● |
| **Glutamate racemase** | 3HFR | **0.13*** | **0.033*** | **0.2** | 0.042 | **0.62** | **0.62** | ●●●○●● |
| **Fructose-1,6-bisphosphatase class 1** | 2Q8M_1 | **0.13*** | -0.049 | 0 | 0.017 | **0.58** | 0.48 | ●○○○●○ |
| **Indole-3-pyruvate decarboxylase** | 2Q5O | **0.12*** | **0.056*** | **0.17** | **0.1** | **0.61** | **0.7** | ●●●●●● |
| **Carbamoyl-phosphate synthase large chain** | 1CE8_1 | **0.12*** | **0.01*** | **0.091** | **0.13** | **0.53** | **0.62** | ●●●●●● |
| **Acetyl-CoA carboxylase** | 1W96 | **0.12*** | -0.05 | 0 | **0.061** | **0.65** | **0.6** | ●○○●●● |
| **Leukotriene A-4 hydrolase** | 3FUD | **0.11*** | -0.041 | 0 | **0.11** | **0.52** | 0.42 | ●○○●●○ |
| **Serum albumin** | 2BXA | **0.1*** | **0.039*** | **0.071** | **0.06** | **0.65** | **0.54** | ●●●●●● |
| **ATP-dependent 6-phosphofructokinase** | 4PFK | **0.1*** | **0.063*** | 0 | **0.11** | **0.51** | 0.38 | ●●○●●○ |
| **Pyruvate kinase PKM** | 4B2D | **0.095*** | **0.11*** | **0.091** | **0.14** | **0.54** | **0.75** | ●●●●●● |
| **Tyrosine-protein** | 1T49 | **0.091*** | -0.1 | **0.17** | 0.038 | **0.7** | 0.38 | ●○●○●○ |

| | | | | | | | | |
|---|---|---|---|---|---|---|---|---|
| **phosphatase non-receptor type 1** | | | | | | | | |
| **Androgen receptor** | 2YLO | **0.08*** | **0.025*** | 0 | 0.016 | **0.58** | 0.42 | ●●○○●○ |
| **Glycogen phosphorylase, muscle form** | 1Z8D_1 | **0.079*** | -0.087 | 0 | 0.034 | **0.55** | 0.39 | ●○○○●○ |
| **Carbamoyl-phosphate synthase large chain** | 1CE8_2 | **0.073*** | **0.067*** | **0.11** | **0.14** | **0.53** | **0.68** | ●●●●●● |
| **Ribonucleoside-diphosphate reductase 1 subunit alpha** | 2R1R | **0.073*** | -0.016 | **0.11** | **0.095** | 0.49 | **0.59** | ●○●●○● |
| **Glycogen phosphorylase, liver form** | 3CEH | **0.071*** | -0.11 | 0 | **0.058** | **0.68** | **0.57** | ●○○●●● |
| **Glucose-1-phosphate thymidylyltransferase 1** | 1H5S | **0.069*** | **0.1*** | **0.21** | **0.097** | **0.53** | **0.73** | ●●●●●● |
| **Androgen receptor** | 2YHD | **0.067*** | **0.022*** | **0.12** | **0.05** | **0.69** | **0.52** | ●●●●●● |
| **L-asparaginase 1** | 2HIM | **0.066*** | **0.15*** | 0 | **0.11** | **0.54** | **0.76** | ●●○●●● |
| **5'-AMP-activated protein kinase catalytic subunit alpha-2** | 4CFE | **0.065*** | **0.11*** | **0.25** | **0.097** | **0.59** | **0.68** | ●●●●●● |
| **Seminal ribonuclease** | 11BG | **0.06*** | **0.00078*** | 0 | **0.077** | **0.73** | 0.48 | ●●○●●● |
| **Tyrosine-protein kinase ABL1** | 3PYY | **0.058*** | **0.056*** | **0.083** | 0.043 | **0.51** | **0.55** | ●●●○●● |
| **Phospho-2-dehydro-3-deoxyheptonate aldolase** | 3PG9 | **0.057*** | **0.049*** | **0.13** | **0.097** | **0.64** | **0.7** | ●●●●●● |
| **Glucose-1-phosphate thymidylyltransferase** | 1LVW | **0.044*** | -0.022 | 0 | **0.069** | **0.5** | **0.62** | ●○○●●● |
| **Toxin A** | 3HO6 | **0.043*** | -0.018 | **0.071** | **0.088** | **0.68** | **0.57** | ●○●●●● |
| **Prephenate dehydratase** | 2QMX | **0.042*** | -0.01 | 0 | 0.049 | **0.67** | **0.65** | ●○○○●● |
| **Isocitrate dehydrogenase [NADP], mitochondrial** | 4JA8 | **0.042*** | -0.064 | 0.043 | 0.0033 | 0.49 | 0.48 | ●○○○○○ |

| | | | | | | | | |
|---|---|---|---|---|---|---|---|---|
| **Hemoglobin subunit beta** | 2D60 | **0.038*** | **0.017*** | **0.1** | 0.043 | **0.68** | **0.62** | ●●●○●● |
| **Fructose-1,6-bisphosphatase 1** | 1FRP | **0.038*** | **0.014*** | **0.067** | **0.057** | **0.56** | **0.52** | ●●●●●● |
| **Chorismate mutase** | 3CSM | **0.029*** | **0.057*** | 0 | 0.015 | **0.57** | **0.53** | ●●○○●● |
| **Mitogen-activated protein kinase 8** | 3O2M | **0.028*** | -0.0066 | 0 | 0.0055 | **0.6** | **0.56** | ●●●●●● |
| **Glutamate receptor 3** | 3M3F | **0.025*** | **0.039*** | **0.17** | **0.054** | **0.64** | 0.49 | ●●●●●○ |
| **Glycogen phosphorylase, muscle form** | 3E3N | **0.02*** | -0.1 | 0 | 0.029 | **0.59** | **0.56** | ●○○○●● |
| **Phospho-2-dehydro-3-deoxyheptonate aldolase AroG** | 3KGF_2 | **0.0054*** | **0.07*** | **0.2** | **0.056** | 0.49 | **0.62** | ●●●●○●● |
| **Glucosamine-6-phosphate isomerase 1** | 1NE7 | **0.0031*** | **0.028*** | 0 | **0.07** | **0.5** | **0.64** | ●●○●●● |
| **Uridylate kinase** | 2V4Y | -0.00068 | **0.012*** | 0 | **0.069** | 0.49 | **0.63** | ○●○●○● |
| **HD domain protein** | 3IRH | -0.0015 | -0.019 | **0.14** | 0.047 | 0.46 | **0.62** | ○○●○○● |
| **Uracil phosphoribosyltransferase** | 1JLR | -0.0044 | -0.02 | **0.077** | **0.063** | **0.7** | **0.63** | ○○●●●● |
| **Probable aspartokinase** | 3C1N | -0.0056 | -0.13 | 0 | 0 | 0.35 | 0.46 | ○○○○○○ |
| **Glucose-1-phosphate thymidylyltransferase** | 1MP3 | -0.019 | **0.099*** | **0.071** | **0.11** | **0.52** | **0.67** | ○●●●●● |
| **Uridylate kinase** | 3EK5 | -0.023 | -0.016 | **0.067** | **0.083** | **0.56** | **0.6** | ○○●●●● |
| **Kinesin-like protein KIF11** | 4BBG | -0.026 | -0.1 | **0.071** | **0.064** | **0.6** | 0.43 | ○○●●○○ |
| **Myosin-2 heavy chain** | 2JHR | -0.029 | **0.052*** | 0 | **0.084** | **0.53** | **0.59** | ○●○●●● |
| **Hemoglobin subunit alpha** | 1IWH | -0.03 | -0.15 | 0 | **0.075** | **0.86** | 0.43 | ○○○●●○ |
| **Anaerobic ribonucleoside-triphosphate reductase** | 1H78 | -0.03 | -0.035 | 0 | 0.037 | **0.51** | 0.43 | ○○○○●○ |
| **Response regulator PleD** | 1W25 | -0.032 | **0.062*** | 0 | **0.079** | **0.66** | **0.72** | ○●○●●● |
| **Glycogen phosphorylase, muscle form** | 2IEG | -0.035 | -0.012 | 0 | 0.03 | **0.59** | **0.68** | ○○○○●● |

| | | | | | | | | |
|---|---|---|---|---|---|---|---|---|
| **N-acetylglutamate kinase / N-acetylglutamate synthase** | 4KZT | -0.042 | **0.01*** | 0 | 0.043 | 0.24 | **0.56** | ○●○○○● |
| **Glucose-1-phosphate thymidylyltransferase** | 1G3L | -0.043 | **0.044*** | 0 | **0.064** | **0.51** | **0.7** | ○●○●●● |
| **ATP-dependent 6-phosphofructokinase isozyme 1** | 1PFK | -0.051 | -0.14 | 0 | 0.028 | **0.51** | 0.42 | ○○○○●○ |
| **Phospho-2-dehydro-3-deoxyheptonate aldolase AroG** | 3KGF_1 | -0.053 | -0.052 | **0.091** | 0.0087 | 0.46 | **0.53** | ○○●○○● |
| **Prostaglandin G/H synthase 2** | 3QH0 | -0.053 | -0.12 | 0 | 0.011 | 0.41 | 0.43 | ○○○○○○ |
| **NAD(P)-dependent glyceraldehyde-3-phosphate dehydrogenase** | 1UXV | -0.057 | -0.08 | 0 | 0.029 | 0.45 | 0.45 | ○○○○○○ |
| **Cytosolic purine 5'-nucleotidase** | 2XJC | -0.058 | **0.019*** | 0 | **0.058** | **0.54** | **0.56** | ○●○●●● |
| **Glutamate racemase** | 2VVT | -0.063 | **0.016*** | **0.17** | **0.085** | **0.51** | **0.57** | ○●●●●● |
| **Kinesin-like protein KIF11** | 3ZCW | -0.064 | **0.016*** | 0 | 0.034 | **0.61** | **0.55** | ○●○○●● |
| **Glutamine--fructose-6-phosphate aminotransferase** | 2PUV | -0.066 | -0.024 | **0.077** | 0.044 | **0.56** | **0.58** | ○○●○●● |
| **Glutamate receptor 3** | 3LSW | -0.066 | **0.015*** | **0.14** | **0.078** | **0.55** | **0.55** | ○●●●●● |
| **Phospho-2-dehydro-3-deoxyheptonate aldolase, Phe-sensitive** | 1KFL | -0.067 | -0.046 | 0 | 0.046 | **0.51** | **0.59** | ○○○○●● |
| **Phospho-2-dehydro-3-deoxyheptonate aldolase, tyrosine-inhibited** | 1OF6 | -0.068 | -0.038 | **0.077** | 0.045 | 0.41 | **0.53** | ○○●○○● |
| **NAD-dependent malic** | 1GZ3 | -0.071 | -0.11 | 0 | 0.049 | 0.49 | **0.6** | ○○○○○● |

| | | | | | | | | |
|---|---|---|---|---|---|---|---|---|
| **enzyme, mitochondrial** | | | | | | | | |
| **Glucose-1-phosphate thymidylyltransferase** | 4HO6 | -0.073 | -0.0046 | 0 | 0.034 | **0.53** | **0.53** | ○○○○●● |
| **Glutamate dehydrogenase 1, mitochondrial** | 3MW9 | -0.077 | -0.0093 | 0 | 0.049 | 0.37 | **0.63** | ○○○○○● |
| **Pyruvate kinase 1** | 1A3W | -0.081 | **0.077*** | **0.067** | **0.063** | 0.34 | **0.51** | ○●●●○● |
| **Mitogen-activated protein kinase 14** | 4E6C | -0.082 | -0.051 | 0 | **0.1** | 0.24 | 0.24 | ○○○●○○ |
| **Glycogen phosphorylase, muscle form** | 4MRA | -0.083 | -0.24 | 0 | 0.034 | **0.53** | 0.41 | ○○○○●○ |
| **UDP-glucose 6-dehydrogenase** | 3PTZ | -0.087 | -0.2 | **0.091** | 0.037 | 0.38 | 0.4 | ○○●○○○ |
| **Acetylglutamate kinase, chloroplastic** | 2RD5 | -0.089 | -0.076 | 0 | 0.011 | 0.45 | **0.54** | ○○○○○● |
| **Pyruvate kinase 1** | 4HYW | -0.089 | -0.015 | 0 | 0.025 | 0.35 | 0.43 | ○○○○○○ |
| **Myosin-2 heavy chain** | 3BZ7 | -0.096 | -0.039 | **0.12** | **0.061** | **0.63** | **0.61** | ○○●●●● |
| **Multifunctional 2-oxoglutarate metabolism enzyme** | 2Y0P | -0.099 | -0.052 | 0 | 0.039 | 0.4 | **0.54** | ○○○○○● |
| **Putative deoxycytidylate deaminase** | 2HVW | -0.099 | -0.19 | 0 | **0.11** | **0.62** | 0.36 | ○○○●●○ |
| **HD domain protein** | 4LRL | -0.11 | -0.052 | 0 | 0.041 | 0.39 | **0.62** | ○○○○○● |
| **Glutaminase kidney isoform, mitochondrial** | 4JKT | -0.11 | -0.061 | 0 | 0.014 | 0.27 | **0.56** | ○○○○○● |
| **Ornithine decarboxylase** | 1NJJ | -0.12 | **0.023*** | 0 | **0.068** | 0.43 | **0.58** | ○●○●○● |
| **Glutamate dehydrogenase 1, mitochondrial** | 3ETE_2 | -0.12 | -0.094 | **0.091** | 0.035 | 0.29 | **0.54** | ○○●○○● |
| **Ribose-phosphate pyrophosphokinase** | 1DKU | -0.13 | -0.012 | 0 | **0.086** | 0.48 | **0.56** | ○○○●○● |
| **Prephenate dehydratase** | 3MWB | -0.13 | -0.11 | 0 | 0.028 | 0.46 | 0.48 | ○○○○○○ |

| | | | | | | | | |
|---|---|---|---|---|---|---|---|---|
| **Cyclin-dependent kinase 2** | 3PXF | -0.13 | -0.14 | 0 | 0.027 | **0.5** | 0.41 | ○○○○●○ |
| **Glutamate racemase** | 4B1F | -0.15 | -0.15 | 0 | 0.014 | 0.46 | 0.48 | ○○○○○○ |
| **Pyruvate kinase PKLR** | 2VGI | -0.19 | -0.028 | 0 | 0.023 | 0.4 | **0.66** | ○○○○○● |
| **Glutaminase kidney isoform, mitochondrial** | 3UO9 | -0.21 | -0.13 | 0 | 0.015 | 0.29 | **0.53** | ○○○○○● |
| **2-dehydro-3-deoxyphosphoheptonate aldolase** | 4GRS | -0.22 | -0.16 | 0 | 0 | 0.36 | **0.51** | ○○○○○● |
| **Pyruvate kinase** | 3HQP | -0.24 | -0.22 | 0 | 0.016 | 0.38 | 0.44 | ○○○○○○ |
| **Pyruvate kinase PKM** | 3H6O | -0.27 | -0.17 | 0 | 0.029 | 0.33 | 0.49 | ○○○○○○ |
| **Amino-acid acetyltransferase** | 3D2P | -0.28 | -0.17 | 0 | 0.0083 | 0.17 | 0.34 | ○○○○○○ |
| **Glycogen phosphorylase, muscle form** | 3BCR | -0.29 | -0.21 | 0 | 0.028 | 0.36 | **0.55** | ○○○○○● |
| **Glutamate racemase** | 2W4I | -0.29 | -0.23 | 0 | 0.022 | 0.32 | 0.43 | ○○○○○○ |
| **UDP-glucose 6-dehydrogenase** | 3PJG | -0.29 | -0.23 | 0 | 0.0079 | 0.38 | 0.27 | ○○○○○○ |
| **D-3-phosphoglycerate dehydrogenase** | 2PA3 | -0.3 | -0.15 | 0 | 0 | 0.039 | 0.089 | ○○○○○○ |
| **Ribonucleoside-diphosphate reductase 1 subunit alpha** | 3R1R | -0.32 | -0.24 | 0 | 0 | 0.18 | 0.35 | ○○○○○○ |
| **Sulfate adenylyltransferase** | 1M8P | -0.36 | -0.32 | 0 | 0 | 0.19 | 0.24 | ○○○○○○ |
| **ATP phosphoribosyltransferase** | 2VD3 | -0.39 | -0.25 | 0 | 0 | 0.12 | 0.32 | ○○○○○○ |
| **D-3-phosphoglycerate dehydrogenase** | 3DC2 | -0.43 | -0.41 | 0 | 0 | 0.0062 | 0.017 | ○○○○○○ |
| **Glycogen phosphorylase, muscle form** | 1Z8D_2 | -0.44 | -0.27 | 0 | 0 | 0.15 | 0.47 | ○○○○○○ |

| | | | | | | | | |
|---|---|---|---|---|---|---|---|---|
| **4-hydroxy-tetrahydrodipicolinate synthase** | 2ATS | -0.47 | -0.24 | 0 | 0 | 0.05 | 0.25 | ○○○○○○ |
| **Hemoglobin subunit beta** | 1B86 | -0.48 | -0.35 | 0 | 0 | 0.075 | 0.22 | ○○○○○○ |

**Table S5: Protein structure used from the CASBench database.** Orthosteric and allosteric ligands and site residues can be found directly in the CASBench database.

| cas number | PDB |
|---|---|
| **cas0001** | 1nxe, 1nxg, 1owc, 4g6b, 4jad, 4jae, 4jaf, 4jag |
| **cas0002** | 3i1y, 3i28, 3koo, 3otq, 5ahx, 5ai4, 5ai5, 5aia, 5ak4, 5ak5, 5ake, 5akh, 5akx, 5aky, 5ald, 5alf, 5alh, 5alm, 5aln, 5alo, 5alt, 5alu, 5alv, 5alw, 5aly, 5am0, 5am4, 5am5 |
| **cas0003** | 3ion, 3iop, 3rwp, 4rqk, 4rqv, 4rrv, 4xx9 |
| **cas0004** | 4ey5, 5hf6, 5hf8, 5hf9, 5hfa |
| **cas0010** | 1i2d |
| **cas0011** | 2ewn |
| **cas0015** | 2ym4, 2ym8, 4fst, 4fsy, 4ft3, 4ft7, 4fta, 4ftn, 4fto, 4ftr, 4ftu, 4gh2 |
| **cas0016** | 3csm, 4csm |
| **cas0021** | 3l9h, 4zhi |
| **cas0024** | 4ald |
| **cas0027** | 1cza, 1dgk |
| **cas0028** | 1lld, 1lth |
| **cas0029** | 1ldn |
| **cas0030** | 2him, 2p2d |
| **cas0039** | 3ddn |
| **cas0040** | 1psd, 1yba, 2p9c, 2p9e, 2p9g, 2pa3 |
| **cas0047** | 4r1r |
| **cas0050** | 1xtu, 1xtv |
| **cas0051** | 11ba, 11bg, 1bsr, 1n3z, 1r3m, 1r5c, 1tq9, 3bcm, 3djo, 3djp, 3djq, 3djv, 3djx, 4n4c |
| **cas0052** | 2z60 |
| **cas0054** | 2jfx, 2jfy, 2jfz, 4b1f |
| **cas0056** | 3hmi |
| **cas0060** | 1boz, 1dhf, 1dlr, 1dls, 1drf, 1hfp, 1hfq, 1hfr, 1kms, 1mvs, 1mvt, 1ohj, 1ohk, 1pd8, 1pd9, 1s3u, 1s3v, 1s3w, 1u71, 1u72, 2c2s, 2c2t, 2dhf, |

| | |
|---|---|
| | 2w3a, 2w3m, 3f8y, 3f8z, 3fs6, 3ghc, 3ghw, 3gi2, 3gyf, 3l3r, 3n0h, 3ntz, 3nu0, 3nxr, 3nxt, 3nxv, 3nxx, 3nxy, 3nzd, 3oaf, 3s3v, 3s7a, 4ddr, 4g95, 4kd7, 4keb, 4m6k, 4m6l, 4qjc, 5hpb, 5hqy, 5hqz, 5hsr, 5hsu, 5ht4, 5ht5, 5hui, 5hvb, 5hve |
| **cas0061** | 1dre, 1ra2, 1ra3, 1ra8, 1rb2, 1rb3, 1rc4, 1rd7, 1re7, 1rg7, 1rh3, 1rx4, 1rx5, 1rx6, 1rx7, 3dau, 4ej1, 4fhb, 4i13, 4i1n, 4kjj, 4kjl, 4p3r, 4qle, 4qlg, 4x5f, 4x5g, 4x5h, 4x5i, 4x5j, 5cc9, 5ccc, 7dfr |
| **cas0067** | 2psq, 3ril |
| **cas0070** | 1ibc, 1ice, 1rwm, 1rwn, 1rwo, 1rwv, 2h4w, 2h4y, 2h51, 2h54, 2hbq, 2hbr, 2hby, 2hbz, 3d6f, 3d6h, 3d6m |
| **cas0071** | 1i4o |
| **cas0074** | 1ibv, 1ibw, 1pya |
| **cas0079** | 1bzc, 1bzj, 1c83, 1c84, 1c85, 1c86, 1c87, 1c88, 1ecv, 1g1g, 1g7g, 1gfy, 1kak, 1kav, 1l8g, 1nwe, 1ptt, 1ptu, 1ptv, 1q1m, 1xbo, 2azr, 2b07, 2bge, 2cm7, 2cma, 2h4g, 2h4k, 2hb1, 2nt7, 2nta, 2qbp, 2qbq, 2qbr, 2qbs, 2veu, 2vey, 2zn7, 4i8n |
| **cas0080** | 1i00, 1juj, 3ob7, 5hs3, 5x5q, 5x67 |
| **cas0085** | 1fuo, 1fup, 1fuq, 1kq7 |
| **cas0086** | 1gos, 1oj9, 1oja, 1ojc, 1s2q, 1s2y, 1s3b, 1s3e, 2bk3, 2bk4, 2bk5, 2byb, 2c64, 2c65, 2c66, 2c67, 2c70, 2c72, 2c73, 2c75, 2c76, 2v5z, 2v60, 2v61, 2vrl, 2vrm, 2vz2, 2xcg, 2xfn, 2xfo, 2xfp, 2xfq, 2xfu, 3po7, 3zyx, 4a79, 4a7a, 4crt, 5mrl |
| **cas0091** | 1fin, 2b54, 2cch, 4ez7 |

**Table S6: Allosteric site quantile scores of proteins in Table S5 (orthosteric ligands as the source)** The results from six statistical scores described in Methods. Average site residue and bond quantile scores are compared with those of 1000 surrogate sites of the same size. The difference is shown in bold if it is above 0 and starred if it is above the 95% confidence interval. The proportion of residues/bonds with $p_{R/b,\text{allo}} > 0.95$ and the average reference quantile score $\overline{p^{\text{ref}}_{R/b,\text{ allo}}}$ are shown in bold if they are above the expected values of 0.05 and 0.5 respectively.

| cas Number | PDB | $\overline{p_{R,\text{allo}}} - \langle\overline{p_{R,\text{site}}}\rangle_{\text{surr}}$ | $\overline{p_{b,\text{allo}}} - \langle\overline{p_{b,\text{site}}}\rangle_{\text{surr}}$ | P($p_{R,\text{allo}}$ > 0.95) | P($p_{b,\text{allo}}$ > 0.95) | $\overline{p^{\text{ref}}_{R,\text{ allo}}}$ | $\overline{p^{\text{ref}}_{b,\text{ allo}}}$ | Summary |
|---|---|---|---|---|---|---|---|---|
| cas0001 | 1nxe | -0.052 | -0.087 | 0 | 0.032 | **0.61** | **0.66** | ○○○○●● |
| cas0001 | 1nxg | -0.03 | -0.067 | 0 | 0.024 | **0.64** | **0.68** | ○○○○●● |
| cas0001 | 1owc | -0.011 | -0.04 | 0 | 0.028 | **0.63** | **0.68** | ○○○○●● |
| cas0001 | 4g6b | -0.019 | -0.056 | 0 | 0.037 | **0.64** | **0.67** | ○○○○●● |
| cas0001 | 4jad | -0.043 | -0.067 | 0 | 0.017 | **0.54** | **0.62** | ○○○○●● |
| cas0001 | 4jae | -0.026 | -0.09 | 0 | 0.033 | **0.57** | **0.63** | ○○○○●● |
| cas0001 | 4jaf | -0.018 | -0.082 | 0 | 0.026 | **0.63** | **0.64** | ○○○○●● |
| cas0001 | 4jag | -0.024 | -0.06 | 0 | 0.02 | **0.6** | **0.65** | ○○○○●● |
| cas0002 | 3i1y | -0.027 | **0.027*** | 0 | **0.064** | 0.18 | 0.19 | ○●○●○○ |
| cas0002 | 3i28 | **0.029*** | **0.016*** | **0.067** | **0.069** | 0.21 | 0.19 | ●●●●○○ |
| cas0002 | 3ion | **0.13*** | -0.072 | **0.17** | **0.073** | **0.84** | **0.6** | ●○●●●● |
| cas0002 | 3iop | **0.16*** | -0.06 | **0.11** | **0.06** | **0.85** | **0.61** | ●○●●●● |
| cas0002 | 3koo | -0.00057 | -0.015 | 0.033 | 0.032 | 0.18 | 0.17 | ○○○○○○ |
| cas0002 | 3otq | **0.021*** | **0.025*** | **0.067** | **0.074** | 0.2 | 0.21 | ●●●●○○ |
| cas0002 | 5ahx | -0.12 | -0.022 | 0 | 0.034 | **0.92** | **0.86** | ○○○○●● |
| cas0002 | 5ai4 | **0.029*** | **0.022*** | 0 | **0.061** | 0.18 | 0.17 | ●●○●○○ |
| cas0002 | 5ai5 | **0.00027*** | **0.056*** | 0.033 | **0.072** | **0.57** | **0.56** | ●●○●●● |
| cas0002 | 5aia | -0.24 | -0.17 | 0 | 0 | 0.22 | 0.27 | ○○○○○○ |
| cas0002 | 5ak4 | **0.023*** | **0.024*** | 0 | **0.061** | 0.17 | 0.15 | ●●○●○○ |
| cas0002 | 5ak5 | **0.039*** | **0.013*** | 0.033 | **0.053** | **0.6** | **0.5** | ●●○●●● |

| | | | | | | | | |
|---|---|---|---|---|---|---|---|---|
| cas0002 | 5ake | **0.024*** | **0.027*** | **0.1** | **0.071** | **0.58** | **0.52** | ●●●●●● |
| cas0002 | 5akh | **0.062*** | **0.042*** | **0.067** | **0.052** | **0.58** | **0.53** | ●●●●●● |
| cas0002 | 5akx | **0.02*** | **0.0027*** | 0 | 0.029 | **0.59** | **0.52** | ●●○○●● |
| cas0002 | 5aky | **0.033*** | **0.031*** | 0.033 | **0.08** | **0.57** | **0.52** | ●●○●●● |
| cas0002 | 5ald | **0.041*** | **0.042*** | 0.033 | **0.062** | **0.6** | **0.54** | ●●○●●● |
| cas0002 | 5alf | -0.001 | **0.033*** | 0 | **0.053** | **0.58** | **0.53** | ○●○●●● |
| cas0002 | 5alh | **0.013*** | **0.022*** | 0.033 | 0.044 | **0.57** | **0.52** | ●●○○●● |
| cas0002 | 5alm | **0.0016*** | **0.039*** | **0.1** | **0.063** | **0.6** | **0.55** | ●●●●●● |
| cas0002 | 5alo | **0.041*** | **0.026*** | 0.033 | **0.062** | 0.17 | 0.16 | ●●○●○○ |
| cas0002 | 5alu | **0.013*** | **0.027*** | **0.067** | **0.088** | 0.19 | 0.19 | ●●●●○○ |
| cas0002 | 5alv | **0.068*** | **0.027*** | **0.067** | **0.066** | 0.14 | 0.13 | ●●●●○○ |
| cas0002 | 5alw | **0.033*** | **0.022*** | 0.033 | **0.068** | 0.26 | 0.24 | ●●○●○○ |
| cas0002 | 5aly | **0.034*** | **0.025*** | 0 | 0.047 | 0.15 | 0.16 | ●●○○○○ |
| cas0002 | 5am0 | **0.046*** | **0.066*** | 0 | **0.078** | 0.18 | 0.19 | ●●○●○○ |
| cas0002 | 5am4 | **0.034*** | **0.013*** | 0.033 | **0.067** | 0.22 | 0.21 | ●●○●○○ |
| cas0002 | 5am5 | **0.052*** | **0.05*** | 0.033 | **0.089** | 0.19 | 0.19 | ●●○●○○ |
| cas0003 | 4ey5 | **0.049*** | -0.0074 | **0.083** | **0.078** | **0.57** | **0.64** | ●○●●●● |
| cas0003 | 4rqv | **0.11*** | -0.065 | **0.17** | 0.049 | **0.83** | **0.61** | ●○●○●● |
| cas0003 | 4rrv | **0.11*** | **0.0087*** | **0.11** | **0.087** | **0.83** | **0.69** | ●●●●●● |
| cas0003 | 4xx9 | **0.12*** | -0.053 | **0.11** | **0.058** | **0.87** | **0.64** | ●○●●●● |
| cas0003 | 5aln | -0.067 | -0.07 | 0 | 0.022 | 0.088 | 0.1 | ○○○○○○ |
| cas0003 | 5alt | -0.19 | -0.17 | 0 | 0 | 0.014 | 0.023 | ○○○○○○ |
| cas0003 | 5hf6 | **0.022*** | -0.056 | 0 | 0.045 | **0.6** | **0.66** | ●○○○●● |
| cas0004 | 3rwp | **0.14*** | -0.03 | **0.056** | **0.073** | **0.83** | **0.6** | ●○●●●● |
| cas0004 | 4rqk | **0.11*** | -0.094 | **0.056** | **0.057** | **0.85** | **0.58** | ●○●●●● |
| cas0004 | 5hf8 | -0.054 | -0.079 | 0 | 0.024 | **0.55** | **0.64** | ○○○○●● |

| | | | | | | | | |
|---|---|---|---|---|---|---|---|---|
| cas0004 | 5hf9 | -0.066 | -0.12 | 0 | 0.016 | **0.51** | **0.61** | ○○○○●● |
| cas0004 | 5hfa | -0.14 | -0.11 | 0 | 0.025 | 0.47 | **0.64** | ○○○○○● |
| cas0010 | 1i2d | **0.076*** | **0.014*** | **0.14** | 0.045 | **0.52** | **0.58** | ●●●○●● |
| cas0011 | 2ewn | **0.0068*** | -0.19 | 0 | 0.0067 | **0.68** | 0.44 | ●○○○●○ |
| cas0015 | 1ldn | **0.005*** | **0.085*** | 0 | 0.037 | 0.23 | **0.56** | ●●○○○● |
| cas0015 | 1lld | -0.15 | -0.068 | 0 | 0 | 0.46 | **0.57** | ○○○○○● |
| cas0015 | 1lth | -0.072 | -0.14 | 0 | 0.027 | **0.63** | **0.6** | ○○○○●● |
| cas0015 | 2him | -0.014 | **0.038*** | 0 | 0.018 | 0.41 | **0.65** | ○●○○○● |
| cas0015 | 3l9h | **0.021*** | **0.013*** | **0.13** | 0.049 | **0.61** | **0.61** | ●●●○●● |
| cas0015 | 4ald | **0.17*** | -0.045 | 0 | 0 | **0.74** | **0.65** | ●○○○●● |
| cas0015 | 4fst | -0.24 | -0.19 | 0 | 0.046 | **0.7** | **0.58** | ○○○○●● |
| cas0015 | 4fsy | -0.26 | -0.099 | 0 | 0.029 | **0.65** | **0.65** | ○○○○●● |
| cas0015 | 4ft3 | -0.25 | -0.22 | 0 | 0.033 | **0.71** | **0.55** | ○○○○●● |
| cas0015 | 4ft7 | -0.25 | -0.22 | 0 | 0.032 | **0.72** | **0.56** | ○○○○●● |
| cas0015 | 4fto | -0.24 | -0.21 | 0.048 | 0.042 | **0.71** | **0.56** | ○○○○●● |
| cas0015 | 4gh2 | -0.25 | -0.22 | 0 | 0.041 | **0.7** | **0.59** | ○○○○●● |
| cas0016 | 4ftn | -0.21 | -0.19 | 0.048 | 0.034 | **0.73** | **0.54** | ○○○○●● |
| cas0016 | 4ftr | -0.22 | -0.21 | **0.095** | 0.036 | **0.74** | **0.54** | ○○●○●● |
| cas0021 | 2ym4 | -0.16 | -0.23 | 0.048 | 0.023 | **0.75** | **0.53** | ○○○○●● |
| cas0021 | 4zhi | -0.072 | -0.049 | 0.032 | 0.021 | **0.51** | **0.54** | ○○○○●● |
| cas0024 | 4ftu | -0.24 | -0.24 | 0 | 0.041 | **0.73** | **0.53** | ○○○○●● |
| cas0027 | 2ym8 | -0.18 | -0.22 | 0 | 0.031 | **0.71** | 0.49 | ○○○○●○ |
| cas0027 | 4fta | -0.25 | -0.24 | 0.048 | 0.023 | **0.73** | **0.51** | ○○○○●● |
| cas0028 | 3csm | **0.048*** | **0.018*** | **0.056** | **0.054** | **0.6** | 0.49 | ●●●●●○ |
| cas0028 | 4csm | **0.069*** | **0.0099*** | 0.028 | **0.08** | **0.62** | 0.47 | ●●○●●○ |
| cas0029 | 1cza | **0.26*** | **0.039*** | **0.067** | **0.1** | 0.47 | 0.31 | ●●●●○○ |

| | | | | | | | | |
|---|---|---|---|---|---|---|---|---|
| cas0030 | 1dgk | **0.27*** | **0.073*** | **0.067** | **0.11** | 0.37 | 0.26 | ●●●●○○ |
| cas0030 | 2p2d | -0.12 | **0.0063*** | 0 | 0.0095 | 0.33 | **0.6** | ○●○○○● |
| cas0039 | 3ddn | -0.24 | -0.14 | 0 | 0.0047 | **0.8** | **0.79** | ○○○○●● |
| cas0040 | 1psd | -0.071 | **0.063*** | 0 | **0.051** | **0.59** | **0.71** | ○●○●●● |
| cas0040 | 1yba | **0.017*** | **0.095*** | **0.067** | **0.11** | 0.44 | **0.59** | ●●●●○● |
| cas0040 | 2p9c | -0.035 | **0.08*** | 0 | **0.062** | **0.54** | **0.67** | ○●○●●● |
| cas0040 | 2p9e | -0.013 | **0.096*** | **0.067** | **0.1** | 0.44 | **0.61** | ○●●●○● |
| cas0040 | 2p9g | **0.027*** | **0.16*** | **0.17** | **0.14** | **0.68** | **0.83** | ●●●●●● |
| cas0040 | 2pa3 | -0.0033 | **0.19*** | **0.13** | **0.11** | **0.66** | **0.85** | ○●●●●● |
| cas0047 | 1xtv | -0.034 | -0.036 | 0.01 | 0.026 | 0.49 | **0.56** | ○○○○○● |
| cas0050 | 1bsr | **0.1*** | -0.038 | **0.091** | **0.06** | **0.81** | **0.51** | ●○●●●● |
| cas0050 | 1n3z | -0.065 | **0.027*** | 0 | **0.073** | **0.76** | 0.48 | ○●○●●○ |
| cas0051 | 11ba | -0.0036 | -0.03 | **0.091** | **0.056** | **0.61** | 0.47 | ○●●●○○ |
| cas0051 | 11bg | **0.1*** | -0.014 | **0.18** | **0.075** | **0.66** | 0.44 | ●○●●●○ |
| cas0051 | 1boz | -0.099 | **0.038*** | 0.033 | **0.062** | **0.7** | **0.61** | ○●○●●● |
| cas0051 | 1dlr | -0.056 | **0.018*** | **0.067** | **0.054** | **0.77** | **0.6** | ○●●●●● |
| cas0051 | 1dls | -0.079 | **0.034*** | 0 | **0.06** | **0.69** | **0.57** | ○●○●●● |
| cas0051 | 1drf | -0.1 | **0.037*** | 0.033 | **0.058** | **0.66** | **0.63** | ○●○●●● |
| cas0051 | 1hfp | -0.11 | **0.0082*** | 0.033 | 0.046 | **0.67** | **0.57** | ○●○○●● |
| cas0051 | 1r3m | **0.042*** | -0.034 | **0.18** | **0.064** | **0.65** | 0.48 | ●○●●●○ |
| cas0051 | 1r5c | **0.027*** | -0.041 | **0.18** | **0.068** | **0.56** | 0.37 | ●○●●●○ |
| cas0051 | 1tq9 | **0.034*** | -0.0038 | **0.18** | 0.047 | **0.65** | 0.46 | ●○●○●○ |
| cas0051 | 1xtu | **0.013*** | **0.025*** | 0.035 | 0.046 | 0.34 | 0.48 | ●●○○○○ |
| cas0051 | 3bcm | **0.026*** | -0.013 | **0.091** | **0.1** | **0.56** | 0.44 | ●○●●●○ |
| cas0051 | 3djo | **0.09*** | -0.021 | **0.091** | **0.058** | **0.73** | 0.47 | ●○●●●○ |
| cas0051 | 4r1r | -0.16 | -0.14 | 0 | 0.044 | 0.25 | 0.38 | ○○○○○○ |

| | | | | | | | | |
|---|---|---|---|---|---|---|---|---|
| cas0052 | 2z60 | -0.0083 | -0.034 | **0.079** | 0.042 | **0.68** | **0.54** | ○○●○●● |
| cas0054 | 1dhf | -0.15 | **0.0015*** | 0.033 | **0.066** | 0.45 | **0.52** | ○●○●○● |
| cas0054 | 2jfx | **0.056*** | -0.087 | **0.1** | 0.029 | **0.68** | **0.54** | ●○●○●● |
| cas0054 | 2jfy | **0.045*** | -0.092 | **0.1** | 0.022 | **0.67** | **0.54** | ●○●○●● |
| cas0054 | 3djv | **0.099*** | -0.0025 | **0.091** | **0.067** | **0.72** | 0.48 | ●○●●●○ |
| cas0056 | 4b1f | -0.13 | -0.16 | 0 | 0.016 | **0.57** | 0.48 | ○○○○●● |
| cas0060 | 1hfq | -0.077 | **0.034*** | 0.033 | **0.065** | **0.72** | **0.61** | ○●○●●● |
| cas0060 | 1hfr | -0.073 | **0.037*** | 0.033 | 0.049 | **0.71** | **0.62** | ○●○○●● |
| cas0060 | 1i4o | **0.2*** | **0.1*** | **0.15** | **0.12** | **0.89** | **0.77** | ●●●●●● |
| cas0060 | 1kms | -0.085 | **0.054*** | 0.033 | 0.038 | **0.74** | **0.63** | ○●○○●● |
| cas0060 | 1mvs | -0.11 | **0.026*** | 0.033 | **0.051** | **0.69** | **0.62** | ○●○●●● |
| cas0060 | 1mvt | -0.072 | -0.014 | 0 | 0.039 | **0.73** | **0.56** | ○○○○●● |
| cas0060 | 1ohj | -0.14 | **0.022*** | 0 | 0.033 | **0.61** | **0.61** | ○●○○●● |
| cas0060 | 1ohk | -0.093 | **0.023*** | **0.1** | 0.043 | **0.67** | **0.63** | ○●●○●● |
| cas0060 | 1pd8 | -0.1 | -0.0069 | **0.067** | **0.051** | **0.71** | **0.59** | ○○●●●● |
| cas0060 | 1pd9 | -0.12 | -0.016 | 0.033 | 0.034 | **0.69** | **0.58** | ○○○○●● |
| cas0060 | 1s3u | -0.11 | -0.047 | 0 | 0.039 | **0.7** | **0.54** | ○○○○●● |
| cas0060 | 1s3v | -0.12 | **0.029*** | 0 | 0.038 | **0.71** | **0.63** | ○●○○●● |
| cas0060 | 1s3w | -0.046 | **0.028*** | 0.033 | **0.05** | **0.77** | **0.62** | ○●○●●● |
| cas0060 | 1u71 | -0.088 | **0.036*** | 0.033 | **0.068** | **0.75** | **0.65** | ○●○●●● |
| cas0060 | 1u72 | -0.084 | -0.015 | 0.033 | **0.06** | **0.7** | **0.54** | ○○○●●● |
| cas0060 | 2c2s | **0.03*** | **0.064*** | 0.017 | **0.065** | 0.35 | 0.37 | ●●○●○○ |
| cas0060 | 2c2t | **0.045*** | **0.06*** | 0.033 | **0.062** | 0.36 | 0.37 | ●●○●○○ |
| cas0060 | 2dhf | -0.15 | **0.013*** | 0.033 | 0.042 | **0.5** | **0.54** | ○●○○●● |
| cas0060 | 2jfz | -0.14 | -0.16 | 0 | 0.013 | **0.57** | 0.48 | ○○○○●○ |
| cas0060 | 2psq | -0.011 | **0.054*** | 0 | **0.1** | **0.71** | **0.7** | ○●○●●● |

| | | | | | | | | |
|---|---|---|---|---|---|---|---|---|
| cas0060 | 2w3a | **0.029*** | **0.074*** | 0.033 | **0.059** | 0.33 | 0.35 | ●●○●○○ |
| cas0060 | 2w3m | **0.017*** | **0.089*** | 0.017 | **0.076** | 0.35 | 0.38 | ●●○●○○ |
| cas0060 | 3d6h | -0.0023 | -0.014 | **0.091** | **0.065** | **0.8** | **0.7** | ○○●●●● |
| cas0060 | 3djp | **0.059*** | -0.052 | **0.091** | **0.05** | **0.69** | 0.45 | ●○●●●○ |
| cas0060 | 3djq | **7.7e-05*** | -0.037 | **0.091** | 0.041 | **0.66** | 0.45 | ●○●○●○ |
| cas0060 | 3djx | **0.12*** | -0.0041 | **0.091** | **0.074** | **0.73** | 0.47 | ●○●●●○ |
| cas0060 | 3f8y | -0.12 | **0.029*** | 0 | 0.044 | **0.66** | **0.59** | ○●○○●● |
| cas0060 | 3f8z | -0.11 | **0.017*** | 0 | **0.053** | **0.69** | **0.6** | ○●○●●● |
| cas0060 | 3fs6 | -0.11 | **0.025*** | 0 | **0.052** | **0.7** | **0.58** | ○●○●●● |
| cas0060 | 3ghc | -0.074 | -0.021 | 0 | **0.051** | **0.74** | **0.54** | ○○○●●● |
| cas0060 | 3ghw | -0.081 | **0.013*** | 0 | 0.048 | **0.74** | **0.57** | ○●○○●● |
| cas0060 | 3gi2 | -0.099 | **0.022*** | 0.033 | 0.042 | **0.71** | **0.59** | ○●○○●● |
| cas0060 | 3gyf | -0.13 | **0.0025*** | 0 | **0.05** | **0.66** | **0.57** | ○●○●●● |
| cas0060 | 3hmi | -0.059 | **0.043*** | **0.12** | **0.058** | 0.46 | 0.43 | ○●●●○○ |
| cas0060 | 3l3r | -0.064 | **0.017*** | 0.033 | **0.053** | **0.75** | **0.58** | ○●○●●● |
| cas0060 | 3n0h | -0.16 | -0.022 | 0 | 0.039 | **0.56** | **0.52** | ○○○○●● |
| cas0060 | 3ntz | -0.099 | **0.0062*** | 0 | **0.058** | **0.71** | **0.57** | ○●○●●● |
| cas0060 | 3nu0 | -0.093 | **0.0038*** | 0 | **0.061** | **0.73** | **0.56** | ○●○●●● |
| cas0060 | 3nxr | -0.076 | **0.076*** | 0.033 | **0.061** | **0.74** | **0.65** | ○●○●●● |
| cas0060 | 3nxt | -0.15 | -0.013 | 0.033 | 0.042 | **0.61** | **0.55** | ○○○○●● |
| cas0060 | 3nxv | -0.048 | **0.019*** | **0.067** | **0.056** | **0.77** | **0.62** | ○●●●●● |
| cas0060 | 3nxx | -0.11 | **0.041*** | **0.067** | **0.063** | **0.71** | **0.62** | ○●●●●● |
| cas0060 | 3nxy | -0.047 | **0.011*** | **0.067** | **0.062** | **0.75** | **0.61** | ○●●●●● |
| cas0060 | 3nzd | -0.11 | **0.057*** | 0.033 | **0.061** | **0.72** | **0.63** | ○●○●●● |
| cas0060 | 3oaf | -0.057 | **0.049*** | 0.033 | **0.059** | **0.76** | **0.62** | ○●○●●● |
| cas0060 | 3s3v | -0.2 | -0.022 | 0 | 0.034 | **0.51** | **0.51** | ○○○○●● |

| | | | | | | | | |
|---|---|---|---|---|---|---|---|---|
| cas0060 | 3s7a | -0.025 | **0.082*** | **0.067** | **0.07** | **0.8** | **0.64** | ○●●●●● |
| cas0060 | 4ddr | -0.046 | **0.021*** | **0.067** | 0.049 | **0.77** | **0.58** | ○●●○●● |
| cas0060 | 4ej1 | **0.09*** | **0.095*** | **0.054** | **0.086** | **0.68** | **0.69** | ●●●●●○ |
| cas0060 | 4g95 | -0.13 | **0.021*** | 0 | 0.041 | **0.69** | **0.59** | ○●○○●● |
| cas0060 | 4i13 | -0.00094 | **0.11*** | **0.081** | **0.054** | **0.67** | **0.74** | ○●●●●● |
| cas0060 | 4i1n | -0.038 | **0.056*** | **0.081** | 0.034 | **0.64** | **0.71** | ○●●○●● |
| cas0060 | 4kd7 | -0.053 | **0.046*** | 0.017 | 0.049 | 0.49 | 0.48 | ○●○○○○ |
| cas0060 | 4keb | **0.048*** | **0.095*** | **0.067** | **0.087** | 0.48 | 0.46 | ●●●●○○ |
| cas0060 | 4m6k | -0.11 | **0.019*** | 0.033 | **0.087** | **0.67** | **0.58** | ○●○●●● |
| cas0060 | 4m6l | -0.1 | **0.042*** | 0.033 | **0.05** | **0.71** | **0.63** | ○●○●●● |
| cas0060 | 4n4c | -0.021 | **0.016*** | **0.091** | **0.057** | **0.5** | 0.45 | ○●●●●○ |
| cas0060 | 4qjc | -0.17 | -0.013 | 0.033 | 0.031 | **0.51** | **0.51** | ○○●●●● |
| cas0060 | 5hpb | -0.065 | **0.0032*** | 0 | 0.044 | **0.73** | **0.55** | ○●○○●● |
| cas0060 | 5hqy | -0.12 | -0.0025 | 0.033 | 0.04 | **0.71** | **0.53** | ○○○○●● |
| cas0060 | 5hqz | -0.11 | **0.0062*** | 0.033 | 0.048 | **0.72** | **0.6** | ○●○○●● |
| cas0060 | 5hsr | -0.032 | **0.016*** | 0.033 | **0.053** | **0.76** | **0.56** | ○●○●●● |
| cas0061 | 1dre | -0.08 | -0.041 | **0.081** | 0.029 | **0.65** | **0.57** | ○○●○●● |
| cas0061 | 1ibc | -0.13 | -0.065 | 0.045 | 0.042 | **0.67** | **0.64** | ○○○○●● |
| cas0061 | 1ibv | **0.46*** | **0.022*** | **0.67** | **0.15** | **0.85** | **0.65** | ●●●●●● |
| cas0061 | 1ibw | **0.32*** | **0.086*** | **0.67** | **0.25** | **0.76** | **0.67** | ●●●●●● |
| cas0061 | 1ice | -0.12 | -0.12 | **0.091** | 0.04 | **0.7** | **0.59** | ○○●○●● |
| cas0061 | 1pya | **0.25*** | **0.032*** | 0 | **0.12** | **0.66** | **0.6** | ●●○●●● |
| cas0061 | 1rc4 | -0.049 | -0.037 | **0.081** | 0.045 | **0.69** | **0.6** | ○○●○●● |
| cas0061 | 1rwm | -0.0098 | -0.017 | 0.045 | **0.088** | **0.77** | **0.66** | ○○○●●● |
| cas0061 | 1rwn | -0.081 | -0.069 | 0.045 | 0.042 | **0.74** | **0.62** | ○○○○●● |
| cas0061 | 1rwo | -0.073 | -0.087 | **0.091** | 0.042 | **0.75** | **0.6** | ○○●○●● |

| | | | | | | | | |
|---|---|---|---|---|---|---|---|---|
| cas0061 | 1rwv | -0.099 | -0.055 | **0.091** | **0.059** | **0.73** | **0.65** | ○○●●●● |
| cas0061 | 1rx4 | -0.062 | -0.027 | **0.054** | 0.02 | **0.73** | **0.59** | ○○●○●● |
| cas0061 | 1rx6 | -0.081 | -0.014 | **0.054** | 0.03 | **0.73** | **0.6** | ○○●○●● |
| cas0061 | 2h4w | -0.067 | -0.031 | 0.045 | **0.052** | **0.74** | **0.66** | ○○○●●● |
| cas0061 | 2h4y | -0.024 | -0.067 | **0.091** | 0.036 | **0.79** | **0.66** | ○○●○●● |
| cas0061 | 2h51 | -0.11 | -0.08 | 0 | 0.019 | **0.72** | **0.64** | ○○○○●● |
| cas0061 | 2h54 | -0.021 | -0.049 | **0.091** | 0.047 | **0.79** | **0.65** | ○○●○●● |
| cas0061 | 2hbq | -0.039 | -0.035 | **0.091** | **0.054** | **0.78** | **0.66** | ○○●●●● |
| cas0061 | 2hbr | -0.11 | -0.049 | 0 | 0.049 | **0.71** | **0.66** | ○○○○●● |
| cas0061 | 2hby | -0.043 | **0.0056*** | 0.045 | **0.059** | **0.76** | **0.67** | ○●○●●● |
| cas0061 | 2hbz | -0.085 | -0.072 | 0 | **0.052** | **0.73** | **0.65** | ○○○●●● |
| cas0061 | 3d6f | **0.014*** | -0.016 | 0.045 | **0.079** | **0.81** | **0.67** | ●○○●●● |
| cas0061 | 3d6m | -0.033 | -0.05 | **0.14** | **0.053** | **0.8** | **0.65** | ○○●●●● |
| cas0061 | 3ri1 | -0.0066 | **0.0086*** | 0 | **0.081** | **0.71** | **0.67** | ○●○●●● |
| cas0061 | 4kjl | -0.069 | **0.018*** | **0.081** | 0.044 | **0.72** | **0.56** | ○●●○●● |
| cas0061 | 4p3r | -0.061 | -0.015 | **0.11** | 0.044 | **0.72** | **0.59** | ○○●○●● |
| cas0061 | 5ccc | -0.08 | **0.02*** | **0.054** | 0.022 | **0.68** | **0.64** | ○●●○●● |
| cas0061 | 5hsu | -0.13 | -0.0024 | 0 | 0.035 | **0.72** | **0.58** | ○○○○●● |
| cas0061 | 5ht4 | -0.06 | **0.017*** | 0.033 | **0.062** | **0.76** | **0.61** | ○●○●●● |
| cas0061 | 5ht5 | -0.1 | -0.0063 | 0.033 | 0.038 | **0.7** | **0.57** | ○○○○●● |
| cas0061 | 5hui | -0.11 | **0.024*** | 0.033 | **0.054** | **0.74** | **0.59** | ○●○●●● |
| cas0061 | 5hvb | -0.083 | **0.014*** | 0 | 0.039 | **0.75** | **0.59** | ○●○○●● |
| cas0061 | 5hve | -0.14 | **0.027*** | 0.033 | 0.046 | **0.69** | **0.61** | ○●○○●● |
| cas0067 | 1rx7 | -0.041 | -0.085 | **0.11** | 0.044 | **0.7** | **0.56** | ○○●○●● |
| cas0067 | 4qle | -0.033 | **0.0056*** | **0.081** | 0.024 | **0.57** | **0.56** | ○●●○●● |
| cas0070 | 1ra2 | -0.11 | -0.1 | **0.054** | 0.012 | **0.6** | 0.48 | ○○●○●○ |

| | | | | | | | | |
|---|---|---|---|---|---|---|---|---|
| cas0070 | 1ra3 | -0.12 | -0.052 | **0.054** | 0 | **0.61** | **0.51** | ○○●○●● |
| cas0070 | 1ra8 | -0.093 | -0.1 | **0.054** | 0.012 | **0.62** | 0.48 | ○○●○●○ |
| cas0070 | 1rb3 | -0.062 | -0.03 | **0.068** | 0.011 | 0.48 | 0.47 | ○○●○○○ |
| cas0070 | 1rg7 | -0.095 | -0.062 | 0.027 | 0.016 | **0.61** | 0.46 | ○○○○●○ |
| cas0070 | 1rh3 | -0.085 | -0.061 | **0.081** | 0.013 | **0.65** | **0.54** | ○○●○●● |
| cas0070 | 1rx5 | -0.077 | -0.11 | **0.11** | 0.01 | **0.7** | **0.55** | ○○●○●● |
| cas0070 | 3dau | -0.083 | -0.0078 | 0.027 | 0.023 | **0.65** | **0.53** | ○○○○●● |
| cas0070 | 4fhb | -0.011 | -0.11 | **0.11** | 0.022 | **0.58** | **0.51** | ○○●○●● |
| cas0070 | 4kjj | -0.059 | -0.029 | **0.054** | 0.033 | **0.7** | **0.53** | ○○●○●● |
| cas0070 | 4qlg | -0.02 | -0.037 | 0.041 | 0.031 | **0.51** | 0.48 | ○○○○●○ |
| cas0070 | 4x5g | -0.14 | -0.14 | **0.054** | 0.011 | **0.6** | **0.55** | ○○●○●● |
| cas0070 | 4x5h | -0.078 | -0.094 | **0.054** | 0.022 | **0.62** | **0.5** | ○○●○●● |
| cas0070 | 4x5i | -0.063 | -0.13 | **0.054** | 0.022 | **0.62** | 0.46 | ○○●○●○ |
| cas0070 | 4x5j | -0.064 | -0.1 | **0.081** | 0.02 | **0.63** | 0.48 | ○○●○●○ |
| cas0070 | 5cc9 | -0.12 | -0.074 | 0.027 | 0.024 | **0.63** | **0.55** | ○○○○●● |
| cas0070 | 7dfr | -0.073 | -0.075 | **0.081** | 0.036 | **0.7** | **0.5** | ○○●○●● |
| cas0071 | 1rd7 | -0.039 | -0.065 | **0.081** | 0.022 | 0.47 | 0.44 | ○○●○○○ |
| cas0074 | 1rb2 | -0.0079 | -0.1 | **0.11** | 0.02 | **0.5** | 0.42 | ○○●○●○ |
| cas0074 | 1re7 | -0.045 | -0.1 | **0.11** | 0 | 0.46 | 0.41 | ○○●○○○ |
| cas0074 | 4x5f | -0.13 | -0.14 | **0.068** | 0.029 | **0.52** | 0.42 | ○○●○●○ |
| cas0079 | 1bzc | **0.16*** | **0.05*** | **0.11** | **0.084** | **0.82** | **0.59** | ●●●●●● |
| cas0079 | 1bzj | **0.23*** | **0.12*** | **0.22** | **0.11** | **0.87** | **0.66** | ●●●●●● |
| cas0079 | 1c83 | **0.19*** | **0.099*** | **0.17** | **0.11** | **0.87** | **0.66** | ●●●●●● |
| cas0079 | 1c84 | **0.19*** | **0.068*** | **0.22** | **0.072** | **0.88** | **0.63** | ●●●●●● |
| cas0079 | 1c85 | **0.22*** | **0.067*** | **0.17** | **0.11** | **0.88** | **0.63** | ●●●●●● |
| cas0079 | 1c86 | **0.28*** | **0.14*** | **0.33** | **0.13** | **0.91** | **0.7** | ●●●●●● |

| | | | | | | | | |
|---|---|---|---|---|---|---|---|---|
| cas0079 | 1c87 | **0.25*** | **0.1*** | **0.22** | **0.13** | **0.9** | **0.66** | ●●●●●● |
| cas0079 | 1c88 | **0.27*** | **0.1*** | **0.33** | **0.12** | **0.91** | **0.67** | ●●●●●● |
| cas0079 | 1ecv | **0.21*** | **0.12*** | **0.17** | **0.13** | **0.88** | **0.66** | ●●●●●● |
| cas0079 | 1g1g | **0.21*** | **0.087*** | **0.11** | **0.1** | **0.81** | **0.57** | ●●●●●● |
| cas0079 | 1g7g | **0.17*** | **0.07*** | **0.11** | **0.12** | **0.87** | **0.63** | ●●●●●● |
| cas0079 | 1gfy | **0.23*** | **0.089*** | **0.22** | **0.11** | **0.89** | **0.66** | ●●●●●● |
| cas0079 | 1kak | **0.14*** | **0.074*** | **0.056** | **0.1** | **0.84** | **0.62** | ●●●●●● |
| cas0079 | 1kav | **0.16*** | **0.1*** | **0.056** | **0.083** | **0.84** | **0.62** | ●●●●●● |
| cas0079 | 1l8g | **0.23*** | **0.11*** | **0.17** | **0.13** | **0.9** | **0.68** | ●●●●●● |
| cas0079 | 1nwe | **0.12*** | **0.11*** | **0.11** | **0.076** | **0.82** | **0.66** | ●●●●●● |
| cas0079 | 1ptt | **0.18*** | **0.071*** | **0.056** | **0.059** | **0.78** | **0.57** | ●●●●●● |
| cas0079 | 1ptu | **0.25*** | **0.11*** | **0.11** | **0.099** | **0.74** | **0.62** | ●●●●●● |
| cas0079 | 1ptv | **0.25*** | **0.13*** | **0.22** | **0.12** | **0.89** | **0.67** | ●●●●●● |
| cas0079 | 1q1m | **0.19*** | **0.1*** | **0.28** | **0.13** | **0.87** | **0.67** | ●●●●●● |
| cas0079 | 1xbo | **0.2*** | **0.095*** | **0.33** | **0.11** | **0.87** | **0.66** | ●●●●●● |
| cas0079 | 2azr | **0.24*** | **0.1*** | **0.33** | **0.14** | **0.9** | **0.67** | ●●●●●● |
| cas0079 | 2b07 | **0.18*** | **0.089*** | **0.22** | **0.12** | **0.88** | **0.67** | ●●●●●● |
| cas0079 | 2bge | **0.23*** | **0.11*** | **0.39** | **0.11** | **0.88** | **0.65** | ●●●●●● |
| cas0079 | 2cm7 | **0.22*** | **0.1*** | **0.22** | **0.11** | **0.88** | **0.64** | ●●●●●● |
| cas0079 | 2cma | **0.16*** | **0.092*** | **0.17** | **0.09** | **0.85** | **0.63** | ●●●●●● |
| cas0079 | 2h4g | **0.17*** | **0.082*** | **0.17** | **0.11** | **0.87** | **0.67** | ●●●●●● |
| cas0079 | 2h4k | **0.18*** | **0.073*** | **0.22** | **0.13** | **0.87** | **0.65** | ●●●●●● |
| cas0079 | 2hb1 | **0.2*** | **0.094*** | **0.22** | **0.11** | **0.89** | **0.66** | ●●●●●● |
| cas0079 | 2nt7 | **0.19*** | **0.12*** | **0.28** | **0.12** | **0.88** | **0.68** | ●●●●●● |
| cas0079 | 2nta | **0.2*** | **0.087*** | **0.11** | **0.12** | **0.86** | **0.64** | ●●●●●● |
| cas0079 | 2qbp | **0.16*** | **0.12*** | **0.22** | **0.12** | **0.87** | **0.69** | ●●●●●● |

| | | | | | | | | |
|---|---|---|---|---|---|---|---|---|
| cas0079 | 2qbq | **0.17*** | **0.11*** | **0.22** | **0.097** | **0.88** | **0.68** | ●●●●●● |
| cas0079 | 2qbr | **0.19*** | **0.081*** | **0.22** | **0.11** | **0.88** | **0.66** | ●●●●●● |
| cas0079 | 2qbs | **0.15*** | **0.11*** | **0.17** | **0.1** | **0.87** | **0.69** | ●●●●●● |
| cas0079 | 2veu | **0.19*** | **0.087*** | **0.11** | **0.091** | **0.87** | **0.62** | ●●●●●● |
| cas0079 | 2vey | **0.16*** | **0.1*** | **0.22** | **0.11** | **0.85** | **0.63** | ●●●●●● |
| cas0079 | 2zn7 | **0.21*** | **0.075*** | **0.22** | **0.13** | **0.89** | **0.65** | ●●●●●● |
| cas0079 | 4i8n | **0.26*** | **0.13*** | **0.33** | **0.16** | **0.91** | **0.67** | ●●●●●● |
| cas0080 | 1i00 | -0.021 | **0.034*** | **0.19** | **0.095** | **0.68** | **0.64** | ●○●●●● |
| cas0080 | 1s2q | **0.25*** | **0.069*** | **0.18** | 0.025 | **0.59** | **0.63** | ●●●○●● |
| cas0080 | 1s2y | **0.2*** | **0.045*** | **0.21** | 0.022 | **0.56** | **0.61** | ●●●○●● |
| cas0080 | 1s3b | **0.24*** | **0.058*** | **0.21** | 0.025 | **0.58** | **0.62** | ●●●○●● |
| cas0080 | 1s3e | **0.25*** | **0.058*** | **0.24** | 0.03 | **0.6** | **0.62** | ●●●○●● |
| cas0080 | 5x67 | **0.066*** | **0.051*** | **0.19** | **0.13** | **0.76** | **0.66** | ●●●●●● |
| cas0085 | 1gos | **0.18*** | **0.017*** | **0.29** | **0.086** | **0.56** | **0.58** | ●●●●●● |
| cas0085 | 1oj9 | **0.18*** | **0.02*** | **0.15** | 0.011 | **0.55** | **0.6** | ●●●○●● |
| cas0085 | 1oja | **0.14*** | -0.018 | **0.12** | 0.0077 | **0.51** | **0.56** | ●○●○●● |
| cas0085 | 5x5q | -0.065 | **0.043*** | **0.19** | **0.1** | 0.38 | **0.58** | ○●●●○● |
| cas0086 | 1fin | **0.12*** | **0.11*** | **0.12** | **0.064** | **0.81** | **0.78** | ●●●●●● |
| cas0086 | 1fuo | -0.24 | -0.15 | 0 | 0.014 | 0.32 | **0.53** | ○○○○○● |
| cas0086 | 1fup | -0.26 | -0.18 | 0 | 0.014 | 0.3 | **0.5** | ○○○○○● |
| cas0086 | 1fuq | -0.23 | -0.19 | 0 | 0.0066 | 0.34 | 0.48 | ○○○○○○ |
| cas0086 | 1juj | -0.16 | -0.063 | 0 | 0.028 | 0.32 | **0.55** | ○○○○○● |
| cas0086 | 1kq7 | -0.2 | -0.2 | 0 | 0.014 | 0.36 | 0.48 | ○○○○○○ |
| cas0086 | 1ojc | **0.054*** | -0.086 | **0.059** | 0.016 | 0.48 | **0.53** | ●○●○○● |
| cas0086 | 2b54 | **0.025*** | -0.028 | **0.12** | 0.039 | **0.77** | **0.64** | ●○●○●● |
| cas0086 | 2bk3 | **0.17*** | **0.012*** | **0.12** | 0.0054 | **0.54** | **0.58** | ●●●○●● |

| | | | | | | | | |
|---|---|---|---|---|---|---|---|---|
| cas0086 | 2bk4 | **0.28*** | **0.075*** | **0.18** | 0.029 | **0.62** | **0.63** | ●●●○●● |
| cas0086 | 2bk5 | **0.18*** | **0.0045*** | **0.12** | 0.0049 | **0.54** | **0.58** | ●●●○●● |
| cas0086 | 2byb | **0.23*** | **0.064*** | **0.21** | **0.053** | **0.58** | **0.62** | ●●●●●● |
| cas0086 | 2c64 | **0.19*** | **0.045*** | **0.21** | 0.029 | **0.56** | **0.61** | ●●●○●● |
| cas0086 | 2c65 | **0.17*** | **0.0097*** | **0.059** | 0.011 | **0.54** | **0.59** | ●●●○●● |
| cas0086 | 2c66 | **0.21*** | **0.07*** | **0.29** | **0.068** | **0.58** | **0.63** | ●●●●●● |
| cas0086 | 2c67 | **0.14*** | -0.021 | **0.12** | 0.005 | **0.51** | **0.56** | ●○●○●● |
| cas0086 | 2c70 | **0.2*** | **0.03*** | **0.12** | 0.01 | **0.57** | **0.59** | ●●●○●● |
| cas0086 | 2c72 | **0.22*** | **0.057*** | **0.29** | 0.039 | **0.58** | **0.62** | ●●●○●● |
| cas0086 | 2c73 | **0.23*** | **0.073*** | **0.21** | 0.047 | **0.59** | **0.63** | ●●●○●● |
| cas0086 | 2c75 | **0.26*** | **0.075*** | **0.18** | 0.027 | **0.6** | **0.63** | ●●●○●● |
| cas0086 | 2c76 | **0.27*** | **0.079*** | **0.18** | 0.027 | **0.61** | **0.63** | ●●●○●● |
| cas0086 | 2cch | **0.12*** | **0.066*** | **0.058** | **0.084** | **0.68** | **0.69** | ●●●●●● |
| cas0086 | 2v5z | **0.14*** | **0.012*** | **0.12** | 0.0055 | **0.53** | **0.58** | ●●●○●● |
| cas0086 | 2v60 | **0.19*** | **0.03*** | **0.18** | 0.011 | **0.56** | **0.6** | ●●●○●● |
| cas0086 | 2v61 | **0.15*** | **0.015*** | **0.15** | 0.0053 | **0.53** | **0.58** | ●●●○●● |
| cas0086 | 2vrl | **0.13*** | **0.0056*** | **0.12** | 0.0057 | **0.51** | **0.58** | ●●●○●● |
| cas0086 | 2vrm | **0.17*** | **0.034*** | **0.15** | 0.011 | **0.54** | **0.6** | ●●●○●● |
| cas0086 | 2vz2 | **0.13*** | **0.0068*** | **0.059** | 0.01 | **0.51** | **0.58** | ●●●○●● |
| cas0086 | 2xfn | **0.14*** | **0.016*** | **0.15** | 0.0052 | **0.53** | **0.58** | ●●●○●● |
| cas0086 | 2xfp | **0.21*** | **0.024*** | **0.18** | 0.0048 | **0.56** | **0.59** | ●●●○●● |
| cas0086 | 2xfq | **0.2*** | **0.018*** | **0.12** | 0.0092 | **0.56** | **0.59** | ●●●○●● |
| cas0086 | 3ob7 | -0.014 | -0.014 | **0.05** | **0.079** | **0.54** | **0.51** | ○○●●●● |
| cas0086 | 3po7 | **0.22*** | **0.024*** | **0.15** | 0.0074 | **0.57** | **0.6** | ●●●○●● |
| cas0086 | 3zyx | **0.075*** | -0.0062 | **0.15** | 0.011 | 0.48 | **0.56** | ●○●○○● |
| cas0086 | 4a79 | **0.085*** | -0.017 | **0.088** | 0.0087 | 0.49 | **0.56** | ●○●○○● |

| | | | | | | | | |
|---|---|---|---|---|---|---|---|---|
| cas0086 | 4a7a | **0.15*** | **0.021*** | **0.088** | 0.0055 | **0.53** | **0.59** | ●●●○●● |
| cas0086 | 4crt | **0.17*** | **0.011*** | **0.15** | 0.0081 | **0.54** | **0.58** | ●●●○●● |
| cas0086 | 5hs3 | -0.12 | -0.056 | 0.042 | **0.076** | 0.25 | 0.47 | ○○○●○○ |
| cas0086 | 5mrl | **0.18*** | **0.038*** | **0.18** | **0.059** | **0.55** | **0.6** | ●●●●●● |
| cas0091 | 2xcg | **0.082*** | -0.0064 | **0.059** | 0.0052 | 0.48 | **0.56** | ●○●○○● |
| cas0091 | 2xfo | **0.064*** | -0.012 | **0.088** | 0.0083 | 0.46 | **0.55** | ●○●○○● |
| cas0091 | 2xfu | **0.029*** | -0.056 | **0.059** | 0.011 | 0.44 | **0.51** | ●○●○○● |
| cas0091 | 4ez7 | -0.096 | -0.13 | 0 | 0.0043 | **0.55** | 0.42 | ○○○○●○ |

**Table S7: Allosteric site quantile scores of proteins in Table S5 (orthosteric site residues as the source with orthosteric ligands removed)** The results from six statistical scores described in Methods. Average site residue and bond quantile scores are compared with those of 1000 surrogate sites of the same size. The difference is shown in bold if it is above 0 and starred if it is above the 95% confidence interval. The proportion of residues/bonds with $p_{R/b,\text{allo}} > 0.95$ and the average reference quantile score $\overline{p^{\text{ref}}_{R/b,\text{ allo}}}$ are shown in bold if they are above the expected values of 0.05 and 0.5 respectively.

| cas Number | PDB | $\overline{p_{R,\text{allo}}}$ - $\langle\overline{p_{R,\text{site}}}\rangle_{\text{surr}}$ | $\overline{p_{b,\text{allo}}}$ - $\langle\overline{p_{b,\text{site}}}\rangle_{\text{surr}}$ | P($p_{R,\text{allo}}$ > 0.95) | P($p_{b,\text{allo}}$ > 0.95) | $\overline{p^{\text{ref}}_{R,\text{ allo}}}$ | $\overline{p^{\text{ref}}_{b,\text{ allo}}}$ | Summary |
|---|---|---|---|---|---|---|---|---|
| cas0001 | 1nxe | -0.016 | -0.053 | 0 | 0.048 | 0.44 | **0.6** | ○○○○○● |
| cas0001 | 1nxg | **0.012*** | -0.034 | 0 | 0.049 | 0.46 | **0.62** | ●○○○○● |
| cas0001 | 1owc | **0.013*** | -0.019 | 0 | **0.053** | 0.46 | **0.63** | ●○○●○● |
| cas0001 | 4g6b | **0.014*** | -0.036 | 0 | **0.062** | 0.46 | **0.61** | ●○○●○● |
| cas0001 | 4jad | **0.02*** | -0.016 | 0 | **0.05** | 0.47 | **0.62** | ●○○●○● |
| cas0001 | 4jae | **0.0033*** | -0.075 | 0 | 0.047 | 0.45 | **0.59** | ●○○○○● |
| cas0001 | 4jaf | **0.038*** | -0.035 | 0 | **0.055** | 0.48 | **0.61** | ●○○●○● |
| cas0001 | 4jag | **0.056*** | -0.0006 | 0 | **0.053** | **0.51** | **0.64** | ●○○●●● |
| cas0002 | 3i1y | **0.012*** | **0.1*** | 0 | 0.043 | 0.095 | 0.11 | ●●○○○○ |
| cas0002 | 3i28 | **0.058*** | **0.083*** | 0.033 | 0.047 | 0.12 | 0.12 | ●●○○○○ |
| cas0002 | 3koo | **0.023*** | **0.061*** | 0 | 0.032 | 0.13 | 0.13 | ●●○○○○ |
| cas0002 | 3otq | **0.051*** | **0.11*** | **0.067** | **0.064** | 0.12 | 0.14 | ●●●●○○ |
| cas0002 | 5ahx | **0.013*** | **0.062*** | 0 | **0.053** | **0.5** | 0.49 | ●●○●●○ |
| cas0002 | 5ai4 | **0.064*** | **0.11*** | 0 | **0.057** | 0.12 | 0.13 | ●●○●○○ |
| cas0002 | 5ai5 | -0.031 | **0.049*** | 0.033 | 0.038 | 0.47 | 0.49 | ○●○○○○ |
| cas0002 | 5aia | -0.22 | -0.14 | 0 | 0 | 0.14 | 0.19 | ○○○○○○ |
| cas0002 | 5ak4 | **0.054*** | **0.1*** | 0 | 0.043 | 0.12 | 0.12 | ●●○○○○ |
| cas0002 | 5ak5 | -0.009 | -0.012 | 0 | 0.02 | 0.48 | 0.44 | ○○○○○○ |
| cas0002 | 5ake | -0.025 | **0.024*** | 0.033 | 0.033 | 0.47 | 0.48 | ○●○○○○ |
| cas0002 | 5akh | -0.0025 | **0.055*** | 0.033 | 0.047 | 0.48 | 0.49 | ○●○○○○ |

| | | | | | | | | |
|---|---|---|---|---|---|---|---|---|
| cas0002 | 5akx | **0.0052*** | **0.042*** | 0 | 0.029 | **0.5** | 0.49 | ●●○○●○ |
| cas0002 | 5aky | -0.015 | **0.018*** | 0 | **0.056** | 0.47 | 0.47 | ○●○●○○ |
| cas0002 | 5ald | **0.0041*** | **0.043*** | 0.033 | 0.047 | **0.5** | 0.49 | ●●○○●○ |
| cas0002 | 5alf | -0.0075 | **0.043*** | 0 | 0.043 | **0.5** | 0.49 | ○●○○●○ |
| cas0002 | 5alh | -0.0048 | **0.05*** | 0 | 0.039 | 0.49 | 0.49 | ○●○○○○ |
| cas0002 | 5alm | -0.031 | **0.046*** | 0.033 | 0.034 | 0.47 | 0.49 | ○●○○○○ |
| cas0002 | 5aln | **0.062*** | **0.089*** | 0 | **0.062** | 0.13 | 0.14 | ●●○●○○ |
| cas0002 | 5alo | **0.065*** | **0.093*** | 0 | 0.044 | 0.12 | 0.13 | ●●○○○○ |
| cas0002 | 5alt | **0.055*** | **0.13*** | 0 | **0.053** | 0.12 | 0.13 | ●●○●○○ |
| cas0002 | 5alu | **0.051*** | **0.11*** | 0 | **0.068** | 0.13 | 0.14 | ●●○●○○ |
| cas0002 | 5alv | **0.086*** | **0.12*** | 0.033 | **0.058** | 0.094 | 0.095 | ●●○●○○ |
| cas0002 | 5alw | **0.046*** | **0.11*** | 0 | **0.068** | 0.15 | 0.16 | ●●○●○○ |
| cas0002 | 5aly | **0.075*** | **0.11*** | 0 | 0.047 | 0.1 | 0.11 | ●●○○○○ |
| cas0002 | 5am0 | **0.065*** | **0.12*** | 0 | **0.068** | 0.12 | 0.13 | ●●○●○○ |
| cas0002 | 5am4 | **0.067*** | **0.097*** | 0 | **0.054** | 0.13 | 0.14 | ●●○●○○ |
| cas0002 | 5am5 | **0.081*** | **0.13*** | 0 | **0.064** | 0.13 | 0.14 | ●●○●○○ |
| cas0003 | 3ion | **0.24*** | -0.012 | **0.17** | **0.073** | **0.82** | **0.55** | ●○●●●● |
| cas0003 | 3iop | **0.26*** | -0.0097 | **0.17** | **0.078** | **0.82** | **0.54** | ●○●●●● |
| cas0003 | 3rwp | **0.24*** | -0.0033 | **0.22** | **0.073** | **0.82** | **0.54** | ●○●●●● |
| cas0003 | 4rqk | **0.24*** | -0.044 | **0.11** | **0.062** | **0.83** | **0.51** | ●○●●●● |
| cas0003 | 4rqv | **0.24*** | -0.038 | **0.22** | **0.065** | **0.81** | **0.51** | ●○●●●● |
| cas0003 | 4rrv | **0.2*** | **0.049*** | **0.17** | **0.082** | **0.79** | **0.62** | ●●●●●● |
| cas0003 | 4xx9 | **0.25*** | -0.039 | **0.17** | **0.082** | **0.83** | **0.53** | ●○●●●● |
| cas0004 | 4ey5 | **0.24*** | **0.098*** | **0.42** | 0.034 | **0.72** | **0.69** | ●●●○●● |
| cas0004 | 5hf6 | **0.2*** | -0.029 | **0.17** | 0.036 | **0.68** | **0.6** | ●○●○●● |
| cas0004 | 5hf8 | **0.22*** | -0.052 | **0.17** | **0.063** | **0.71** | **0.55** | ●○●●●● |

| | | | | | | | | |
|---|---|---|---|---|---|---|---|---|
| cas0004 | 5hf9 | **0.24*** | **0.0047*** | **0.25** | 0.039 | **0.71** | **0.62** | ●●●○●● |
| cas0004 | 5hfa | **0.11*** | -0.048 | **0.17** | 0.025 | **0.63** | **0.57** | ●○●○●● |
| cas0010 | 1i2d | -0.076 | -0.13 | 0 | 0.023 | **0.57** | **0.61** | ○○○○●● |
| cas0011 | 2ewn | **0.46*** | -0.097 | **0.2** | **0.055** | **0.81** | 0.4 | ●○●●●○ |
| cas0015 | 2ym4 | **0.00082*** | -0.16 | **0.095** | 0.041 | **0.74** | 0.49 | ●○●○●○ |
| cas0015 | 2ym8 | **0.043*** | -0.14 | **0.19** | **0.053** | **0.76** | **0.51** | ●○●●●● |
| cas0015 | 4fst | -0.022 | -0.1 | **0.14** | **0.052** | **0.72** | **0.53** | ●○●●●● |
| cas0015 | 4fsy | -0.043 | -0.12 | 0.048 | 0.047 | **0.7** | **0.53** | ○○○○●● |
| cas0015 | 4ft3 | **0.0029*** | -0.12 | **0.095** | 0.042 | **0.74** | **0.52** | ●○●○●● |
| cas0015 | 4ft7 | -0.045 | -0.12 | 0.048 | 0.032 | **0.72** | **0.51** | ○○○○●● |
| cas0015 | 4fta | -0.0054 | -0.14 | **0.19** | **0.055** | **0.76** | **0.51** | ○○●●●● |
| cas0015 | 4ftn | **0.013*** | -0.11 | **0.14** | **0.068** | **0.76** | **0.51** | ●○●●●● |
| cas0015 | 4fto | -0.0047 | -0.12 | **0.095** | 0.047 | **0.73** | **0.52** | ○○●○●● |
| cas0015 | 4ftr | **0.035*** | -0.12 | **0.19** | **0.06** | **0.76** | **0.51** | ●○●●●● |
| cas0015 | 4ftu | **0.024*** | -0.16 | **0.19** | 0.046 | **0.77** | **0.51** | ●○●●●● |
| cas0015 | 4gh2 | -0.028 | -0.12 | **0.14** | **0.06** | **0.72** | **0.54** | ○○●●●● |
| cas0016 | 3csm | **0.044*** | **0.076*** | **0.056** | 0.039 | 0.41 | 0.43 | ●●●○○○ |
| cas0016 | 4csm | **0.046*** | **0.091*** | **0.056** | **0.069** | 0.41 | 0.42 | ●●●●○○ |
| cas0021 | 3l9h | **0.14*** | **0.06*** | **0.097** | 0.047 | **0.56** | **0.56** | ●●●○●● |
| cas0021 | 4zhi | **0.055*** | **0.04*** | **0.065** | 0.035 | **0.52** | **0.54** | ●●●○●● |
| cas0024 | 4ald | **0.26*** | **0.011*** | 0 | 0 | **0.66** | **0.63** | ●●○○●● |
| cas0027 | 1cza | **0.33*** | **0.1*** | **0.13** | **0.12** | 0.42 | 0.29 | ●●●●○○ |
| cas0027 | 1dgk | **0.31*** | **0.1*** | **0.067** | **0.11** | 0.35 | 0.26 | ●●●●○○ |
| cas0028 | 1lld | -0.21 | -0.16 | 0 | 0.0075 | 0.37 | **0.5** | ○○○○○● |
| cas0028 | 1lth | -0.09 | -0.17 | 0 | 0.011 | 0.45 | 0.48 | ○○○○○○ |
| cas0029 | 1ldn | -0.017 | **0.076*** | 0 | 0.016 | 0.079 | 0.46 | ○●○○○○ |

| | | | | | | | | |
|---|---|---|---|---|---|---|---|---|
| cas0030 | 2him | -0.00065 | **0.042*** | 0 | 0.025 | 0.28 | **0.58** | ○●○○○● |
| cas0030 | 2p2d | **0.01*** | **0.094*** | 0.021 | 0.025 | 0.29 | **0.6** | ●●○○○● |
| cas0039 | 3ddn | -0.25 | -0.14 | 0 | 0.0047 | **0.51** | **0.58** | ○○○○●● |
| cas0040 | 1psd | -0.034 | **0.1*** | 0 | **0.051** | 0.43 | **0.64** | ○●○●○● |
| cas0040 | 1yba | **0.11*** | **0.19*** | **0.067** | **0.1** | 0.32 | **0.52** | ●●●●○● |
| cas0040 | 2p9c | **0.04*** | **0.14*** | 0.033 | **0.055** | 0.45 | **0.64** | ●●○●○● |
| cas0040 | 2p9e | **0.069*** | **0.19*** | **0.1** | **0.11** | 0.32 | **0.53** | ●●●●○● |
| cas0040 | 2p9g | **0.16*** | **0.25*** | **0.17** | **0.14** | **0.56** | **0.75** | ●●●●●● |
| cas0040 | 2pa3 | **0.043*** | **0.21*** | **0.067** | **0.09** | **0.51** | **0.75** | ●●●●●● |
| cas0047 | 4r1r | -0.11 | -0.11 | **0.15** | 0.04 | 0.23 | 0.4 | ○○●○○○ |
| cas0050 | 1xtu | **0.044*** | **0.0071*** | **0.11** | **0.059** | 0.19 | 0.41 | ●●●●○○ |
| cas0050 | 1xtv | **0.018*** | -0.012 | **0.12** | **0.052** | 0.18 | 0.4 | ●○●●○○ |
| cas0051 | 11ba | **0.24*** | **0.14*** | **0.27** | **0.078** | **0.63** | 0.49 | ●●●●●○ |
| cas0051 | 11bg | **0.29*** | **0.15*** | **0.18** | **0.066** | **0.67** | 0.47 | ●●●●●○ |
| cas0051 | 1bsr | **0.27*** | **0.15*** | **0.18** | **0.06** | **0.69** | 0.49 | ●●●●●○ |
| cas0051 | 1n3z | **0.2*** | **0.21*** | **0.12** | 0.049 | **0.74** | 0.49 | ●●●○●○ |
| cas0051 | 1r3m | **0.18*** | **0.14*** | **0.27** | **0.064** | **0.57** | 0.47 | ●●●●●○ |
| cas0051 | 1r5c | **0.27*** | **0.13*** | **0.27** | **0.087** | **0.66** | 0.46 | ●●●●●○ |
| cas0051 | 1tq9 | **0.26*** | **0.16*** | **0.18** | **0.07** | **0.66** | 0.49 | ●●●●●○ |
| cas0051 | 3bcm | **0.21*** | **0.19*** | **0.18** | **0.1** | **0.58** | 0.48 | ●●●●●○ |
| cas0051 | 3djo | **0.25*** | **0.15*** | **0.27** | **0.058** | **0.67** | 0.48 | ●●●●●○ |
| cas0051 | 3djp | **0.24*** | **0.12*** | **0.18** | **0.059** | **0.67** | 0.46 | ●●●●●○ |
| cas0051 | 3djq | **0.23*** | **0.13*** | **0.27** | **0.071** | **0.66** | 0.47 | ●●●●●○ |
| cas0051 | 3djv | **0.25*** | **0.13*** | **0.27** | **0.077** | **0.66** | 0.46 | ●●●●●○ |
| cas0051 | 3djx | **0.27*** | **0.13*** | **0.27** | **0.083** | **0.67** | 0.45 | ●●●●●○ |
| cas0051 | 4n4c | **0.22*** | **0.22*** | **0.18** | **0.071** | **0.56** | **0.52** | ●●●●●● |

| | | | | | | | | |
|---|---|---|---|---|---|---|---|---|
| cas0052 | 2z60 | **0.03*** | -0.026 | **0.095** | 0.049 | **0.69** | **0.55** | ●○●○●● |
| cas0054 | 2jfx | **0.015*** | -0.063 | 0.033 | 0.017 | 0.49 | 0.47 | ●○○○○○ |
| cas0054 | 2jfy | **0.0038*** | -0.079 | **0.067** | 0.011 | 0.48 | 0.46 | ●○●○○○ |
| cas0054 | 2jfz | -0.11 | -0.12 | 0.033 | 0 | 0.4 | 0.42 | ○○○○○○ |
| cas0054 | 4b1f | -0.15 | -0.14 | 0 | 0.0033 | 0.38 | 0.41 | ○○○○○○ |
| cas0056 | 3hmi | **0.015*** | **0.031*** | **0.12** | 0.014 | **0.52** | 0.46 | ●●●○●○ |
| cas0060 | 1boz | -0.068 | **0.039*** | 0.033 | 0.029 | **0.59** | **0.52** | ○●○○●● |
| cas0060 | 1dhf | -0.053 | **0.037*** | **0.05** | 0.03 | 0.44 | 0.48 | ○●●○○○ |
| cas0060 | 1dlr | **0.051*** | **0.061*** | **0.1** | 0.045 | **0.7** | 0.48 | ●●●○●● |
| cas0060 | 1dls | -0.02 | **0.063*** | 0 | 0.042 | **0.65** | **0.52** | ○●○○●● |
| cas0060 | 1drf | -0.033 | **0.066*** | 0.033 | 0.024 | **0.6** | **0.57** | ○●○○●● |
| cas0060 | 1hfp | -0.051 | **0.027*** | **0.067** | 0.028 | **0.58** | **0.52** | ○●●○●● |
| cas0060 | 1hfq | -0.031 | **0.046*** | **0.1** | 0.037 | **0.62** | **0.53** | ○●●○●● |
| cas0060 | 1hfr | **0.0022*** | **0.1*** | 0.033 | 0.039 | **0.67** | **0.58** | ●●○○●● |
| cas0060 | 1kms | -0.031 | **0.092*** | 0.033 | 0.024 | **0.62** | **0.55** | ○●○○●● |
| cas0060 | 1mvs | -0.07 | **0.031*** | 0 | 0.028 | **0.56** | **0.52** | ○●○○●● |
| cas0060 | 1mvt | -0.013 | **0.031*** | 0 | 0.035 | **0.64** | 0.49 | ○●○○●○ |
| cas0060 | 1ohj | -0.054 | **0.065*** | 0 | 0.033 | **0.54** | **0.56** | ○●○○●● |
| cas0060 | 1ohk | -0.017 | **0.08*** | 0 | 0.043 | **0.61** | **0.56** | ○●○○●● |
| cas0060 | 1pd8 | -0.031 | **0.05*** | **0.067** | 0.041 | **0.62** | **0.52** | ○●●○●● |
| cas0060 | 1pd9 | -0.04 | **0.043*** | 0 | 0.03 | **0.61** | **0.52** | ○●○○●● |
| cas0060 | 1s3u | -0.023 | **0.031*** | 0.033 | 0.038 | **0.64** | **0.51** | ○●○○●● |
| cas0060 | 1s3v | -0.034 | **0.077*** | 0.033 | 0.029 | **0.63** | **0.56** | ○●○○●● |
| cas0060 | 1s3w | -0.0054 | **0.055*** | **0.067** | 0.041 | **0.64** | **0.53** | ○●●○●● |
| cas0060 | 1u71 | -0.029 | **0.057*** | **0.067** | 0.039 | **0.62** | **0.56** | ○●●○●● |
| cas0060 | 1u72 | **0.0019*** | **0.024*** | 0 | 0.047 | **0.66** | 0.49 | ●●○○●○ |

| | | | | | | | | |
|---|---|---|---|---|---|---|---|---|
| cas0060 | 2c2s | -0.036 | **0.06*** | 0.017 | 0.04 | 0.45 | 0.49 | ○●○○○○ |
| cas0060 | 2c2t | -0.026 | **0.061*** | **0.05** | **0.051** | 0.46 | 0.48 | ○●●●○○ |
| cas0060 | 2dhf | -0.054 | **0.067*** | **0.05** | 0.037 | 0.45 | 0.49 | ○●●○○○ |
| cas0060 | 2w3a | -0.025 | **0.062*** | 0.017 | 0.044 | 0.45 | 0.49 | ○●○○○○ |
| cas0060 | 2w3m | -0.0046 | **0.089*** | 0.017 | 0.048 | 0.48 | **0.5** | ○●○○○● |
| cas0060 | 3f8y | -0.048 | **0.096*** | 0.033 | **0.059** | **0.59** | **0.55** | ○●○●●● |
| cas0060 | 3f8z | -0.051 | **0.063*** | 0.033 | 0.043 | **0.59** | **0.54** | ○●○○●● |
| cas0060 | 3fs6 | -0.026 | **0.08*** | **0.067** | 0.047 | **0.64** | **0.53** | ○●●○●● |
| cas0060 | 3ghc | -0.0013 | **0.052*** | 0 | **0.055** | **0.66** | 0.49 | ○●○●●○ |
| cas0060 | 3ghw | **0.0078*** | **0.082*** | 0.033 | **0.052** | **0.67** | **0.53** | ●●○●●● |
| cas0060 | 3gi2 | -0.016 | **0.099*** | **0.067** | **0.051** | **0.65** | **0.54** | ○●●●●● |
| cas0060 | 3gyf | -0.011 | **0.11*** | **0.067** | **0.055** | **0.63** | **0.55** | ○●●●●● |
| cas0060 | 3l3r | -0.024 | **0.081*** | 0.033 | 0.048 | **0.66** | **0.53** | ○●○○●● |
| cas0060 | 3n0h | -0.011 | **0.088*** | **0.1** | **0.053** | **0.64** | **0.57** | ○●●●●● |
| cas0060 | 3ntz | -0.022 | **0.067*** | **0.067** | **0.05** | **0.65** | **0.54** | ○●●●●● |
| cas0060 | 3nu0 | -0.0027 | **0.086*** | 0.033 | **0.057** | **0.66** | **0.52** | ○●○●●● |
| cas0060 | 3nxr | -0.067 | **0.082*** | **0.067** | 0.038 | **0.58** | **0.55** | ○●●○●● |
| cas0060 | 3nxt | -0.082 | **0.049*** | **0.067** | 0.033 | **0.59** | **0.53** | ○●●○●● |
| cas0060 | 3nxv | -0.041 | **0.051*** | **0.067** | 0.034 | **0.63** | **0.53** | ○●●○●● |
| cas0060 | 3nxx | -0.08 | **0.042*** | **0.067** | 0.043 | **0.57** | **0.52** | ○●●○●● |
| cas0060 | 3nxy | -0.021 | **0.049*** | **0.067** | 0.031 | **0.62** | **0.52** | ○●●○●● |
| cas0060 | 3nzd | -0.058 | **0.06*** | 0.033 | 0.042 | **0.6** | **0.53** | ○●○○●● |
| cas0060 | 3oaf | -0.017 | **0.097*** | **0.067** | 0.04 | **0.66** | **0.56** | ○●●○●● |
| cas0060 | 3s3v | -0.064 | **0.1*** | **0.067** | 0.039 | **0.6** | **0.57** | ○●●○●● |
| cas0060 | 3s7a | -0.023 | **0.078*** | **0.067** | 0.037 | **0.64** | **0.54** | ○●○●●● |
| cas0060 | 4ddr | -0.019 | **0.037*** | **0.1** | 0.041 | **0.64** | 0.49 | ○○○○●● |

| | | | | | | | | |
|---|---|---|---|---|---|---|---|---|
| cas0060 | 4g95 | -0.079 | **0.06*** | **0.067** | 0.04 | **0.6** | **0.53** | ○●●○●● |
| cas0060 | 4kd7 | -0.026 | **0.077*** | 0.033 | **0.051** | 0.47 | **0.51** | ○●○●○● |
| cas0060 | 4keb | -0.034 | **0.077*** | **0.05** | 0.047 | 0.45 | 0.49 | ○●●○○○ |
| cas0060 | 4m6k | -0.021 | **0.084*** | 0.033 | 0.048 | **0.64** | **0.54** | ○●○○●● |
| cas0060 | 4m6l | -0.06 | **0.084*** | **0.067** | 0.03 | **0.6** | **0.55** | ○●●○●● |
| cas0060 | 4qjc | -0.086 | **0.044*** | **0.067** | 0.036 | **0.57** | **0.52** | ○●●○●● |
| cas0060 | 5hpb | -0.047 | **0.039*** | 0.033 | 0.04 | **0.62** | 0.47 | ○●○○●○ |
| cas0060 | 5hqy | -0.064 | **0.021*** | 0.033 | 0.031 | **0.61** | 0.46 | ○●○○●○ |
| cas0060 | 5hqz | -0.07 | **0.073*** | 0.033 | 0.043 | **0.6** | **0.54** | ○●○○●● |
| cas0060 | 5hsr | -0.014 | **0.038*** | 0.033 | 0.045 | **0.63** | 0.48 | ○●○○●○ |
| cas0060 | 5hsu | -0.046 | **0.072*** | 0.033 | 0.039 | **0.62** | **0.52** | ○●○○●● |
| cas0060 | 5ht4 | -0.024 | **0.057*** | 0.033 | 0.04 | **0.65** | **0.52** | ○●○○●● |
| cas0060 | 5ht5 | -0.041 | **0.071*** | 0.033 | 0.042 | **0.61** | **0.52** | ○●○○●● |
| cas0060 | 5hui | -0.06 | **0.057*** | 0.033 | 0.04 | **0.6** | **0.5** | ○●○○●● |
| cas0060 | 5hvb | -0.022 | **0.052*** | 0.033 | 0.048 | **0.65** | **0.52** | ○●○○●● |
| cas0060 | 5hve | -0.054 | **0.052*** | 0.033 | 0.037 | **0.6** | **0.53** | ○●○○●● |
| cas0061 | 1dre | -0.021 | **0.14*** | 0.027 | **0.071** | **0.56** | **0.63** | ○●○●●● |
| cas0061 | 1ra2 | -0.043 | -0.024 | 0.027 | 0.047 | **0.52** | 0.49 | ○○○○●○ |
| cas0061 | 1ra3 | -0.046 | -0.014 | 0.027 | **0.05** | **0.53** | 0.49 | ○○○●●○ |
| cas0061 | 1ra8 | -0.033 | -0.034 | 0.027 | **0.058** | **0.54** | **0.5** | ○○○●●● |
| cas0061 | 1rb2 | **0.0022*** | **0.026*** | **0.054** | **0.067** | **0.52** | **0.55** | ●●●●●● |
| cas0061 | 1rb3 | **0.0024*** | **0.024*** | 0.027 | **0.053** | 0.38 | 0.48 | ●●○●○○ |
| cas0061 | 1rc4 | -0.035 | -0.061 | 0.027 | 0.046 | **0.56** | 0.46 | ○○○○●○ |
| cas0061 | 1rd7 | -0.019 | -0.0052 | 0.027 | **0.06** | 0.38 | 0.46 | ○○○●○○ |
| cas0061 | 1re7 | -0.023 | -0.0013 | 0.027 | 0.045 | 0.37 | 0.45 | ○○○○○○ |
| cas0061 | 1rg7 | -0.036 | **0.077*** | 0.027 | **0.079** | **0.54** | **0.55** | ○●○●●● |

| | | | | | | | | |
|---|---|---|---|---|---|---|---|---|
| cas0061 | 1rh3 | -0.024 | **0.1*** | 0.027 | **0.078** | **0.58** | **0.6** | ○●○●●● |
| cas0061 | 1rx4 | -0.036 | -0.021 | 0.027 | 0.02 | **0.6** | 0.48 | ○○○○●○ |
| cas0061 | 1rx5 | -0.047 | -0.036 | 0.027 | **0.051** | **0.58** | 0.47 | ○○○●●○ |
| cas0061 | 1rx6 | -0.017 | -0.048 | 0 | 0.036 | **0.63** | 0.46 | ○○○○●○ |
| cas0061 | 1rx7 | -0.024 | -0.022 | 0.027 | 0.041 | **0.61** | 0.47 | ○○○○●○ |
| cas0061 | 3dau | -0.0062 | **0.13*** | 0.027 | **0.068** | **0.58** | **0.58** | ○●○●●● |
| cas0061 | 4ej1 | **0.039*** | **0.023*** | 0.027 | **0.067** | **0.55** | **0.54** | ●●○●●● |
| cas0061 | 4fhb | -0.056 | -0.0025 | 0.027 | 0.043 | **0.5** | **0.56** | ○○○○●● |
| cas0061 | 4i13 | **0.059*** | **0.082*** | **0.054** | **0.074** | **0.58** | **0.6** | ●●●●●● |
| cas0061 | 4i1n | **0.021*** | **0.075*** | 0.027 | **0.056** | **0.57** | **0.61** | ●●○●●● |
| cas0061 | 4kjj | **0.0083*** | **0.064*** | 0.027 | **0.077** | **0.65** | **0.53** | ●●○●●● |
| cas0061 | 4kjl | **0.046*** | **0.1*** | **0.081** | 0.036 | **0.72** | 0.49 | ●●●○●○ |
| cas0061 | 4p3r | **0.00061*** | **0.11*** | **0.054** | **0.088** | **0.59** | **0.56** | ●●●●●● |
| cas0061 | 4qle | -0.01 | **0.048*** | 0.027 | 0.047 | **0.51** | **0.53** | ○●○○●● |
| cas0061 | 4qlg | -0.0042 | **0.011*** | 0.014 | 0.042 | 0.39 | 0.46 | ○●○○○○ |
| cas0061 | 4x5f | -0.061 | -0.084 | 0.014 | 0.034 | 0.38 | 0.4 | ○○○○○○ |
| cas0061 | 4x5g | -0.083 | -0.063 | 0.027 | 0.033 | **0.53** | **0.51** | ○○○○●● |
| cas0061 | 4x5h | -0.034 | -0.032 | 0 | 0.044 | **0.55** | 0.49 | ○○○○●○ |
| cas0061 | 4x5i | -0.041 | -0.027 | 0 | 0.043 | **0.54** | 0.48 | ○○○○●○ |
| cas0061 | 4x5j | -0.042 | -0.05 | 0.027 | 0.04 | **0.55** | 0.44 | ○○○○●○ |
| cas0061 | 5cc9 | -0.029 | **0.027*** | 0 | **0.067** | **0.59** | 0.49 | ○●○●●○ |
| cas0061 | 5ccc | -0.032 | **0.074*** | 0 | **0.067** | **0.54** | **0.56** | ○●○●●● |
| cas0061 | 7dfr | -0.018 | **0.063*** | 0.027 | **0.063** | **0.59** | 0.47 | ○●○●●○ |
| cas0067 | 2psq | **0.037*** | **0.057*** | **0.054** | **0.11** | **0.59** | **0.64** | ●●●●●● |
| cas0067 | 3ri1 | **0.033*** | **0.022*** | **0.071** | **0.085** | **0.58** | **0.61** | ●●●●●● |
| cas0070 | 1ibc | -0.0096 | **0.00064*** | **0.091** | **0.078** | **0.67** | **0.64** | ○●●●●● |

| | | | | | | | | |
|---|---|---|---|---|---|---|---|---|
| cas0070 | 1ice | -0.028 | -0.087 | **0.091** | **0.054** | **0.63** | **0.54** | ○○●●●● |
| cas0070 | 1rwm | **0.051*** | **0.011*** | **0.091** | **0.1** | **0.73** | **0.65** | ●●●●●● |
| cas0070 | 1rwn | **0.051*** | -0.019 | **0.091** | **0.079** | **0.73** | **0.61** | ●○●●●● |
| cas0070 | 1rwo | **0.046*** | -0.043 | **0.091** | **0.066** | **0.73** | **0.6** | ●○●●●● |
| cas0070 | 1rwv | **0.05*** | **0.016*** | **0.091** | **0.094** | **0.73** | **0.65** | ●●●●●● |
| cas0070 | 2h4w | **0.01*** | -0.0088 | 0 | **0.081** | **0.69** | **0.63** | ●○○●●● |
| cas0070 | 2h4y | **0.061*** | -0.015 | **0.091** | **0.068** | **0.74** | **0.63** | ●○●●●● |
| cas0070 | 2h51 | -0.028 | -0.032 | 0 | **0.075** | **0.65** | **0.61** | ○○○●●● |
| cas0070 | 2h54 | **0.057*** | -0.018 | **0.091** | **0.089** | **0.74** | **0.62** | ●○●●●● |
| cas0070 | 2hbq | **0.044*** | **0.0079*** | **0.14** | **0.087** | **0.72** | **0.64** | ●●●●●● |
| cas0070 | 2hbr | -0.031 | -0.012 | 0 | **0.08** | **0.64** | **0.64** | ○○○●●● |
| cas0070 | 2hby | **0.018*** | **0.0086*** | 0.045 | **0.076** | **0.69** | **0.64** | ●●○●●● |
| cas0070 | 2hbz | **0.00072*** | -0.021 | 0.045 | **0.064** | **0.67** | **0.62** | ●○○●●● |
| cas0070 | 3d6f | **0.049*** | -0.011 | **0.14** | **0.09** | **0.72** | **0.63** | ●○●●●● |
| cas0070 | 3d6h | **0.077*** | **0.012*** | **0.091** | **0.082** | **0.75** | **0.67** | ●●●●●● |
| cas0070 | 3d6m | **0.048*** | -0.02 | **0.091** | **0.084** | **0.74** | **0.62** | ●○●●●● |
| cas0071 | 1i4o | **0.17*** | **0.073*** | **0.2** | **0.075** | **0.82** | **0.7** | ●●●●●● |
| cas0074 | 1ibv | **0.48*** | **0.087*** | **0.67** | **0.23** | **0.85** | **0.69** | ●●●●●● |
| cas0074 | 1ibw | **0.34*** | **0.13*** | **0.67** | **0.3** | **0.73** | **0.7** | ●●●●●● |
| cas0074 | 1pya | **0.47*** | **0.18*** | **0.33** | **0.16** | **0.85** | **0.76** | ●●●●●● |
| cas0079 | 1bzc | **0.26*** | **0.17*** | **0.11** | **0.065** | **0.79** | **0.6** | ●●●●●● |
| cas0079 | 1bzj | **0.31*** | **0.2*** | **0.33** | **0.11** | **0.83** | **0.66** | ●●●●●● |
| cas0079 | 1c83 | **0.31*** | **0.19*** | **0.33** | **0.12** | **0.83** | **0.65** | ●●●●●● |
| cas0079 | 1c84 | **0.32*** | **0.16*** | **0.17** | **0.11** | **0.85** | **0.62** | ●●●●●● |
| cas0079 | 1c85 | **0.31*** | **0.14*** | **0.22** | **0.091** | **0.84** | **0.61** | ●●●●●● |
| cas0079 | 1c86 | **0.33*** | **0.2*** | **0.33** | **0.097** | **0.85** | **0.67** | ●●●●●● |

| | | | | | | | | |
|---|---|---|---|---|---|---|---|---|
| cas0079 | 1c87 | **0.33*** | **0.19*** | **0.33** | **0.11** | **0.85** | **0.64** | ●●●●●● |
| cas0079 | 1c88 | **0.35*** | **0.18*** | **0.28** | **0.11** | **0.86** | **0.64** | ●●●●●● |
| cas0079 | 1ecv | **0.32*** | **0.21*** | **0.28** | **0.081** | **0.83** | **0.65** | ●●●●●● |
| cas0079 | 1g1g | **0.3*** | **0.17*** | **0.33** | **0.083** | **0.81** | **0.59** | ●●●●●● |
| cas0079 | 1g7g | **0.32*** | **0.17*** | **0.33** | **0.1** | **0.83** | **0.61** | ●●●●●● |
| cas0079 | 1gfy | **0.33*** | **0.18*** | **0.39** | **0.11** | **0.84** | **0.64** | ●●●●●● |
| cas0079 | 1kak | **0.25*** | **0.15*** | **0.22** | **0.087** | **0.8** | **0.61** | ●●●●●● |
| cas0079 | 1kav | **0.26*** | **0.16*** | **0.22** | **0.071** | **0.81** | **0.61** | ●●●●●● |
| cas0079 | 1l8g | **0.34*** | **0.18*** | **0.33** | **0.11** | **0.85** | **0.66** | ●●●●●● |
| cas0079 | 1nwe | **0.23*** | **0.2*** | **0.17** | **0.083** | **0.77** | **0.65** | ●●●●●● |
| cas0079 | 1ptt | **0.32*** | **0.18*** | **0.28** | **0.11** | **0.84** | **0.63** | ●●●●●● |
| cas0079 | 1ptu | **0.31*** | **0.16*** | **0.33** | **0.14** | **0.75** | **0.66** | ●●●●●● |
| cas0079 | 1ptv | **0.3*** | **0.21*** | **0.28** | **0.11** | **0.82** | **0.65** | ●●●●●● |
| cas0079 | 1q1m | **0.31*** | **0.18*** | **0.39** | **0.1** | **0.84** | **0.66** | ●●●●●● |
| cas0079 | 1xbo | **0.32*** | **0.18*** | **0.33** | **0.12** | **0.85** | **0.65** | ●●●●●● |
| cas0079 | 2azr | **0.32*** | **0.19*** | **0.39** | **0.096** | **0.84** | **0.64** | ●●●●●● |
| cas0079 | 2b07 | **0.32*** | **0.19*** | **0.39** | **0.097** | **0.85** | **0.64** | ●●●●●● |
| cas0079 | 2bge | **0.3*** | **0.16*** | **0.39** | **0.1** | **0.83** | **0.62** | ●●●●●● |
| cas0079 | 2cm7 | **0.31*** | **0.16*** | **0.33** | **0.086** | **0.82** | **0.6** | ●●●●●● |
| cas0079 | 2cma | **0.27*** | **0.15*** | **0.22** | **0.084** | **0.8** | **0.6** | ●●●●●● |
| cas0079 | 2h4g | **0.32*** | **0.19*** | **0.44** | **0.11** | **0.84** | **0.66** | ●●●●●● |
| cas0079 | 2h4k | **0.32*** | **0.18*** | **0.44** | **0.11** | **0.82** | **0.63** | ●●●●●● |
| cas0079 | 2hb1 | **0.3*** | **0.19*** | **0.39** | **0.1** | **0.84** | **0.64** | ●●●●●● |
| cas0079 | 2nt7 | **0.3*** | **0.19*** | **0.39** | **0.089** | **0.84** | **0.65** | ●●●●●● |
| cas0079 | 2nta | **0.32*** | **0.17*** | **0.44** | **0.1** | **0.84** | **0.64** | ●●●●●● |
| cas0079 | 2qbp | **0.31*** | **0.23*** | **0.33** | **0.11** | **0.84** | **0.67** | ●●●●●● |

| | | | | | | | | |
|---|---|---|---|---|---|---|---|---|
| cas0079 | 2qbq | **0.31*** | **0.19*** | **0.33** | **0.091** | **0.83** | **0.64** | ●●●●●● |
| cas0079 | 2qbr | **0.34*** | **0.18*** | **0.44** | **0.099** | **0.85** | **0.65** | ●●●●●● |
| cas0079 | 2qbs | **0.31*** | **0.19*** | **0.39** | **0.095** | **0.83** | **0.65** | ●●●●●● |
| cas0079 | 2veu | **0.3*** | **0.14*** | **0.28** | **0.085** | **0.82** | **0.6** | ●●●●●● |
| cas0079 | 2vey | **0.28*** | **0.15*** | **0.33** | **0.084** | **0.8** | **0.6** | ●●●●●● |
| cas0079 | 2zn7 | **0.34*** | **0.18*** | **0.39** | **0.1** | **0.84** | **0.62** | ●●●●●● |
| cas0079 | 4i8n | **0.33*** | **0.19*** | **0.39** | **0.1** | **0.85** | **0.64** | ●●●●●● |
| cas0080 | 1i00 | -0.075 | -0.052 | 0 | **0.086** | 0.36 | 0.47 | ○○○●○○ |
| cas0080 | 1juj | -0.15 | -0.022 | 0 | 0.018 | 0.16 | 0.43 | ○○○○○○ |
| cas0080 | 3ob7 | -0.043 | -0.1 | **0.075** | **0.072** | 0.18 | 0.34 | ○○●●○○ |
| cas0080 | 5hs3 | -0.15 | -0.095 | 0.021 | 0.038 | 0.081 | 0.29 | ○○○○○○ |
| cas0080 | 5x5q | -0.1 | -0.039 | **0.083** | **0.062** | 0.12 | 0.4 | ○○●●○○ |
| cas0080 | 5x67 | -0.022 | -0.022 | **0.062** | **0.079** | 0.42 | 0.47 | ○○●●○○ |
| cas0085 | 1fuo | -0.072 | -0.069 | 0 | 0.045 | 0.28 | **0.51** | ○○○○○● |
| cas0085 | 1fup | -0.079 | -0.099 | 0 | **0.05** | 0.28 | 0.48 | ○○○●○○ |
| cas0085 | 1fuq | -0.022 | -0.1 | 0 | **0.06** | 0.31 | 0.47 | ○○○●○○ |
| cas0085 | 1kq7 | -0.073 | -0.15 | 0 | **0.062** | 0.28 | 0.44 | ○○○●○○ |
| cas0086 | 1gos | **0.17*** | -0.0072 | 0.029 | 0.013 | 0.4 | 0.48 | ●●●●●● |
| cas0086 | 1oj9 | **0.13*** | -0.024 | 0 | 0 | 0.38 | 0.47 | ●○○○○○ |
| cas0086 | 1oja | **0.11*** | -0.041 | 0 | 0.0026 | 0.35 | 0.45 | ●○○○○○ |
| cas0086 | 1ojc | **0.11*** | -0.03 | 0 | 0 | 0.36 | 0.46 | ●○○○○○ |
| cas0086 | 1s2q | **0.15*** | -2.70E-05 | 0 | 0.0025 | 0.39 | 0.49 | ●○○○○○ |
| cas0086 | 1s2y | **0.13*** | -0.021 | 0 | 0.0049 | 0.37 | 0.47 | ●○○○○○ |
| cas0086 | 1s3b | **0.15*** | -0.0078 | 0 | 0.0051 | 0.38 | 0.48 | ●○○○○○ |
| cas0086 | 1s3e | **0.18*** | **0.00018*** | 0 | 0.0025 | 0.41 | 0.48 | ●●○○○○ |
| cas0086 | 2bk3 | **0.13*** | -0.026 | 0 | 0 | 0.38 | 0.46 | ●○○○○○ |

| | | | | | | | | |
|---|---|---|---|---|---|---|---|---|
| cas0086 | 2bk4 | **0.2*** | **0.009*** | 0 | 0 | 0.42 | **0.5** | ●●○○○○ |
| cas0086 | 2bk5 | **0.17*** | -0.018 | 0 | 0.0025 | 0.39 | 0.47 | ●○○○○○ |
| cas0086 | 2byb | **0.15*** | -0.0091 | 0 | 0.0077 | 0.38 | 0.48 | ●○○○○○ |
| cas0086 | 2c64 | **0.13*** | -0.019 | 0 | 0.0054 | 0.37 | 0.47 | ●○○○○○ |
| cas0086 | 2c65 | **0.14*** | -0.026 | 0 | 0.0082 | 0.38 | 0.46 | ●○○○○○ |
| cas0086 | 2c66 | **0.14*** | -0.012 | 0 | 0.0052 | 0.38 | 0.48 | ●○○○○○ |
| cas0086 | 2c67 | **0.15*** | -0.01 | 0 | 0.0025 | 0.38 | 0.47 | ●○○○○○ |
| cas0086 | 2c70 | **0.17*** | -0.014 | 0 | 0.0052 | 0.39 | 0.47 | ●○○○○○ |
| cas0086 | 2c72 | **0.16*** | -0.011 | 0 | 0.0049 | 0.38 | 0.47 | ●○○○○○ |
| cas0086 | 2c73 | **0.14*** | **0.011*** | 0 | 0.0051 | 0.38 | **0.5** | ●●○○○○ |
| cas0086 | 2c75 | **0.21*** | **0.0079*** | 0 | 0.0025 | 0.42 | **0.5** | ●●○○○○ |
| cas0086 | 2c76 | **0.16*** | -0.021 | 0 | 0.0049 | 0.39 | 0.48 | ●○○○○○ |
| cas0086 | 2v5z | **0.12*** | -0.019 | 0 | 0.0028 | 0.36 | 0.47 | ●○○○○○ |
| cas0086 | 2v60 | **0.14*** | -0.014 | 0 | 0.0053 | 0.37 | 0.47 | ●○○○○○ |
| cas0086 | 2v61 | **0.13*** | -0.016 | 0 | 0.0026 | 0.37 | 0.47 | ●○○○○○ |
| cas0086 | 2vrl | **0.14*** | -0.011 | **0.059** | 0 | 0.38 | 0.47 | ●○●○○○ |
| cas0086 | 2vrm | **0.17*** | **0.012*** | **0.059** | 0.0026 | 0.39 | 0.49 | ●●●○○○ |
| cas0086 | 2vz2 | **0.17*** | **0.011*** | 0 | 0.0026 | 0.4 | 0.48 | ●●○○○○ |
| cas0086 | 2xcg | **0.14*** | -0.017 | 0.029 | 0 | 0.38 | 0.48 | ●○○○○○ |
| cas0086 | 2xfn | **0.15*** | -0.013 | 0 | 0 | 0.38 | 0.47 | ●○○○○○ |
| cas0086 | 2xfo | **0.1*** | -0.012 | 0 | 0 | 0.36 | 0.48 | ●○○○○○ |
| cas0086 | 2xfp | **0.16*** | -0.015 | 0 | 0 | 0.39 | 0.47 | ●○○○○○ |
| cas0086 | 2xfq | **0.18*** | **0.0045*** | 0 | 0 | 0.4 | 0.48 | ●●○○○○ |
| cas0086 | 2xfu | **0.087*** | -0.049 | **0.059** | 0.0056 | 0.34 | 0.44 | ●○●○○○ |
| cas0086 | 3po7 | **0.15*** | -0.022 | 0 | 0 | 0.38 | 0.47 | ●○○○○○ |
| cas0086 | 3zyx | **0.049*** | -0.037 | 0 | 0.0058 | 0.31 | 0.45 | ●○○○○○ |

| | | | | | | | | |
|---|---|---|---|---|---|---|---|---|
| cas0086 | 4a79 | **0.07*** | -0.047 | 0 | 0.0058 | 0.33 | 0.44 | ●○○○○○ |
| cas0086 | 4a7a | **0.13*** | -0.016 | 0 | 0.0083 | 0.37 | 0.48 | ●○○○○○ |
| cas0086 | 4crt | **0.12*** | -0.029 | 0 | 0.0082 | 0.37 | 0.46 | ●○○○○○ |
| cas0086 | 5mrl | **0.18*** | **0.0074*** | 0 | 0.0025 | 0.41 | 0.49 | ●●○○○○ |
| cas0091 | 1fin | **0.1*** | **0.029*** | 0.019 | 0.049 | 0.42 | **0.54** | ●●○○○● |
| cas0091 | 2b54 | **0.083*** | **0.0001*** | **0.12** | 0.027 | **0.64** | **0.51** | ●●●○●● |
| cas0091 | 2cch | **0.2*** | **0.03*** | **0.077** | **0.059** | **0.51** | **0.55** | ●●●●●● |
| cas0091 | 4ez7 | **0.15*** | **0.039*** | 0.038 | 0.026 | **0.7** | **0.53** | ●●○○●● |
